# Unraveling cellular complexity with unlimited multiplexed super-resolution imaging

**DOI:** 10.1101/2023.05.17.541061

**Authors:** Florian Schueder, Felix Rivera-Molina, Maohan Su, Phylicia Kidd, James E. Rothman, Derek Toomre, Joerg Bewersdorf

**Affiliations:** Department of Cell Biology, Yale School of Medicine, New Haven, CT, USA; Department of Microbial Pathogenesis, Yale School of Medicine, New Haven, CT, USA; Nanobiology Institute, Yale University, West Haven, CT, USA; Department of Biomedical Engineering, Yale University, New Haven, CT, USA; Kavli Institute for Neuroscience, Yale School of Medicine, New Haven, CT, USA; Department of Physics, Yale University, New Haven, CT, USA

## Abstract

Mapping the intricate spatial relationships between the many different molecules inside a cell is essential to understanding cellular functions in all their complexity. Super-resolution fluorescence microscopy offers the required spatial resolution but struggles to reveal more than four different targets simultaneously. Exchanging labels in subsequent imaging rounds for multiplexed imaging extends this number but is limited by its low throughput. Here we present a novel imaging method for rapid multiplexed super-resolution microscopy of a nearly unlimited number of molecular targets by leveraging fluorogenic labeling in conjunction with Transient Adapter-mediated switching for high-throughput DNA-PAINT (FLASH-PAINT). We demonstrate the cell biological versatility of FLASH-PAINT in mammalian cells in four applications: i) mapping nine proteins in a single mammalian cell, ii) elucidating the functional organization of primary cilia by nine-target imaging, iii) revealing the changes in proximity of twelve different targets in unperturbed and dissociated Golgi stacks and iv) investigating inter-organelle contacts at 3D super-resolution.

## Highlights

– **FLASH-PAINT enables spectrally unlimited multiplexed super-resolution imaging**
– **Transient Adapters and Erasers allow for fast, efficient and gentle label exchange**
– **Multiplexed super-resolution imaging reveals complex cilia and Golgi organization**
– **3D FLASH-PAINT allows for quantification of inter-organelle site numbers and areas**

## Introduction

Understanding cellular function is intimately tied to our ability to visualize how organelles and the molecules constituting them respond to diverse physiological and disease states. Meaningful, information-rich visualization is, however, a challenge as it depends on both of our abilities to identify molecules, in particular proteins, and their many interaction partners, and resolve their spatial organization. Fluorescence light microscopy has long been key here, revealing specific proteins at hundreds of nanometers resolution, or, with the advent of optical super-resolution microscopy^1–3^ at tens of nanometers or even sub-ten nanometer^4^ resolution. Among the different super-resolution microscopy modalities, single-molecule localization microscopy (SMLM) is a preferred choice for cell biological investigations due to its high 3D resolution (usually ∼20-70 nm), sensitivity (single molecules), and relatively low instrumentational requirements. In SMLM, single molecules spontaneously switch between ‘ON’ (bright) and ‘OFF’ (dark) states and super-resolved images are built up by computationally localizing individual ON molecules over thousands of camera frames^5^. In contrast to SMLM techniques such as (F)PALM^6, 7^ and (d)STORM^8, 9^ that rely on photophysical switching between bright and dark fluorescent states, DNA-PAINT^10^ utilizes the transient reversible binding of fluorescently tagged short oligonucleotide strands, called ‘Imagers’ (or ‘Imager probes’), to complementary ‘docking strands’ that are linked to targets of interest (e.g. proteins usually tagged via antibodies). In conventional DNA-PAINT there is no true dark fluorescent state, rather the ‘OFF’ state relies on unbound Imagers being blurred to a homogeneous background by their rapid diffusion, thereby preventing their localization; only when an Imager binds to the docking strand, the transiently immobile Imager is observed as a discrete fluorescent spot (‘ON’) that can be localized. Freed of the constraints of photophysical switching, dyes and buffers in DNA-PAINT can be selected for maximum brightness. Additionally, as there is a large reserve pool of Imagers in solution, even as a bound Imager probe bleaches it can be replaced. This allows for higher densities of localization events in the final image, an otherwise limiting factor, especially when imaging volumes as thick as a cell^11^. The combination of these advantages allows for <5 nm resolution^12, 13^.

While DNA-PAINT and other super-resolution techniques feature an impressive resolution improvement of a factor of ten or more over conventional fluorescence microscopy, its impact on biomedical research has been limited by a lack of multicolor imaging techniques which are instrumental to decode the intricate organization of the cell at the molecular level. The mammalian Golgi complex, for example, is organized in stacks of multiple cisternae arranged cis-to-trans. These stacks are usually connected laterally forming a highly convoluted ‘ribbon’^14^. The complex role and structure of the Golgi and its interactions with the trans-Golgi network (TGN), endoplasmic reticulum (ER) exit sites (ERES), the ER Golgi Intermediate Compartment (ERGIC) and many other organelles is mediated by more than one thousand^15^ different proteins which interact in a selective, well-orchestrated manner as governed by their specific spatial distributions. While in our mind’s eye there is a prototypical “textbook” Golgi, the Golgi ribbon in reality varies dramatically in shape and orientation from cell to cell. This variability makes it impossible to combine individual, independently recorded super-resolution images of different subsets of two or three different proteins into a comprehensive ten or more color image that would cover more than just a small facet of the Golgi’s role in cell biology.

Multicolor SMLM has traditionally been constrained by the limited availability of bright, spectrally distinguishable probes. As a result, two-color imaging has been the standard in SMLM, with three or four colors being the exception^16, 17^. Multiplexing approaches in which different labels are imaged sequentially, offer an avenue to overcome this limitation^18^ and have, for example, been demonstrated to extend diffraction-limited multicolor fluorescence imaging up to ∼100 labels^19^. In super-resolution microscopy, multiplexing has been realized by the DNA-PAINT variant Exchange-PAINT^20^. Here, different targets are labeled with orthogonal ssDNA docking strands and then imaged sequentially using different Imager probes. However, the Imager probes used so far feature slow binding kinetics^21^ which result in data acquisition of an hour or more per color channel. Adding time for washing between sequential imaging cycles, total data acquisition times typically accumulate to several days for a single cell.

The recent development of speed-optimized^22, 23^ and fluorogenic^24^ Imager probes which allow up ∼100-fold faster imaging in DNA-PAINT, are at a first glance a solution to this severe limitation of throughput. However, due to DNA sequence design constraints of these specialized probes only six speed-optimized probes^23^ and two fluorogenic probes^24^ have so far been found. This limits the prospect of fast Exchange-PAINT to only a hand full of targets. Additionally, requiring a specific Imager probe for each target does not scale well to tens, hundreds or even thousands of probes since dye-conjugated oligonucleotides are expensive and probe exchange by extensive washing after each imaging cycle is time consuming and in accumulation damages the sample.

Here we introduce fluorogenic labeling in conjunction with Transient Adapter-mediated switching for high-throughput DNA-PAINT (FLASH-PAINT), a method that allows for rapid, essentially unlimited multiplexing in super-resolution imaging. Using orthogonal ssDNA-based adapters that direct any Imager probe (e.g. a speed-optimized^22, 23^ or fluorogenic^24^ one) to a specific target selected from a complementary set of docking strands (**Figure 1a**) eliminates the color-limitation of super-resolution microscopy. Key to the success of FLASH-PAINT is that the adapters bind only *transiently* to the docking strands. This allows for fast, efficient, and gentle exchange of adapters between imaging cycles. Furthermore, it enables the introduction of ‘Erasers’, oligonucleotides that are complementary to individual Transient Adapters. Hybridized to any Transient Adapter of choice, these Erasers neutralize it in a highly efficient manner and thereby eliminate the need for any washing steps.

**Figure 1.**
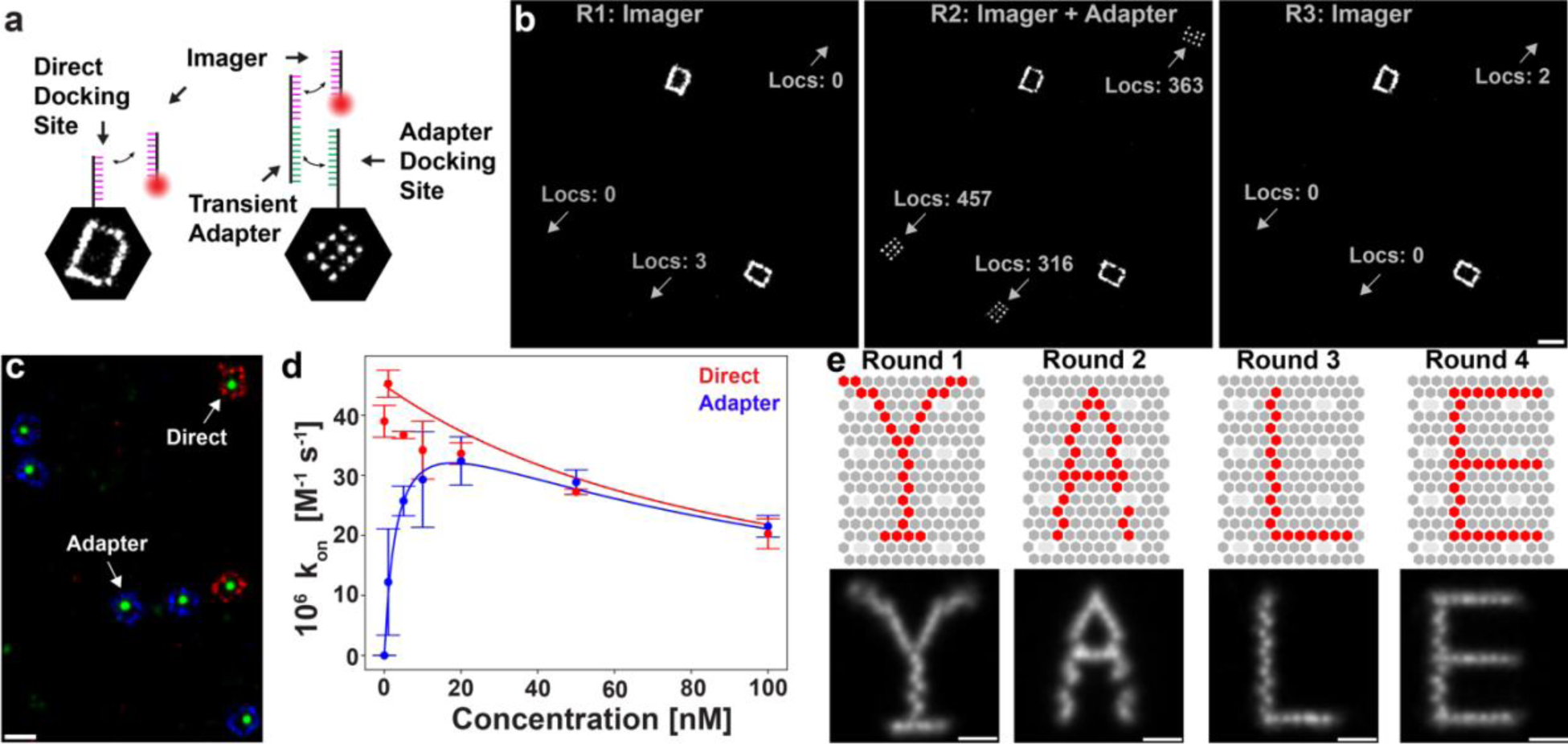
Proof of concept of FLASH-PAINT. **(a)** Schematic representation of ‘classical’ DNA-PAINT using direct binding of Imagers to docking sites and FLASH-PAINT using Transient Adapters. **(b)** Proof of concept with DNA origami nanostructures. Frame pattern DNA origamis are sampled via direct DNA-PAINT, 20-nm grid DNA origamis via Transient Adapters (FLASH-PAINT). **(c)** DNA origami experiment to compare association rates for direct binding and binding via Transient Adapters. (**d**) Measured (data points) and calculated (curves) association rates for direct and adapter-mediated binding for different Transient Adapter concentrations. **(e)** 4-plex imaging of 5-nm grid DNA origamis featuring binding sites arranged as letters (‘Y’, ‘A’, ‘L’, ‘E’). The data was acquired in 25 min per round and ∼100 min total imaging time. 121 to 246 super-resolution images of individual letters were averaged to generate the displayed letters. A representative field of view and individual DNA origami letters are shown in **SI** Figures 10-14. Scale bars: (**b, c**) 100 nm; (c) 20 nm.

We demonstrate the broad utility of FLASH-PAINT by mapping spatial distributions of nine different proteins across a U-2 OS cell and revealing the complex spatial arrangement of nine proteins on individual primary cilia and twelve Golgi-related proteins in single cells. Additionally, we characterize the number and size of contacts between the ER, mitochondria, lysosomes, and the Golgi complex at 3D super-resolution.

## Results

### Adapter design

Minimal crosstalk between targets is a key requirement for successful multiplexed imaging. Multiplexing approaches therefore traditionally have put emphasis on efficiently erasing previous rounds of fluorescent labels (e.g., by photobleaching, UV-cleaving or chemical stripping) before imaging the next round of labels^25–28^. This difficult process of removing labels stands in stark contrast to the close to 100% dissociation efficiency of Imagers from docking sites in DNA-PAINT which is facilitated simply by the transient nature of Imager-docking site association (∼1 s). We hypothesized that this same principle of transient binding can be applied to adapters that bind only transiently to their target. The conceptional challenge with this approach is that such an adapter will inevitably be bound to the target only for a fraction of the time and thereby reduce the overall binding frequency of Imager probes to the docking sites as compared to the conventional adapter-less DNA-PAINT approach. Importantly, however, the Transient Adapter itself is not fluorescent and therefore can be used at concentrations orders of magnitude higher (e.g., *c_TA_* = 10 nM – 100 nM) than an Imager strand in a conventional DNA-PAINT experiment: for example, at a 50-nM Transient Adapter concentration, an average binding time of 100 s, and an association rate of 2 x 10^6^ M^-1^s^-1^, ∼91% of docking sites are occupied by an adapter (see **Supplemental Information).**

We designed a set of Transient Adapters, where each adapter consisted of two binding motifs, one to an Imager probe and the other one to a docking sequence, separated by a short 2-nucleotide (nt) spacer. As Imager probe motifs, we selected three previously published sequences: a conventional DNA-PAINT Imager, a speed-optimized Imager and a fluorogenic Imager (**Table S1**). To realize the targeted dissociation rate of the order of 0.01 s^-1^, we designed 12 orthogonal 10-nt motifs (**Table S2, S3**) with a GC content of 40% - 50%.

### Transient Adapters are highly specific and bind efficiently and reversibly

For an initial proof of concept of FLASH-PAINT, we used DNA origami nanostructures. To directly compare adapter-mediated binding with direct binding, we imaged a mixture of two different DNA origami species with a SMLM instrument. One species featured binding sites for the Imager probe arranged in a rectangular frame, the other species featured adapter docking sites arranged in a 3×4 grid with a 20-nm spacing (**Figure 1a, b**). In the first round of imaging, we introduced only the Imager strand. As expected, we could only observe the first species since the Imager should not bind to the adapter docking site on the second origami species. In the second round, we introduced the adapter along with the Imager and consequently both DNA origami species were visible. For the third round of imaging, to test the adapter dissociation efficiency, we washed out the mix of adapter and Imager strands, and then reintroduced the Imager only. The resulting image resembled the first image, confirming excellent dissociation efficiency. Counting how many Imager probe binding events were registered in the three images, confirmed that unspecific binding of the Imager probe to the adapter docking site lies below 1% (**Figure 1b**; 0-3 vs. 316-457 events) and adapter dissociation is more than 99% efficient (0-2 events after washing).

Next, we designed an experiment to compare the association rate of adapter-mediated binding to the association rate of direct binding (**Figure 1c**). We again mixed two different DNA origami species, one species featuring a single docking site for adapter-mediated binding, the other one having a direct binding site. To distinguish the two DNA origamis from each other, we used orthogonal docking sites arranged in a rectangle framing the single docking site and imaged these frames sequentially in two additional rounds (**Figures 1d, S1 & S2**). We then measured the association rates of Imagers binding to the two species of DNA origamis for a constant Imager concentration (10 nM) but different adapter concentrations (**Figures 1d & S3**) using a speed Imager^23^. In the low adapter-concentration regime, the association rate of Imagers binding to DNA origamis via adapters increases when raising the adapter concentration. This can be explained by the increase in occupancy of the docking sites by adapter strands. Beyond ∼20 nM adapter strand concentration, the association rate decreases, however. A similar decrease can simultaneously be observed in the association rate of the Imagers binding directly to the second DNA origami species and is consistent with a decrease of the available Imager concentration. This can be explained by high concentrations of adapter strands in solution competing for Imagers and thereby depleting the pool of Imagers available to bind to DNA origamis. An analytical description of the direct and adapter-mediated association rates is provided in the supplementary information and matches the experimental data points very well (curve vs. data points in **Figure 1d**). Importantly, the adapter-mediated association rate of Imagers binding to DNA origamis reached with ∼70% of the association rate for direct binding (*c_TA_* = 0 nM) a level comparable to DNA-PAINT relying on direct, adapter-less binding of Imagers. This shows that the introduction of Transient Adapters is possible without substantially compromising the association of Imagers to the targets.

Next, we measured the association and dissociation rates for 36 designed adapters (**Table S4**), 12 each for speed (adapter concentration at 20 nM) (**Figures S4 & S5**), classical (**Figures S6 & S7**) and fluorogenic (**Figures S8 & S9**) Imagers. In all measured cases we found that the association rates of the adapter-mediated binding were in a range similar to the direct binding case. This confirmed that transient adapters can generally be used without substantially compromising the association of Imagers to the targets.

### Multiplexed quantitative super-resolution microscopy at high resolution

To test multiplexed imaging via our Transient Adapters we designed DNA origami structures with four different orthogonal docking sites arranged to the shape of the letter’s ‘Y’, ‘A’, ‘L’ and ‘E’ (**Figures 1e & S2**). The localization precision for all four rounds of imaging was ∼2 nm, which allowed us to clearly differentiate between neighboring binding sites of the letter patterns (**Figures 1e & S10 - S14**) and demonstrates that the use of Transient Adapters does not compromise the resolution. Using the speed-optimized Imagers, each round of imaging took ∼25 min, resulting in a total imaging time of ∼100 min.

Another unique feature of the Transient Adapters is the ability of imaging the same target of interest with different Imagers. This allowed us to compare the imaging performance of different Imagers using the same sample and imaging conditions. The outer membrane protein Tom20 in COS-7 cells was immunolabeled with antibodies featuring a ssDNA docking site and imaged under epi-illumination using speed, fluorogenic and classical Imagers via adapters (**Figure S15**). While the signal-to-noise ratio (SNR) of the fluorogenic Imager (SNR ∼30) and the speed Imager (SNR ∼40) was sufficiently high to easily distinguish bound Imagers from diffuse background, the low SNR in the classical Imager case (SNR ∼8) prevented artifact-free localization of targets in the thick sample. This demonstrates the clear advantage of the speed and fluorogenic Imager probes over the classical DNA-PAINT Imagers.

### Erasers allow for rapid and efficient switching between Adapters without washing

In classical Exchange-PAINT, the switch between targets is achieved by thoroughly washing out one Imager and subsequently introducing the next Imager, but this is time-consuming (typically ∼10 min). We reasoned that the washing step could be eliminated in FLASH-PAINT by introducing an Eraser strand (**Figure 2a** and **Table S5**) that is complementary to the Transient Adapter strand from the previous imaging round. The higher-affinity Eraser binds (effectively permanently) to the Transient Adapter and neutralizes it by preventing it from binding to the corresponding docking site.

**Figure 2.**
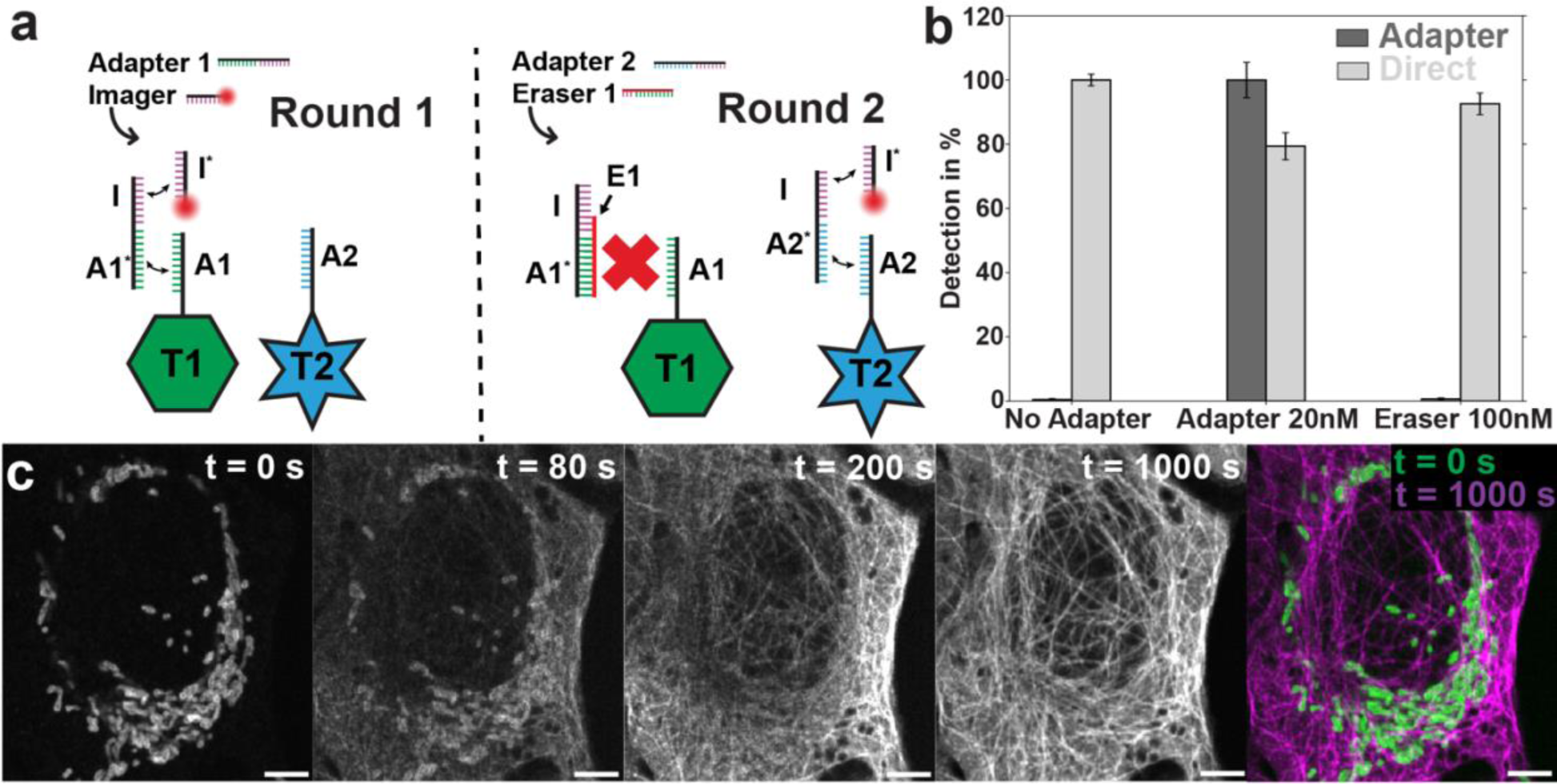
Molecular target switching via a Transient Adapter-Eraser combination. **(a)** Schematic depiction of molecular target switching. Eraser E1 neutralizes in Round 2 Transient Adapter 1 while newly added Transient Adapter 2 directs the Imager probe to a new target. (**b**) Quantification of switching efficiency using DNA origami (at the example of adapter sequence A19). In the first round of imaging only the Imager, not the Transient Adapter, is introduced. In the second round, both Transient Adapter (20 nM) and Imager are applied. Finally, the solution is replaced with an Eraser (100 nM) and an Imager. After a 3-min incubation period, the third round of imaging is carried out. (**c**) Time course of switching the labeled molecular target from Tom20 on mitochondria to α-tubulin (microtubules) in a U-2 OS cell. The sample is in a medium containing Transient Adapter 1 to visualize mitochondria and an Imager pre acquisition. At the start of the acquisition Eraser 1 (to erase signal from mitochondria) and Transient Adapter 2 are added to the medium. Scale bars: 5 µm.

We characterized the erasing efficiency for all twelve Transient Adapter sequences using DNA origami structures and found it to be greater than 98% in all cases (**Figures 2b**, **S16**). As a test in a biological sample, we monitored the redirection of an Imager probe from a docking site on mitochondria (immunolabeling of the mitochondrial outer membrane protein Tom20) to a microtubule docking site (α-tubulin immunolabeling) by simultaneously introducing both an Eraser for the Transient Adapter to the mitochondria docking site and a new Transient Adapter for the microtubule docking site (**Figure 2c** and **Movie S1**). Even without any flow or active perfusion, the mitochondrial signal quickly disappeared (τ_1/2_ ≈ 60 s) as the microtubule signal appeared (τ_1/2_ ≈ 200 s) (**Figure S17a**). Comparing the first with the last frame of the movie shows both the excellent orthogonality and erasing efficiency of the Transient Adapter-mediated labeling as only the mitochondria and only the microtubules are visible at the beginning and end of the movie, respectively. Similarly efficient switching between two targets could be observed in the much denser environment of the cell nucleus by switching the Imager signal from the nucleolus protein NPM1 to Lamin B1 at the nuclear lamina (**Figure S17b** and **Movie S2)**.

To evaluate the non-specific binding of the Transient Adapters (and Imagers) for classical, speed and fluorogenic DNA-PAINT imaging in a cellular imaging context we imaged anti-Tom20 immunolabeled cells with matching and non-matching Transient Adapter-Imager combinations (**Figures S18-S20**). We found that in all three cases the non-specific binding was negligible (<2%) compared to the specific binding, even if eleven non-matching adapters were added.

### FLASH-PAINT enables spectrally unlimited multiplexed super-resolution microscopy in cells

To test FLASH-PAINT’s capability for fast, efficient, spectrally unlimited multiplexed super-resolution microscopy, we imaged nine immunolabeled targets in a U2OS cell including three Golgi proteins (GM130, GRASP55, GRASP65), three mitochondria-associated targets (OMP25, HADHA, dsDNA), two nucleolus-localized targets (NPM1, RPA40) and the nuclear envelope (Lamin-B1) (**Figures 3 & S21**). We imaged one target each in nine subsequent rounds followed by imaging all targets together in a tenth round for spatial alignment. The imaging experiment was completed in only ∼3 hours, which included the time to switch between targets. We achieved an average localization precision of ∼11.2 nm. To broadly demonstrate the cell biological utility of FLASH-PAINT, we tested our new method in three other applications.

**Figure 3.**
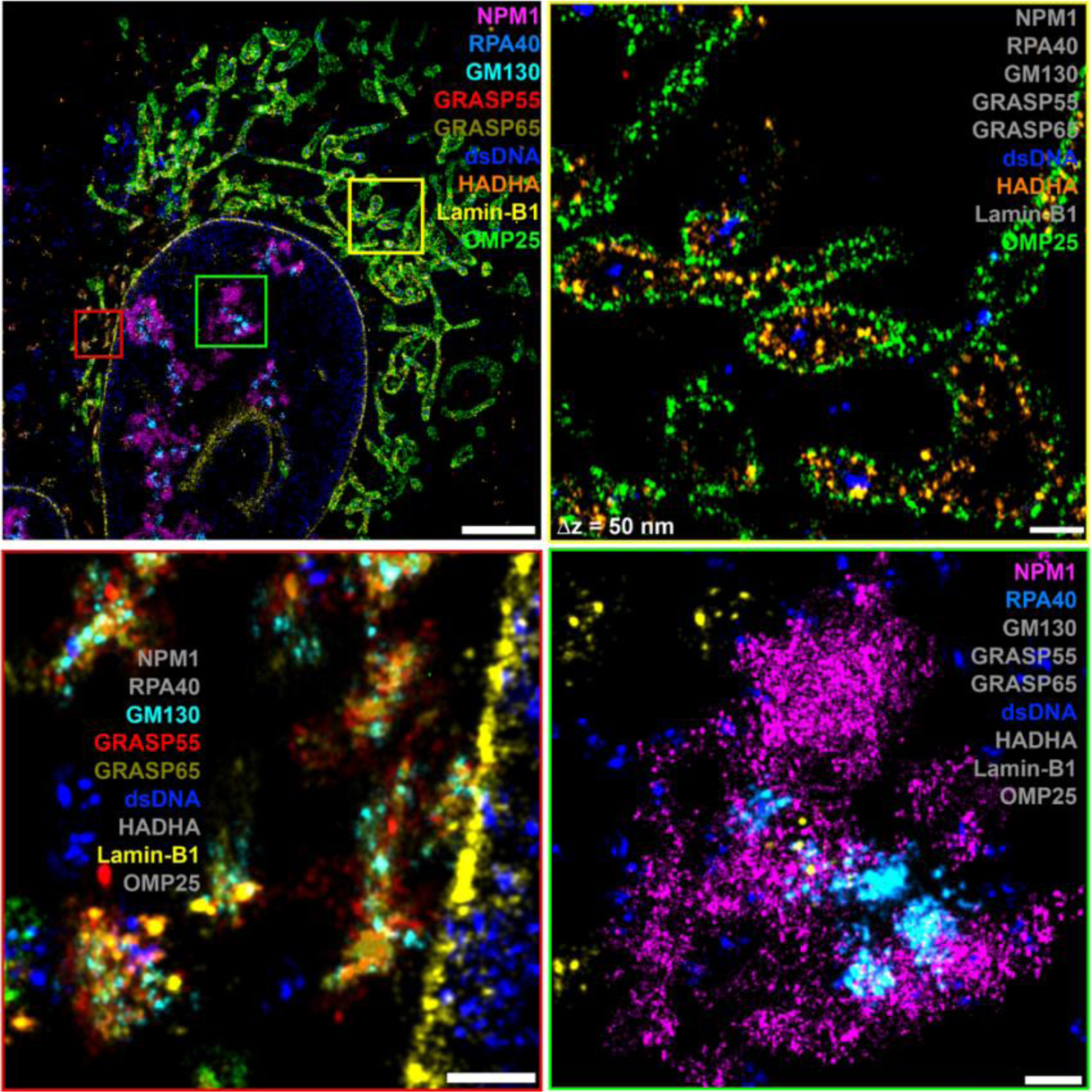
9-target FLASH-PAINT image of a U-2 OS cell. Nine different protein targets located at the Golgi complex, mitochondria, and the nucleus were imaged at super-resolution. The yellow, red, and green subpanels zoom in on mitochondria (yellow box), parts of the Golgi complex and nuclear envelope (red box), and a nucleolus (green box), respectively. For clarity, only a subset of proteins is shown in these subpanels; labels marked in gray are not shown. Scale bars: 5 µm (overview), 500 nm (zoom-ins).

### 9-plexed FLASH-PAINT resolves the molecular organization in primary cilia

Primary cilia function as cellular antenna that not only receive signals, but potentially transmit them by releasing vesicles from their tips^29, 30^. Their characteristic architecture includes a core microtubule axoneme surrounded by a specialized membrane enriched in GPCRs (e.g., Smo) and a transition zone (TZ) structure near the cilia base that gates entry into this privileged domain^31^. To obtain a comprehensive view of primary cilia, the spatial distribution of individual proteins which organize and are enriched in these sub-diffraction (<200 nm) compartments have to be combined. This task is, however, impeded by the different states (e.g., in response to stimuli, assembly and disassembly) primary cilia can exist in, which makes it difficult to combine data from different data sets.

We tested whether FLASH-PAINT can visualize cilia nanostructure in 3D and reveal characteristic protein combinations for individual cilia compartments. To directly conjugate FLASH-PAINT docking sites to antibodies against different cilia targets, we utilized a Light Activated Site-Specific Conjugation (LASIC) protocol ^32^ that directly conjugates the oligos to the primary antibodies. Alternatively, LASIC can be used to conjugate the oligos to secondary antibodies (**Figure S22a**)^32^. We imaged multiple ciliated cells in 3D with 9-plex FLASH-PAINT (**Figures 4, S22b** and **S22c**). Analyzing one cilium, we observed, as expected, cilium membrane proteins (pHSmo, INPP5E and Arl13b) as a tube that surrounded glutamylated and acetylated microtubules (Glu-tub and Ac-tub; yellow and blue boxes in **Figures 4a** and **4b**). The basal body distal appendage protein CEP164 appeared as a ring around the cilia base, with the TZ protein Rpgrip1l just distal to it (blue box, arrowheads **Figures 4a** and **4b**). Other cilia proteins, Sept2 and the cargo transporter Ift88 had more variable distributions. Comparing this 9-color super-resolution image with that of another cilium featuring a large bulbous tip (yellow box, asterisk, **Figure 4b**) revealed striking differences: the latter cilium showed a thinning of the axoneme membrane, represented by pH-Smo, just before the bulbous tip, as well as other varicosities (arrows, **Figure 4b**); interestingly, this was not apparent from the microtubule reporters, potentially as axoneme microtubule complexes can thin and become singlets as they approach the tip^33^.

**Figure 4.**
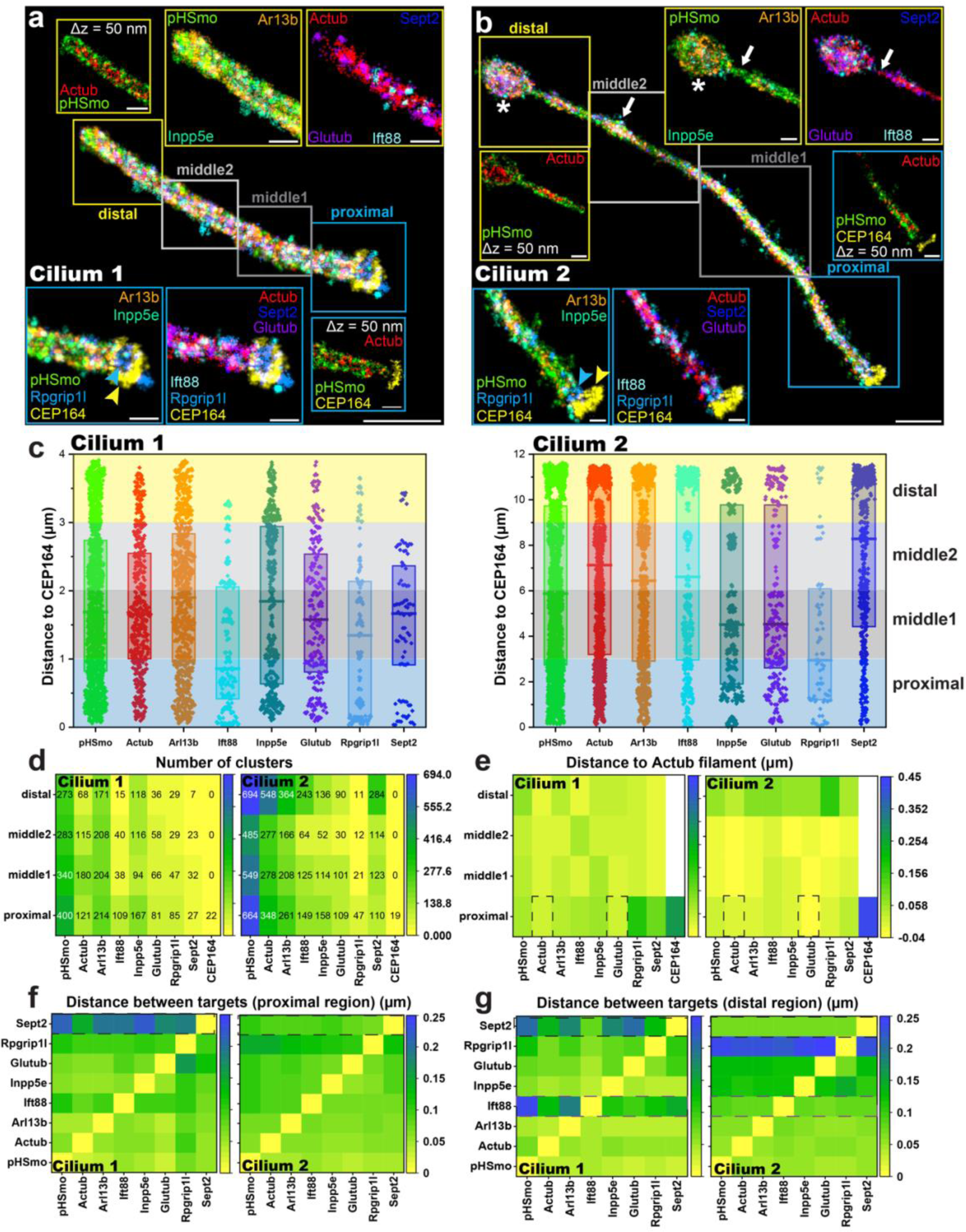
9-target FLASH-PAINT imaging of normal and bulbous-tip cilia in RPE-pHSmo cells. **(a-b)** FLASH-PAINT image of nine different protein targets at a normal (a; Cilium 1) and bulbous tip (b; Cilium 2) cilium. Quartiles along the length of the cilia are represented by square boxes. Subsets of the nine targets of the zoomed-in proximal and distal regions are shown in the blue and yellow boxes, respectively. The zoomed-in boxes labeled “Δz = 50 nm” show 50-nm thick cross sections of the acquired 3D super-resolution data sets to highlight the distribution of acetylated tubulin (Actub) inside the cilia. The yellow and blue arrowheads point at the basal body distal appendage CEP164 and the transition zone (TZ) Rpgrip1l markers, respectively. Between the bulbous tip (asterisk in (b)) and a varicosity (white arrow in middle2 box in (b)), thinning of the ciliary membrane (white arrows in yellow boxes in (b)) can be observed. **(c)** Bar plots summarizing the axial distribution of the nine different targets in the two cilia. For each target, the median (line) and 25% and 75% quartiles (bottom and top of the bar) are indicated. **(d)** Numbers of target clusters in each of the four regions for both cilia. **(e)** Median distances of target clusters to the central acetylated-tubulin (Actub) filament in each of the four regions for both cilia. In the proximal regions, Actub and Glutub axoneme targets show the shortest distance to the filament (black dashed-line rectangles). **(f, g)** Median distances between clusters of two different targets in the proximal (f) and distal (g) regions of both cilia Sept2 clusters are closer to all other targets in the normal cilium (Cilium 1) compared to the bulbous tip cilium (Cilium 2) (black dashed-line rectangles in (f) and (g)). In the distal region, a similar observation can be made for Ift88 (magenta dashed-line rectangles in (g)). Scale bars: 1 µm (overviews); 300 nm (zoom-ins).

We next analyzed the spatial distributions along the cilia axes. We localized single-molecule clusters relative to the pH-Smo signal and to a central filament generated from the Ac-tub signal (**Figure S22d**). In addition, since the TZ is close to the basal body, which has recently been shown to have a differential accumulation of Ac/Glu tubulin ^34^, we segmented the cluster data in quartiles based on their distance to CEP164 (proximal, middle1, middle2 and distal squares; **Figures 4a** and **4b**). Quantifying the cluster distribution in these quartiles revealed differences between the normal and bulbous cilia with Ift88 and Sept2 being enriched in the proximal and two middle regions of the normal cilium, yet recruited along the entire length of the cilium with the bulbous tip, the latter suggesting heightened activation (**Figures 4c** and **4d**)^35, 36^. When we measured the median distances from the central filament, Ac-tub and Glu-tub showed, as expected, the lowest values (dashed rectangles in **Figure 4e**). Compared to the normal cilium, the bulbous tip cilium showed a shorter distance at the proximal and middle regions for all targets (**Figure 4e**). Finally, when we analyzed the median distances between clusters of different proteins (see **Methods**), we observed that the Sept2 median distance to all other proteins was larger in the bulbous tip cilium than in the normal cilium in both the proximal and distal regions (dashed black rectangles, **Figures 4f** and **4g**)., In contrast, for Ift88 this was only the case for the distal region (dashed purple rectangles). These findings underscore the power of multiplexed super-resolution microscopy to identify both distinct nanoscale and longer-range states in primary cilia, including rare/transient stages, that would be missed in ensemble-averaged studies of cilia.

### 12-plexed FLASH-PAINT maps the spatial organization of the secretory pathway

We next tested FLASH-PAINT to better visualize the complex 3D structure of the Golgi. We used 12-plexed super-resolution imaging to study the spatial organization of the secretory pathway by highlighting components of ER exit sites (ERES), the ER-Golgi intermediate compartment (ERGIC), cis, medial and trans cisternae of the Golgi apparatus (GA), the trans Golgi network (TGN), and COPI and COPII vesicles in the same cell. The Golgi ribbon appeared, as expected, as a highly convoluted 3D structure proximal to the nuclear lamina in HeLa cells in interphase (**Figures 5a** and **S23**). ERES (TANGO1), ERGIC (ERGIC-53), and COPI (β′-COP) vesicles were distributed throughout the cytoplasm (**Figure 5b**). A cross section through the Golgi stack revealed the layered organization of cis-(GRASP65, GM130), medial-(ManII-GFP), and trans-cisternae (Golgin97, p230), and the trans-Golgi network (TGN46) (**Figures 5e** and **S23**). An *en face* view of the Golgi stack, revealed that Giantin localized at the rim of Golgi cisternae (**Figures 5c** and **5g**), consistent with EM data ^37^. COPI (β′-COP) vesicles were observed mostly at the periphery of the Golgi ribbon (**Figure 5f**), near their budding location^38, 39^. Most of the COPII coat (Sec31A) puncta were visibly larger than TANGO1 (ERES) puncta and usually one or more TANGO1 puncta decorated each Sec31A punctum (**Figure 5f**), supporting that multiple TANGO1 proteins surround the budding sites of ERES^40^.

**Figure 5.**
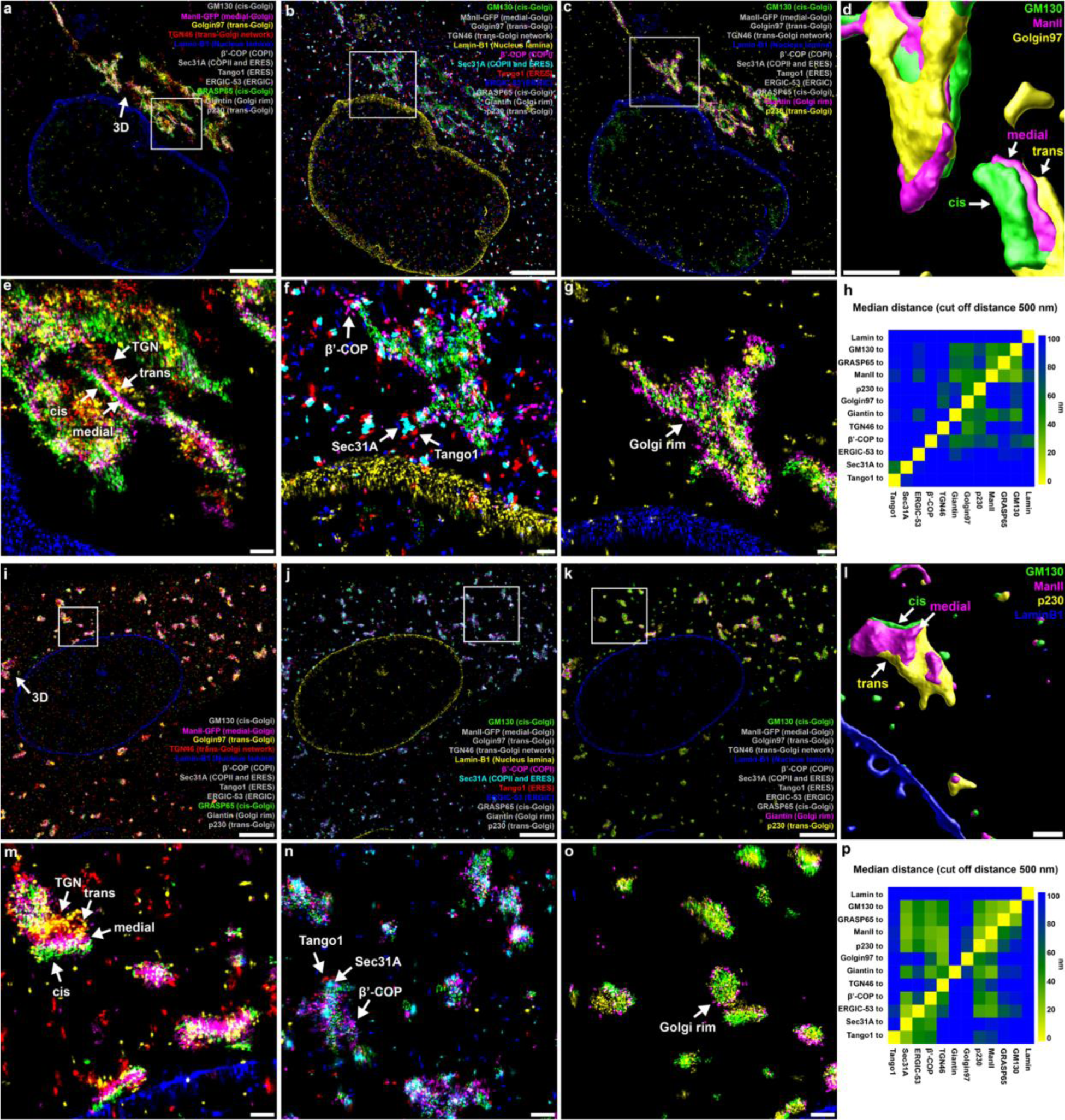
12-target FLASH-PAINT imaging of Golgi complexes in untreated and nocodazole-treated HeLa cells. **(a– c)** Overview of the Golgi complex and in the secretory pathway in a HeLa cell in interphase. Different subsets of protein targets are shown in the three images as indicated by the colored labels (gray-labeled targets not shown). **(d)** 3D surface reconstruction of cis, medial, and trans cisternae of the Golgi complex. **(e–g)** Zoomed in regions of the white boxes shown in (a–c), respectively, highlighting a side view of a Golgi stack revealing the sequential organization of the stack into cis cisternae, medial cisternae, trans cisternae and TGN (e), the spatial relationship between ERES, ERGIC and COPI and II vesicles (f), and an en face view of a Golgi stack showing Giantin located at the Golgi rim (g). **(h)** Median distances between localization events of different targets. Only localization events closer than 500 nm to each other were considered. Median distances >100 nm are shown in blue. **(i–p)** Representation equivalent to (a–h) of Golgi ministacks in a nocodazole-treated HeLa cell in interphase. Scale bars: 5 µm (a–c; I–k), 500 nm (d–g), 1 µm (l–o).

To visualize the 3D organization of the Golgi ribbon, we generated surfaces from the single-molecule localization data of GM130, ManII-GFP and Golgin97 and Lamin-B1 using a recently developed method^41, 42^ (**Figure 5d**). The median distances between the localizations of each label to those within a 500-nm range of all other labels were plotted as a heatmap (**Figure 5h**). Consistent with the expected organization of the Golgi stack, this quantification showed that stack-associated proteins (GM130, GRASP65, ManII-GFP, p230, Golgin97) were closer to each other than proteins outside this group.

Imaging nocodazole-treated cells in interphase with the same labels (**Figure S24**) revealed Golgi ministacks with the cis-to-trans hierarchy (**Figures 5i** and **5m**) and rim localization of Giantin (**Figures 5k** and **5o**) largely intact, supporting the long-standing hypothesis that nocodazole-induced ministacks represent a valid morphological model of native Golgi^37, 43^. Visually comparing this data to that of the Golgi apparatus in non-treated cells, suggested that the ministacks were more proximal to ERES as marked by Sec31A and TANGO1 (**Figures 5j** and **5n**). Quantifying the median distances of Golgi stack proteins to Sec31A between the two conditions showed a substantial reduction from >100 nm to the 50-nm range (**Figures 5p** and **S25**). This observation is consistent with the model that ER export is critical in Golgi regeneration^44, 45^. COPI vesicles were located at the periphery of Golgi ministacks (**Figures 5j** and **5n**), as they were in non-treated cells, suggesting that vesicle budding was not affected by nocodazole. Taken together, our data both visualize and quantify for the first time the complex organization of proteins of the secretory pathway in the same cell, providing powerful morphological context with molecular specificity for future research.

### FLASH-PAINT of whole cells charts the number and size of inter-organelle contact sites

In recent years, the contact between organelles has been recognized to play critical roles in coordinating cellular function, and dysfunction of such contacts may be associated with neurodegenerative disease^46^. Inspired by earlier work using diffraction-limited microscopy^47^, we imaged four different organelles, mitochondria (Tom20), the ER (Sec61beta), the Golgi (ManII) and lysosomes (Lamp1), at 3D super-resolution in a ∼2.5-µm thick volume across a Hela cell (**Figures 6** and **S26-S29**). For these experiments, we used fluorogenic Imagers, which enabled high-quality imaging of thick volumes deep in the cell in two ways: first, the fluorogenic state of the unbound state decreases the background of unbound probes in solution which is critical for large excitation volumes as used in highly inclined and laminated optical sheet (HILO)^48^ or epi-illumination. Second, the fluorogenic nature protects unbound Imagers from bleaching in solution. This is especially important for large excitation volumes since bleaching of Imagers in solution decreases the effective concentration of functional Imagers and thereby reduces the blinking frequency and data acquisition speed in a DNA-PAINT experiment.

**Figure 6.**
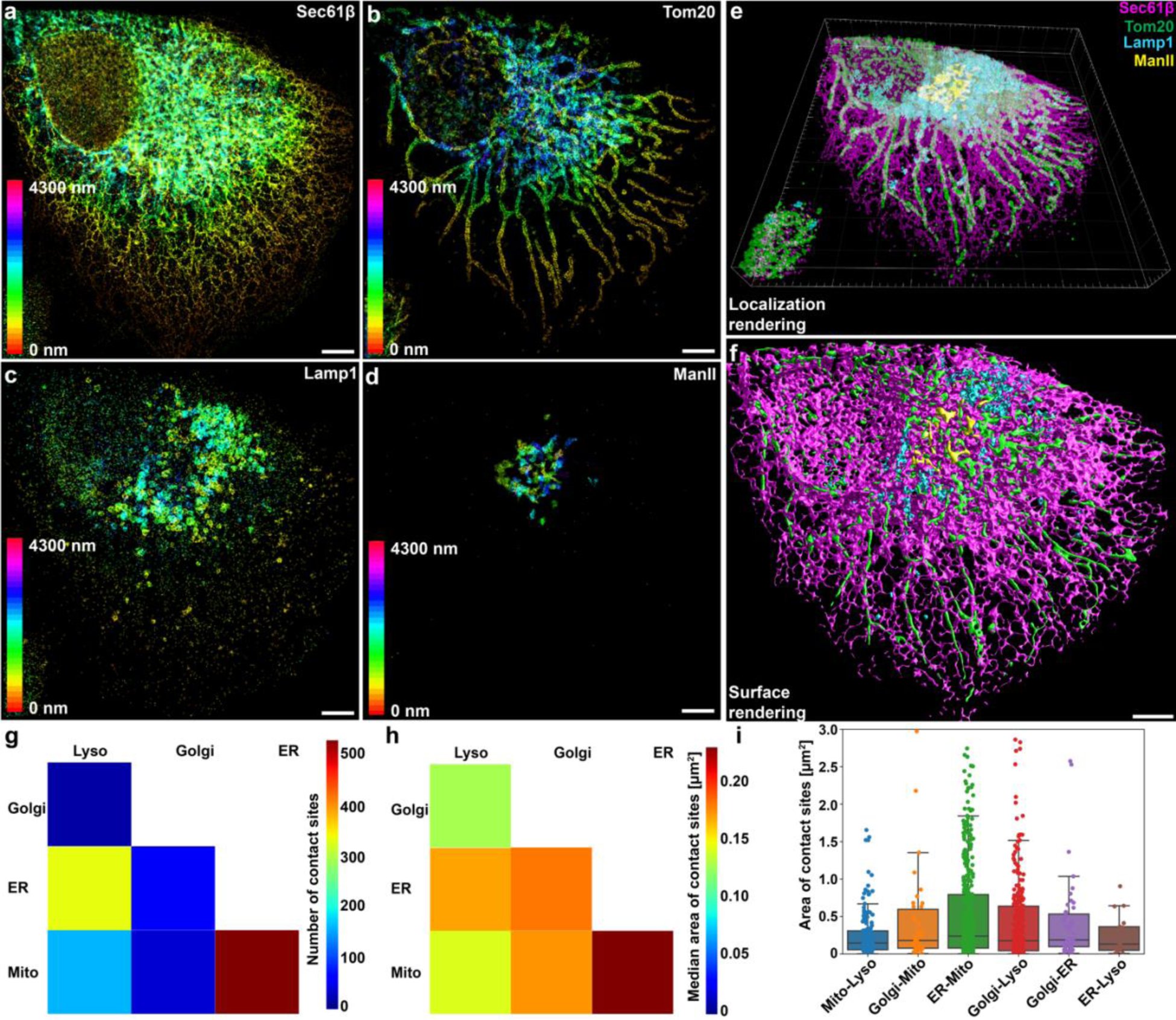
Volumetric multiplexed cellular organelle imaging at super-resolution using FLASH-PAINT. **(a-d)** Depth projection of the ER (**a**; Sec61β), mitochondria (**b**; Tom20), lysosomes (**c**; Lamp1) and Golgi complex (**d**; ManII-GFP) in a ∼2.5-µm thick 4-plexed FLASH-PAINT data set of a HeLa cell. (**e, f**) 3D rendering of localization data (e) and surface rendering (f) of the four labels. (**g, h**) Number of contact sites (g; defined as distance <100 nm) and median area of contact sites (h) between Golgi complex, lysosomes, ER and mitochondria. (**i**) Bar plots of the areas of the individual contact sites. The data points represent the areas of contact for all identified individual contacts between the organelles in the cell. For each type of contact, the median (line) and 25% and 75% quartiles (bottom and top of the bar) are indicated. Scale bars: 5 µm.

We collected 41 million localizations with an average localization precision of 16.6 nm in 173 minutes. Using surface reconstruction of the localized point clouds, we generated 3D representations of the imaged organelles^41, 49^. Using the organelle surfaces we quantified the number of contact sites between the different organelles (**Figure 6f**), defined as a spatial proximity of two membranes of <100 nm. The obtained contact site numbers were consistent with those extracted from diffraction-limited microscopy data by Valm *et al.*^47^. Imaging at super-resolution enabled us to additionally quantify the average area of contact sites. We found that the median of all contact sites was in the range between 0.1 µm^2^ and 0.2 µm^2^, independent of the pair of interaction partners. ER-mitochondria contact sites, which were the most abundant contacts, also showed the largest median values of their sizes and the largest size variance with about 20% of contact sites being larger than 1 µm^2^ (**Figures 6h** and **6i**).

## Discussion

With Transient Adapters and Erasers, we have introduced a new concept in FLASH-PAINT that rapidly switches a fluorescent probe from one target to another. Being based on DNA technology, up to 4^10^, i.e. more than 1 million, designs of (10 nt-long) Transient Adapters are theoretically available – far more than the ∼20,000 different proteins expressed in a cell^50^. While not all 1 million sequences are suitable options due to off-target binding, crosstalk, unwanted secondary structure formation and other effects, our concept yields effectively unlimited multiplexing capabilities for any currently practical proteomics study. Importantly, the same fluorescent Imager probe (or a handful if one wants to image multiple targets simultaneously in different colors) can be used repeatedly. FLASH-PAINT can therefore leverage the newest generation of DNA-PAINT probes, that are optimizes for speed^22, 23^ and fluorogenicity^24^ but are heavily constrained in their sequence design and are thus not directly suitable for highly multiplexed imaging. As demonstrated here, this combination enables the generation of super-resolution images of complex sub-cellular structures such as cilia or the Golgi complex at excellent quality, deep inside cells and in minutes rather than hours per imaged target.

Adapters binding stably, i.e. not transiently, have been successfully used in diffraction-limited and super-resolution microscopy^51–53^. While both types of adapters enable sequential labeling of many targets with just a few fluorescent probes, stable adapters suffer from the same problem that adapter-less sequential multiplexing approaches face: previously imaged targets need to be eliminated before imaging the next one. This is usually achieved by (i) removing the adapters (using dissociation buffers^19, 25, 26, 54^ or toehold-mediated displacement^55, 56^), (ii) permanently blocking them (with blocking strands that saturate the binding site the Imager probe normally binds to), or (iii) destroying them (with enzymes). However, all of these approaches require extensive incubation and washing periods which slow down data acquisition and can be inefficient, thereby causing crosstalk or background, and/or can damage the sample, especially if applied repeatedly over many imaging cycles.

Transient Adapters, in contrast to static adapters, by design easily dissociate from their targets without the need of toehold-mediated displacement or dissociation buffers. This fast and easy dissociation makes the sequence of the Transient Adapter that binds specifically to its target docking site readily accessible to the complementary Eraser strands. This, as we demonstrated (**Figures 2b**, **S16** and **S21-S24**), leads to highly efficient (99% to 99.8%) neutralization of the Transient Adapter. Since Erasers are specific to one particular docking site each, they do not quench the signal of other targets (in contrast to blocking strands described above that bind to the universal Imager probe binding site). Hence, neither the previous Transient Adapter nor its Eraser need to be washed out before the next imaging round. In fact, we can introduce the Eraser at the same time as the next Adapter (**Figures 2c** and **S17**, **Movies S1** and **S2**), which minimizes the transition time between imaging rounds.

Fully transitioning from one target to the next one required ∼1-10 minutes in our experiments (**Figures 2c** and **S17**, **Movies S1** and **S2**). While much faster than alternative sequential multiplexing approaches, we point out, that we did not utilize any flow chamber in our proof-of-concept experiments and were therefore limited by diffusion. We believe that the introduction of flow, combined with further optimized adapter dissociation rate constants, can reduce the transition time between targets to less than 1 minute. We have demonstrated here up to 12-plex imaging which we considered sufficient to demonstrate broad cell biological utility of FLASH-PAINT. The primary obstacles to extending FLASH-PAINT to more targets in this study were the limited access to validated, high-quality antibodies and lacking automated microfluidics – both not of fundamental nature. While we have focused here on immunolabeling, we believe that FLASH-PAINT will be equally useful in spatial transcriptomics studies and to trace DNA in the nucleus using fluorescence in situ hybridization. Barcoded multiplexing schemes, as demonstrated by MERFISH^52^ and SeqFISH+^53^, allow for 1,000-fold and higher multiplexing with only tens of adapters. The low crosstalk of our Transient Adapters has the potential to minimize error rates in barcoded multiplexing. This in turn should enable researchers to use more barcodes from the codebook (i.e., barcodes with a smaller Hamming distance) and thereby provide access to more target species with fewer rounds of imaging.

Even more broadly, we believe that Transient Adapters will find wide application in diffraction-limited spatial omics approaches. Localization of single blinking molecules is only needed for super-resolution – if that is not required, the concentration of the Imager probe can be increased to provide diffraction-limited images as shown in **Figure 2c**, **Figure S17** and **Movies S1 & S2**. In contrast to many established techniques in the field^19, 28^, Transient Adapters and Erasers allow for rapid exchange of labels without the use of harsh, time-consuming treatment steps, such as stripping probes off the sample or photocleaving or bleaching them, between imaging rounds. Additionally, Transient Adapters and Erasers are inexpensive: unlabeled oligos as used here cost only a fraction of their dye-labeled counterparts.

Importantly, FLASH-PAINT is not conceptually limited to imaging a single color at a time. We anticipate that it can be readily combined with Imager probes of multiple fluorescent colors^24, 57^ Furthermore, we envisage synergies with innovative simultaneous multicolor approaches such as super-multiplex vibrational imaging^58^. With synergies such as these and a broad spectrum of potential application that extends to transcriptomics and chromatin tracing, we believe that FLASH-PAINT will be an enabling technology in a wide range of biological applications.

## Acknowledgments

We thank Zach Marin and Lukas Fuentes for the introduction to the surface shrink wrapping approach. We thank Sylvi Stoller for her help with initial kinetics characterization experiments of an early version of the Transient Adapter design. We thank Kenny Chung for fruitful discussions regarding fluorogenic Imagers. We thank Pietro de Camilli and Nisha Mohd Rafiq for their insights about the biological application of FLASH-PAINT. An Andor Dragonfly microscope in the CINEMA imaging facility at Yale was used for our imaging experiments. F.S. acknowledges support from the Human Frontier Science Program (LT000056/2020-C). J.B. acknowledges support by the Wellcome Leap Foundation.

## Author contributions

F.S. and J.B. conceived and designed the study. F.S. performed all experiments except the cilia experiments. F.R.-M. performed the cilia experiments. F.S. and F.R.-M. analyzed the cilia data. P.K. contributed to the design of the 9-plexed experiment. M.S. and F.S. designed, performed, and analyzed the Golgi experiment. F.S. and J.B. wrote the manuscript with input from all authors. D.K.T. and J.E.R. supervised parts of the study. J.B supervised the study.

## Inclusion and Diversity

We support inclusive, diverse, and equitable conduct of research.

## RESOURCE AVAILABILITY

### Lead contact

Further information and requests for resources and reagents should be directed to and will be fulfilled by the Lead Contacts, Florian Schueder and Joerg Bewersdorf.

### Materials availability

Requests for reagents should be directed to and will be fulfilled by the lead contact.

### Data and code availability

All raw data is available upon reasonable request from the authors due to the extensive data size of the data. Data were processed, analyzed and visualized using the open-source software packages Picasso (https://github.com/jungmannlab/picasso), Python Microscopy (https://github.com/python-microscopy/python-microscopy) and the commercially available Imaris Software (Version 10.0).

## Materials and Methods

### MATERIALS

#### Materials

Unmodified DNA oligonucleotides, Cy3b-modified DNA oligonucleotides and biotinylated DNA oligonucleotides were purchased from Integrated DNA Technologies (IDT). M13mp18 scaffold (cat: N4040S) was obtained from New England BioLabs. Tris 1 M pH 8.0 (cat: AM9856), EDTA 0.5 M pH 8.0 (cat: AM9261), Magnesium 1 M (cat: AM9530G) and Sodium chloride 5 M (cat: AM9759) were obtained from Ambion. Ultrapure water (cat: 10977015) was purchased from Gibco. 200 µL PCR tubes (cat: AB-0620) were obtained from Thermo Scientific. Polyethylene glycol (PEG)-8000 (cat: 89510-250G-F) was purchased from Sigma. Streptavidin (cat: S-888) was purchased from Thermo Fisher. BSA-Biotin (cat: A8549) was obtained from Sigma-Aldrich. Tween 20 (cat: P9416-50ML), glycerol (cat: 65516-500ml), methanol (cat: 32213-2.5L), protocatechuate 3,4-dioxygenase pseudomonas (PCD) (cat: P8279), 3,4-dihydroxybenzoic acid (PCA) (cat: 37580-25G-F) and (+−)-6-hydroxy-2,5,7,8-tetra-methylchromane-2-carboxylic acid (Trolox) (cat: 238813-5 G) were ordered from Sigma. Sodium hydroxide (cat: P3911-1kg) was purchased from Sigma Aldrich. Potassium chloride (cat: 3624-01) was ordered from Baker Analyzed A.C.S. Reagent. 30 mL Syringes (cat: 302832) were obtained from BD. Biocompatible silicone tubing (cat: 10831), flow chambers 6-well µ-Slide VI^0.5^ (cat: 80607) and glass-bottomed 8-well µ-slides (cat: 80827) were obtained from ibidi. 8-wells 1.5H glass bottom chambers (cat: C8-1.5H-N) were purchased from Cellvis. 15 mL (cat: 352096) and 50 mL (cat: 352070) Polypropylene Conical Tubes and tissue culture flasks (cat: 353136) were purchased from FALCON. Dulbecco’s Modified Eagle medium (DMEM) (cat: 21063-929), McCoy’s 5A Medium (cat: 16600-082), Opti-MEM (cat: 31985-070), 0.05% Trypsin-EDTA (cat: 25300-054), Fetal Bovine Serum (FBS) (cat: 16000-044) and 1× Phosphate Buffered Saline (PBS) pH 7.2 (cat: 10010-023), was ordered from gibco. 10% Formalin (cat: HT501128-4L), heat inactivated FBS (cat: F4135-500ML) and 1mg/mL fibronectin (cat: F0895-2MG) were purchased from Sigma-Aldrich. HeLa CRM-CCL-2 cells (cat: CRM-CCL-2), U-2 OS cells (cat: HTB-96), COS-7 cells (cat: CRL-1651) and hTERT-RPE cells (cat: CRL-4000) were obtained from ATCC. Paraformaldehyde (cat: 15710) and glutaraldehyde (cat: 16219) were obtained from Electron Microscopy Sciences. Bovine serum albumin (cat: 001-000-162) was ordered from Jackson ImmunoResearch. Triton X-100 (cat: T8787-60ML) was purchased from Sigma. Antibodies against GM130 (cat: 610822), Sec31A (COPII) (cat: 612350) and p230 (cat: 611280) were obtained from BD Biosciences. Antibodies against LaminB1 (cat: ab16048), HADHA (cat: ab110302), GRASP65 (cat: ab174834), dsDNA (cat: ab3519) and Septin2 (ab187654) were obtained from abcam. Antibodies against GRASP55 (cat: 10598-1-AP), GM130 (cat: 11308-1-AP), TGN46 (cat: 10598-1-AP), Inpp5e (17797-1-AP), Arl13b (17711-1-AP), Ift88 (13967-1-AP), CEP164 (22227-1-AP) and, RPGRIP1L (55160-1-AP) were purchased from Proteintech. Antibodies against Tom20 (cat: sc-11415), RPA40 (cat: sc-374443) were ordered from Santa Cruz. Antibodies against NPM1 (cat: NB600-1030) were obtained from Novus Bio. Antibodies against GOLGB1 (Giantin) (cat: HPA011555), Anti-MIA3 (Tango1) (cat: HPA055922), acetylated-tubulin (T6793) and, anti-alpha-tubulin (cat: T5168) were ordered from Sigma. Antibodies against GOLGA1_1 Golgin-97 (cat: HPA044329) were purchased from Atlas Antibodies. Antibodies against LMAN1 ERGIC-53 (cat: MA5-25345) were ordered from Invitrogen. Antibodies against Glutamylated-tubulin (AB3201) were ordered from Millipore. Antibodies against Lamp1 (9091) were purchased from Cell Signaling Technology.

Antibodies against mCherry (GT844 and GT857) were obtained from GeneTex. Antibodies against COPI (CMIA10) were customary made in the Rothman lab. DNA-labeled secondary anti-rabbit antibodies, DNA-labeled secondary anti-mouse antibodies and DNA-labeled GFP nanobodies were custom-ordered from Massive Photonics. Oligos conjugated to the OyOlink probe were purchased from AlphaThera.

### METHODS

#### Buffers

Three buffers were used for sample preparation and imaging: Buffer A (10 mM Tris-HCl pH 7.5, 100 mM NaCl, 0.05% Tween 20, pH 7.5); Buffer B (10 mM MgCl_2_, 5 mM Tris-HCl pH 8, 1 mM EDTA, 0.05% Tween 20, pH 7.5), and Buffer C (1× PBS, 500 mM NaCl). For the experiments shown in **Figures 1b, 1e, 3, 5, 6, S10 – S15, S18 – S21, S23, S24** and **S26 – S29**, the imaging buffers were supplemented with: 1× Trolox, 1× PCA and 1× PCD.

#### Trolox, PCA and PCD

100× Trolox: 100 mg Trolox, 430 μL 100% Methanol, 345 μL 1 M NaOH in 3.2 mL H_2_O. 40× PCA: 154 mg PCA, 10 mL water and NaOH were mixed, and pH was adjusted to 9.0. 100× PCD: 9.3 mg PCD, 13.3 mL of buffer (100 mM Tris-HCl pH 8, 50 mM KCl, 1 mM EDTA, 50% glycerol).

#### DNA origami self-assembly

All DNA origami structures were designed with the Picasso design tool^13^ (see **Figure S2**). Self-assembly of DNA origami was accomplished in a one-pot reaction with 50 μL total volume, consisting of 10 nM scaffold strand (sequence see **Table S6**), 100 nM folding staples (**Tables S7-S13**), 10 nM (or 1 µM (**Figures 1e** and **S10-S14**)) biotinylated staples (**Table S14**), and 1 μM of docking site strands (List of DNA-PAINT or FLASH-PAINT handles see **Table S2**) in folding buffer (1× TE buffer (10 mM Tris and 1 mM EDTA) with 12.5 mM MgCl_2_). The reaction mix was then subjected to a thermal annealing ramp using a thermocycler. The reaction mix was first incubated at 80 °C for 5min, then cooled from 60 to 4 °C in steps of 1 °C every 3.21 min, and then held at 4 °C.

#### DNA origami PEG purification

DNA origami structures featuring letters, a 10-nm and a 20-nm-grid (**Figures 1e** and **S10-S14**) were purified via three rounds of PEG precipitation by adding the same volume of PEG-buffer (15% PEG-8000, 500 mM NaCl in 1× TE buffer, pH 8.0), centrifuging at 14,000 g at 4 °C for 30 min, removing the supernatant, and resuspending in folding buffer.

#### DNA origami sample preparation

For DNA origami sample preparation, a µ-Slide VI^0.5^ (ibidi) was used as sample chamber. First, 100 μL of biotin-labeled bovine albumin (1 mg/mL, dissolved in buffer A) were flushed into the chamber and incubated for 5 min. The chamber was then washed with 500 μL of buffer A. A volume of 100 μL of streptavidin (0.5 mg/mL, dissolved in buffer A) was then flushed through the chamber and allowed to bind for 5 min. After washing with 500 μL of buffer A and subsequently with 500 μL of buffer B, 100 μL of biotin-labeled DNA structures (∼200 pM) in buffer B were flushed into the chamber and incubated for 8 min. The chamber was then washed with 500 μL of buffer B. Finally, 100 μL of the Imager solution in the corresponding imaging buffer (**Table S15**) was flushed into the chamber.

#### Plasmids

For labeling the outer membrane of mitochondria (**Figure 3**), we expressed GFP-OMP25 from a plasmid as described before^59^. For labeling medial Golgi cisternae (**Figures 5** and **6**), we expressed GFP-ManII from a plasmid as described previously^17^. mCherry-Sec61β was acquired from Addgene (plasmid 49155).

#### Cell culture

HeLa cells and COS-7 cells were cultured in DMEM supplemented with 10% Fetal Bovine Serum (FBS). U-2 OS cells were cultured in McCoy’s 5A Medium supplemented with 10% FBS. The night before immunolabeling, cells were seeded on ibidi 8-well glass coverslips at ∼30,000 cells/well. RPE-pHSmo cells were maintained in DMEM/F12 supplemented with 10% FBS, 1× Pen/Strep, 1× non-essential amino acids and 1 mM sodium pyruvate. For ciliogenesis, 250 µL from a 50,000 cells/mL suspension of RPE-pHSmo cells were plated into 4 wells of an 8-well cellvis chamber that was coated for 1 h with 10 µg/mL fibronectin. The cells were incubated for two days at 37 °C to reach confluency. On the third day, the medium was changed to medium supplemented with 0.5% FBS to start the starvation period for another two days.

#### Transient transfection

Transfections were performed using a Super Electroporator NEPA21 Type II (Nepa Gene). Cells were concentrated to approximately 1 million cells in 90 μL in an electroporation cuvette (Bulldog Bio; 12358-346) to which 10 μL of ∼1 μg/μL of plasmid DNA were added. Cells were electroporated using the following program: 125-V poring pulse, 3-ms pulse length, 50-ms pulse interval, two pulses, with decay rate of 10% and + polarity, followed by a 25-V transfer pulse, 50-ms pulse length, 50-ms pulse interval, five pulses, with a decay rate of 40% and ± polarity.

#### Golgi ministack induction

Golgi ministacks were induced by treating HeLa cells with 5 µg/mL of nocodazole in culture medium for 4 h at 37 °C before fixation.

#### Cell fixation and labeling for Figures 2 and S17b

Cells were fixed with 3% PFA and 0.1% GA for 15 min. After four washes (30 s, 60 s, 2× 5 min) cells were blocked and permeabilized with 3% BSA and 0.25% Triton X-100 at room temperature for 1 h. Next, cells were incubated with primary antibodies (**Tables S16** and **S17**) in 3% BSA and 0.1% Triton X-100 at 4 °C overnight. The next day after four washes (30 s, 60 s, 2× 5 min), cells were incubated with secondary antibodies for ∼2 h at room temperature. Next, after four washes (30 s, 60 s, 2× 5 min), the sample was post-fixed with 3% PFA and 0.1% GA for 10 min. Finally, samples were washed three times with 1× PBS for 5 min each before adding the imaging solution.

#### Cell fixation and labeling for **Figures 3 and S21**

Cells were fixed with 4% PFA for 1 h. After four washes (30 s, 60 s, 2× 5 min) cells were blocked and permeabilized with 3% BSA and 0.25% Triton X-100 at room temperature for 1 h. Next, cells were incubated with primary antibodies against GM-130 and LaminB1 (**Tables S16** and **S17**) in 3% BSA and 0.1% Triton X-100 at 4 °C overnight. The other primary antibodies were pre-incubated with the corresponding nanobodies (**Tables S16** and **S17**) at 4 °C overnight^60^. The next day, after four washes (30 s, 60 s, 2× 5 min), cells were incubated with the nanobodies corresponding to anti-GM130 (host: mouse) antibody and anti-LaminB1 (host: rabbit) antibody for ∼2 h at room temperature. Next, to block unlabeled epitopes, unlabeled excess secondary nanobodies were added to pre-incubation antibody and nanobody mixes at room temperature for 5 min. Next, the cells were incubated with the pooled antibody and nanobody mix for ∼2.5 h at room temperature. After four washes (30 s, 60 s, 2× 5 min) the sample was post-fixed with 3% PFA and 0.1% GA for 10 min. Finally, samples were washed three times with 1× PBS for 5 min each before adding the imaging solution.

#### Cell fixation preserving cilia (**Figures 4 and S22**)

After ciliogenesis induction, RPE-pHSmo cells were washed with 1× PBS and fixed with 10% Formalin for 15 min. Next, cells were washed three times with 1× PBS and permeabilized with PBS/0.1% Triton X-100 (PBST) for 10 min. Following permeabilization, the cells were washed with PBST and blocked with 3% BSA/PBST solution for 1 h. To conjugate cilia-targeted primary antibodies (**Tables S16 and S17**) to binder oligo-OyOlink molecules, 1 µg of purified antibody was mixed with 1 µg OyOlink (1:3 molar ratio) in a total of 10 µL with PBS in a 100 µL clear PCR tube. The tubes were then incubated for 2 h on a UV transilluminator box equipped with a 365-nm excitation light source. After light-induced cross-linking, the volumes were mixed and 200 µL of 3% BSA/PBST were added. 1 µL of 2.5 µM Nano-GFP A3 and 0.5 µL of mouse anti-acetylated tubulin were added to the mixture. Then 150 µL of this solution was added to one of the wells with the ciliated pHSmo cells and incubated at 4 °C overnight. The next day, the cells were washed three times with PBST for 5 min each and incubated for 2 h with anti-mouse A19 secondary antibody diluted 1:500 in blocking buffer. The sample was then washed three times with PBST for 5 min each, twice with 1× PBS and incubated for 10 min with 10% PFA and 0.1% GA. After post-fixation, the samples were washed three times with 1× PBS each and stored at 4 °C until imaging.

#### Cell fixation preserving Golgi complex (**Figures 5**, S23 and S24)

Cells were fixed with 4% PFA for 30 min. After four washes (30 s, 60 s, 2× 5 min), cells were blocked and permeabilized with 3% BSA and 0.25% Triton X-100 at room temperature for 1 h. Next, cells were incubated with the anti-MIA3 antibody, anti-p230 antibody, and the GFP-Nanobody in 3% BSA and 0.1% Triton X-100 at 4 °C overnight. Additionally, all other primary antibodies were pre incubated with the corresponding nanobodies (**Tables S16** and **S17**) at 4 °C overnight. The next day, after four washes (30 s, 60 s, 2× 5 min) cells were incubated with the nanobodies corresponding to anti-MIA3 antibody and anti-p230 antibody for ∼2 h at room temperature. Next, unlabeled excess secondary nanobodies (to block unlabeled epitopes) were added to pre-incubation antibody-nanobody mixes at room temperature for 5 min. Next, the cells were incubated with the pooled antibody-nanobody mix for ∼2.5 h at room temperature. After four washes (30 s, 60 s, 2× 5 min), the sample was post-fixed with 3% PFA and 0.1% GA for 10 min. Finally, samples were washed three times with 1× PBS for 5 min each before adding the imaging solution.

#### Cell fixation preserving ER, Golgi complex, lysosomes and mitochondria (**Figures 6 and S26** – S29)

Cells were fixed with 3% PFA and 0.1% GA for 15 min. After four washes (30 s, 60 s, 2× 5 min), cells were blocked and permeabilized with 3% BSA and 0.25% Triton X-100 at room temperature for 1 h. Next, cells were incubated with primary antibodies and nanobodies (**Tables S16** and **S17**) in 3% BSA and 0.1% Triton X-100 at 4 °C overnight. The next day, cells were incubated with secondary antibodies for ∼2 h at room temperature after four washes (30 s, 60 s, 2× 5 min). Next, after four washes (30 s, 60 s, 2× 5 min), the sample was post-fixed with 3% PFA and 0.1% GA for 10 min. Finally, samples were washed three times with 1× PBS for 5 min each before adding the imaging solution.

#### Cell fixation preserving nuclear lamina and nucleoli (Figure S17a)

Cells were fixed with 2.4% PFA for 30 min. After four washes (30 s, 60 s, 2× 5 min), cells were blocked and permeabilized with 3% BSA and 0.25% Triton X-100 at room temperature for 1 h. Next, cells were incubated with primary antibodies (**Tables S16** and **S17**) in 3% BSA and 0.1% Triton X-100 at 4 °C overnight. The next day, after four washes (30 s, 60 s, 2× 5 min), cells were incubated with secondary antibodies for 2 h at room temperature. Next, after four washes (30 s, 60 s, 2× 5 min), the sample was post-fixed with 3% PFA and 0.1% GA for 10 min. Finally, samples were washed three times with 1× PBS for 5 min each before adding the imaging solution.

#### Cell Fixation preserving mitochondria (**Figures S15 and S18** – S20)

Cells were fixed with 3% PFA and 0.1% GA for 15 min. After four washes (30 s, 60 s, 2× 5 min), cells were blocked and permeabilized with 3% BSA and 0.25% Triton X-100 at room temperature for 1 h. Next, cells were incubated with primary antibodies against Tom20 (**Tables S16** and **S17**) in 3% BSA and 0.1% Triton X-100 at 4 °C overnight. The next day, after four washes (30 s, 60 s, 2× 5 min), cells were incubated with secondary antibodies for 2 hours at room temperature. Next, after four washes (30 s, 60 s, 2× 5 min), the sample was post-fixed with 3% PFA and 0.1% GA for 10 min. Finally, samples were washed three times with 1× PBS for 5 min each before adding the imaging solution.

#### Super-resolution microscope setup

Fluorescence imaging was carried out on an inverted Nikon Eclipse Ti2 microscope (Nikon Instruments) with a Perfect Focus System, equipped with an Andor Dragonfly unit. The Dragonfly was used in the BTIRF mode, applying an objective-type TIRF or HiLo configuration with an oil-immersion objective (Nikon Instruments, Apo SR TIRF 60×, NA 1.49, Oil). For the acquisition of **Movie S1** and **Movie S2** the Dragonfly was used in confocal mode (40-µm pinhole disk). For excitation, a 561-nm laser (1 W nominal laser power) was used. The beam was coupled into a multimode fiber going through the Andor Borealis unit reshaping the beam from a Gaussian profile to a homogenous flat top. As dichroic mirror, a CR-DFLY-DMQD-01 was used. Fluorescence light was spectrally filtered with an emission filter (TR-DFLY-F600-050) and imaged with a scientific complementary metal oxide semiconductor (sCMOS) camera (Sona 4BV6X, Andor Technologies) without further magnification, resulting in an effective pixel size of 108 nm. Three-dimensional super-resolution imaging was performed by introducing astigmatism via a cylindrical lens in front of the camera.^61^

#### Imaging conditions

A detailed summary of all DNA origami experiments can be found in **Table S15** and for all cell experiments in **Table S17**. Additionally, we provide a high-level summary of all experiments in **Table S18**.

#### Image analysis

Raw fluorescence microscopy images were subjected to spot-finding and subsequent super-resolution reconstruction, drift correction, filtering and alignment using the ‘Picasso’ software package^13^. x, y and z drift correction were performed with a redundant cross-correlation which is integrated in the same software package. The surface reconstruction from localization data and the subsequent analysis of the contact sites were done using PYMEVisualize^42^. To identify and quantify clusters and distances on the cilia 9-plex data set the individual Picasso reconstructed cilium datasets were loaded into Imaris (Oxford instruments, version 10.0) to generate surfaces that were used to mask the localization data for each target at the cilium. After the surface mask was applied, localizations were processed with a Gaussian filter equivalent to one-pixel size. The filtered data was then used to generate spots using the Imaris spot detection algorithm to represent the size of the localization clusters. These spots were used to quantify the number of cluster and distances between targets. In addition, the Actub clusters were used to generate a filament representing the axoneme location along the length of the cilium.

