## Supplemental Information for "Unraveling cellular complexity with unlimited multiplexed super-resolution imaging"

### Derivation of the effective association rate of Imager probes binding to DNA origami in the presence of Transient Adapters

The observed number of transient binding events per time unit for a given DNA origami as shown in **Figure 1c and d**, can be described by the average dark time, i.e. the time no Imager probe is bound to a DNA origami (depending on the design of the DNA origami either via a Transient Adapter or directly). These times $\tau_{off,TA-mediated}$ and $\tau_{off,direct}$ can be described as functions of the Imager probe concentration $c_{Im}$ that was added to the imaging buffer by introducing effective association rates $k_{a,eff,TA-mediated}$ and $k_{a,eff,direct}$ for the Transient Adapter-mediated and the direct binding case, respectively (Definitions are summarized at the end of this derivation):

$$\tau_{off,TA-mediated} = \frac{1}{k_{a,eff,TA-mediated} * c_{Im}} \quad (1)$$

$$\tau_{off,direct} = \frac{1}{k_{a,eff,direct} * c_{Im}} \quad (2)$$

The effective association rate $k_{a,eff,TA-mediated}$ of Imager probes binding to a DNA origami docking site via Transient Adapters is influenced by both (i) the occupancy of the docking site by a Transient Adapter and (ii) the affinity between the Transient Adapter and Imager probes. The latter not only affects how efficiently Imager probes are recruited to the DNA origami, but Transient Adapters in solution also compete for these Imager probes and thereby reduce the pool of Imager probes readily available to bind to a Transient Adapter bound to the DNA origami target. This latter phenomenon also affects $k_{a,eff,direct}$ since both DNA origami species are imaged in the same sample.

To derive $k_{a,eff,TA-mediated}$, we make two assumptions: an Imager probe can only bind to a docking site when a Transient Adapter strand is present. Second, only the unbound fraction of Imager probes present in the solution can bind to a docking site. $k_{a,eff,TA-mediated}$, can be described as the product of the Duty Cycle, *D*, i.e. the fraction of time a docking site is occupied by a Transient Adapter, and the association rate constant of Imager probes binding to a Transient Adapter, $k_{a,Im}$:

$$k_{a,eff,TA-mediated} = D * k_{a,Im} \quad (3)$$

The Duty Cycle can be expressed as:

$$D = \frac{\tau_{on,TA}}{\tau_{on,TA} + \tau_{off,TA}} \quad (4)$$

The average time $\tau_{off,TA}$ where no Transient Adapter is bound at a docking site depends on both the concentration of the Transient Adapter, $c_{TA}$, and the association rate constant of Transient Adapters binding to a docking site, $k_{a,TA}$:

$$\tau_{off,TA} = \frac{1}{c_{TA} * k_{a,TA}} \quad (5)$$

For a 50-nM Transient Adapter concentration, an average binding time of 100 s, and an association rate of 2 x $10^6$ $M^{-1}s^{-1}$, the Duty Cycle is, for example, 91%.

As discussed above, a high concentration of Transient Adapters in solution, i.e. not bound to any docking site, will cause a non-negligible fraction of Imager probes to be bound to these Transient Adapters without producing localizable signal. Only the free fraction of unbound Imager probes, *f*, is available to bind to Transient Adapters bound to docking sites, reducing the concentration of Imager probes in solution below what was initially added to the imaging buffer:

$$c_{Im,free} = f * c_{Im} \qquad (6)$$

Following Jarmoskaite et al.[1], the fraction *f* of unbound Imager probes can be expressed as:

$$f = 1 - \frac{(c_{Im}+c_{TA}+K_D) - \sqrt{(c_{Im}+c_{TA}+K_D)^2 - 4*c_{Im}*c_{TA}}}{2*c_{Im}} \qquad (7)$$

Here, $K_D$ is the equilibrium dissociation constant between Transient Adapters and Imager probes:

$$K_D = \frac{\tau_{off,Im}*c_{Im}}{\tau_{on,Im}} \qquad (8)$$

$K_D$ can be estimated from measuring the average on and off times of Imager probes bound to DNA origamis featuring the same Imager probe binding site as the Transient Adapter at $c_{TA}$ = 0. Here we assume that these times depend only on the oligonucleotide sequence while possible effects from the surrounding environment (DNA origami vs. Transient Adapter) are negligible.

Combining **Equations 3-8**, the effective association rate constant of Imager probes binding to a docking site via Transient Adapters as a function of $c_{TA}$ can be expressed as:

$$k_{a,eff,TA-mediated}(c_{TA}) = \frac{\tau_{on,TA}}{\tau_{on,TA}+\frac{1}{c_{TA}*k_{a,TA}}} * \left(1 - \frac{\left(c_{Im}+c_{TA}+\frac{\tau_{off,Im}*c_{Im}}{\tau_{on,Im}}\right) - \sqrt{\left(c_{Im}+c_{TA}+\frac{\tau_{off,Im}*c_{Im}}{\tau_{on,Im}}\right)^2 - 4*c_{Im}*c_{TA}}}{2*c_{Im}}\right) * k_{a,Im} \qquad (9)$$

For comparison: the effective association rate $k_{a,eff,direct}$ as a function of $c_{TA}$ for Imager probes directly binding to a complementary docking site on a DNA origami in the presence of Transient Adapters in solution will equally be reduced by a factor *f* (i.e. **Equation 6** applies) compared to $k_{a,Im}$, but is independent of the Duty Cycle *D*. It can be expressed as:

$$k_{a,eff,direct}(c_{TA}) = \left(1 - \frac{\left(c_{Im}+c_{TA}+\frac{\tau_{off,Im}*c_{Im}}{\tau_{on,Im}}\right) - \sqrt{\left(c_{Im}+c_{TA}+\frac{\tau_{off,Im}*c_{Im}}{\tau_{on,Im}}\right)^2 - 4*c_{Im}*c_{TA}}}{2*c_{Im}}\right) * k_{a,Im} \qquad (10)$$

The solid curves in **Figure 1d** were calculated using **Equations 9** and **10** and the following values:

$$c_{Im} = 10\ nM$$
$$\tau_{on,Im} = 0.25\ s$$
$$\tau_{on,TA} = 100\ s$$
$$k_{a,Im} = 45 * 10^6\ M^{-1}s^{-1}$$
$$k_{a,TA} = 3 * 10^6\ M^{-1}s^{-1}$$

The following definitions were used:

| | |
|---|---|
| $\tau_{off,TA-mediated}$: | Average time no Imager probe is bound to a docking site of a DNA origami designed to bind Imager probes via a Transient Adapter |
| $\tau_{off,direct}$: | Average time no Imager probe is bound to a docking site of a DNA origami designed to directly bind Imager probes |
| $\tau_{off,TA}$: | Average time no Transient Adapter is bound to a docking site of a corresponding DNA origami |
| $\tau_{on,TA}$: | Average time a Transient Adapter is bound to a docking site of a corresponding DNA origami |
| $\tau_{off,Im}$: | Average time no Imager probe is bound to a specific docking site in the absence of Transient Adapters |
| $\tau_{on,Im}$: | Average time an Imager probe is bound to its complimentary sequence either as part of a Transient Adapter, or as a direct docking site on a corresponding DNA origami |
| $k_{a,eff,TA-mediated}$: | Effective association rate constant of Imager probe binding to DNA origami docking site via a Transient Adapter, including corrections for Duty Cycle and competition from binding to Transient Adapters in solution |
| $k_{a,eff,direct}$: | Effective association rate constant of Imager probe binding directly to a DNA origami featuring a suitable to docking site in the presence of Transient Adapters, including competition from binding to Transient Adapters in solution |
| $k_{a,Im}$: | Association rate constant of Imager probes binding to their complimentary sequence |
| $k_{a,TA}$: | Association rate constant of Transient Adapter binding to a docking site |
| $D$: | Duty Cycle, i.e. fraction of time a docking site is occupied by a Transient Adapter |
| $c_{Im}$: | Molar concentration of Imager probe initially added to imaging buffer |
| $c_{Im,free}$: | Molar concentration of Imager probes not bound to Transient Adapters in solution |
| $c_{TA}$: | Molar concentration of Transient Adapter |
| $f$: | Unbound fraction of Imager probes |
| $K_D$: | Equilibrium dissociation constant between Imager probes and Transient Adapters |

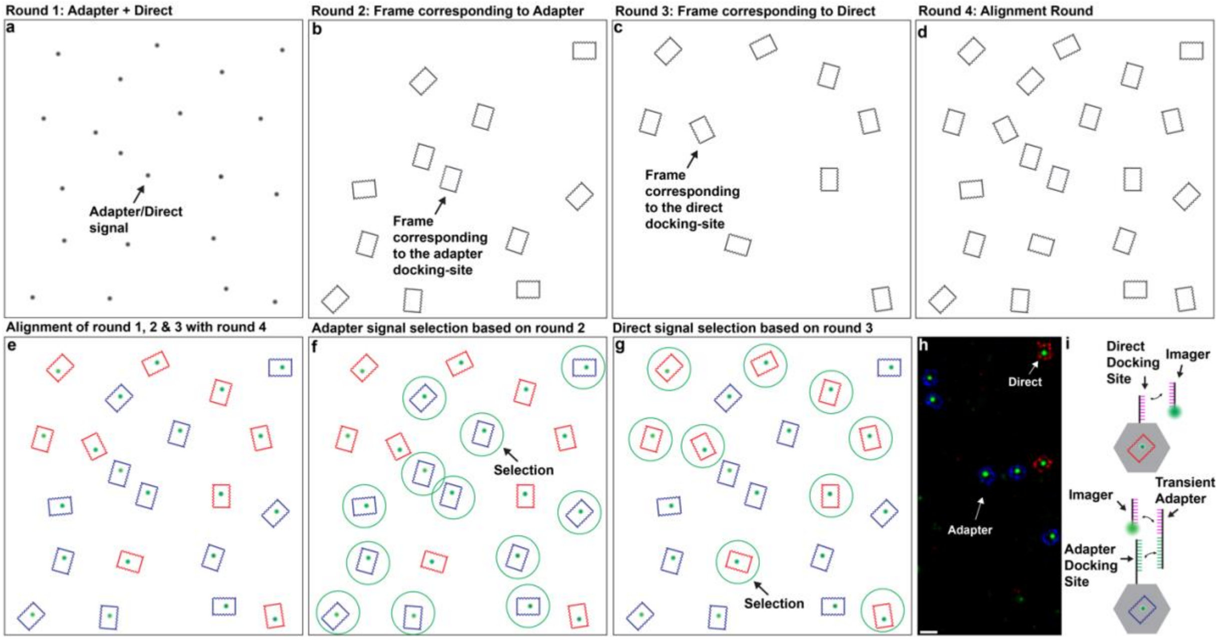

**Figure S1 – Experimental Workflow for Kinetics Measurements using DNA origami.** Two distinct DNA origami species are used simultaneously. The first DNA origami species features a single docking site for Imager probe binding via a Transient Adapter strand and orthogonal docking sites arranged in a frame pattern (**Figure S2**). The second DNA origami species features a single docking site for direct Imager probe binding and the same frame pattern but with a docking sequence orthogonal to that used in the first DNA origami species. (**a**) In the first imaging round, the single docking sites on both DNA origami species are imaged; the same Imager probe can bind to the first DNA origami species via a Transient Adapter and directly to the second DNA origami species. (**b**) In the second imaging round, the frame of the first DNA origami species is imaged. (**c**) In the third imaging round, the frame of the second DNA origami species is imaged. (**d**) In the final imaging round, both frames of both DNA origami species are imaged. (**e**) After applying standard single-molecule localization-based super-resolution microscopy post-processing techniques (i.e., localization fitting and drift correction), the first three imaging rounds are aligned with the last round. (**f, g**) Using the frame images of the two DNA origami species, single docking sites are identified and binding kinetics are extracted for kinetics analysis for Transient Adapter-mediated (f) and direct (g) binding. (**h**) Cut-out of exemplary field of view. Colors are assigned based on in which imaging round the signal was recorded. (**i**) Schematic representation of direct and Transient Adapter-mediated Imager binding. Scale bar 100 nm.

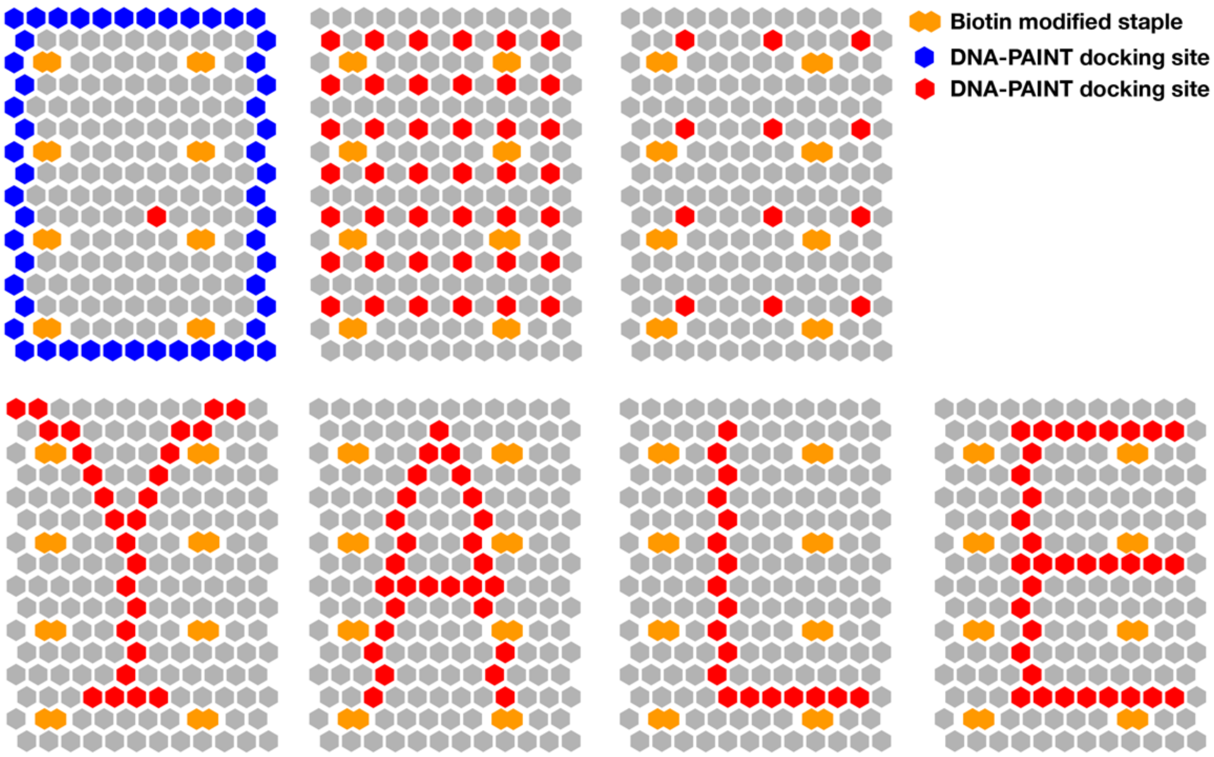

**Figure S2 – Used DNA origami designs.** Schematic representation of all DNA origami designs used in this study. The hexagons represent 3'-staple positions. Blue and red hexagons are representing two different staples extended with docking sites for transient binding of either Imager probes or Transient Adapters. The orange hexagons depict staples extended with a biotin modification for immobilization on the cover slip surface.

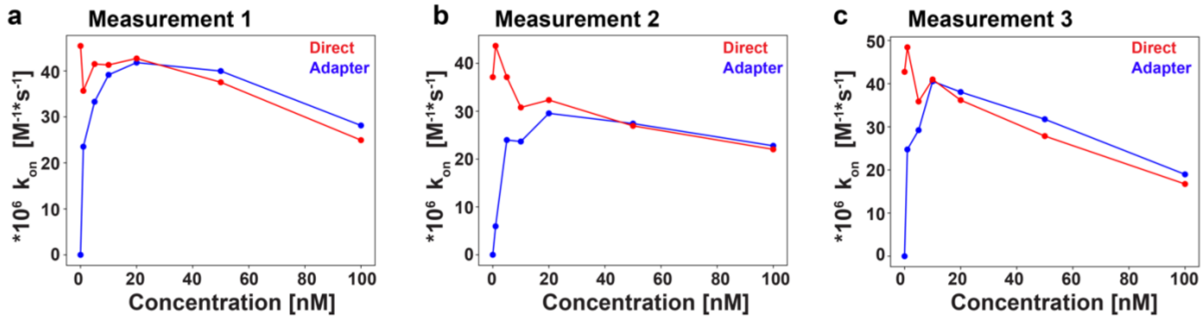

**Figure S3 – Three independent replicates of kinetics experiments measuring effective association rates of Imager probes binding directly or via Transient Adapters to DNA origami docking sites as a function of Transient Adapter concentration.** The A5-5xR2 Transient Adapter was used following the workflow described in Figure S1. The data points shown in **Figure 1d** are the averaged values from these three experiments.

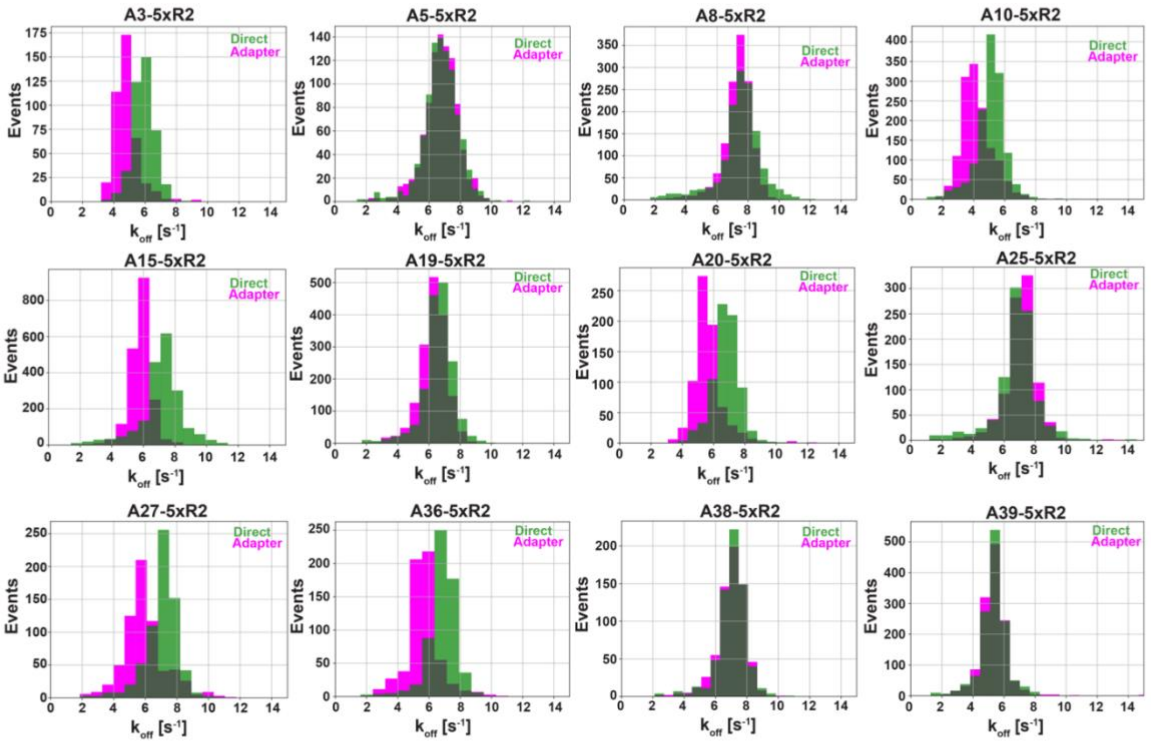

**Figure S4 – Measured dissociation rates for Transient Adapters for Imager R2 (speed Imager).** Dissociation rates for direct and Transient Adapter-mediated binding for all 12 Transient Adapters with the 5xR2 Imager docking site sequence using the workflow of Figure S1.

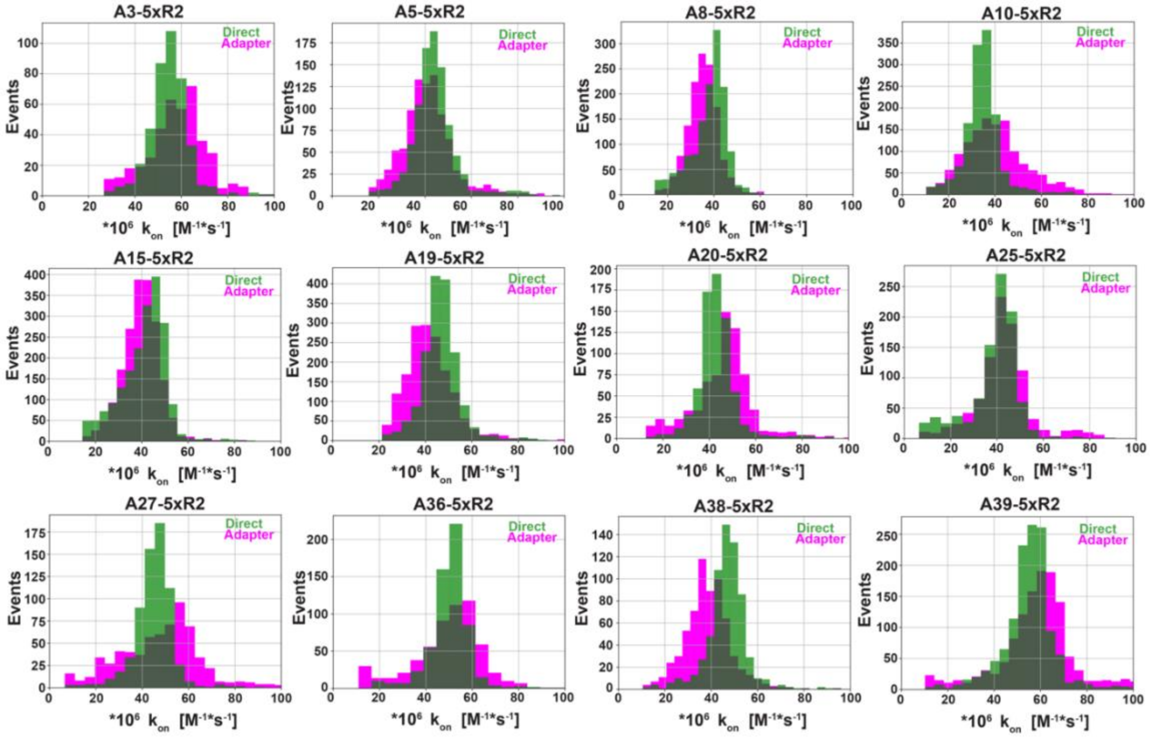

**Figure S5 – Measured association rates for Transient Adapters for Imager R2 (speed Imager).** Association rates for direct and Transient Adapter-mediated binding for all 12 Transient Adapters with the 5xR2 Imager docking site sequence using the workflow of Figure S1.

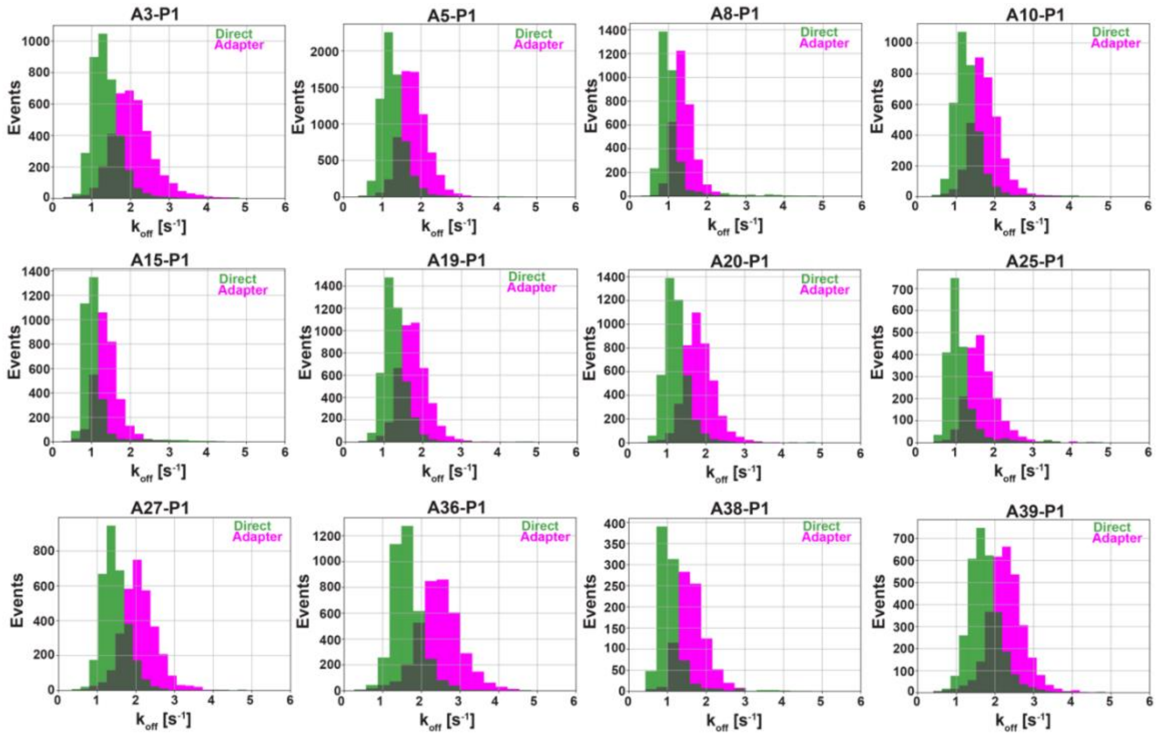

**Figure S6 – Measured dissociation rates for Transient Adapters for Imager P1 (classical Imager).** Dissociation rates for direct and Transient Adapter-mediated binding for all 12 Transient Adapters with the P1 Imager docking site sequence using the workflow of Figure S1.

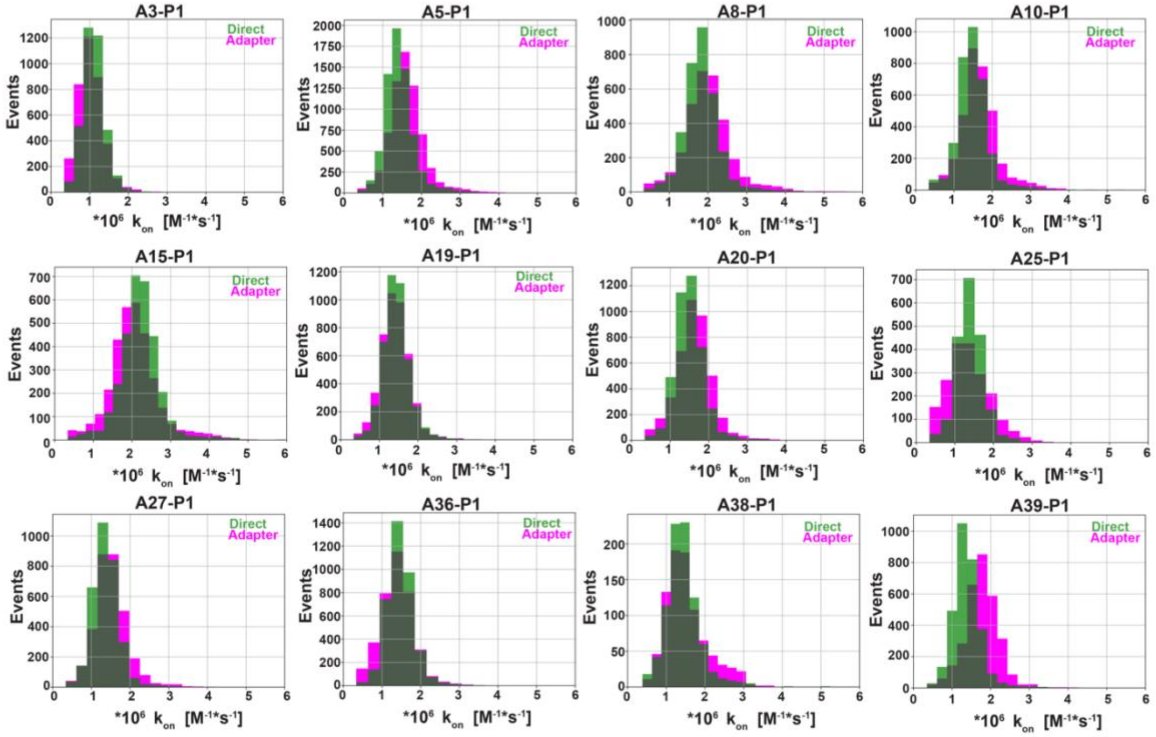

**Figure S7 – Measured association rates for Transient Adapters for Imager P1 (classical Imager).** Association rates for direct and Transient Adapter-mediated binding for all 12 Transient Adapters with the P1 Imager docking site sequence using the workflow of Figure S1.

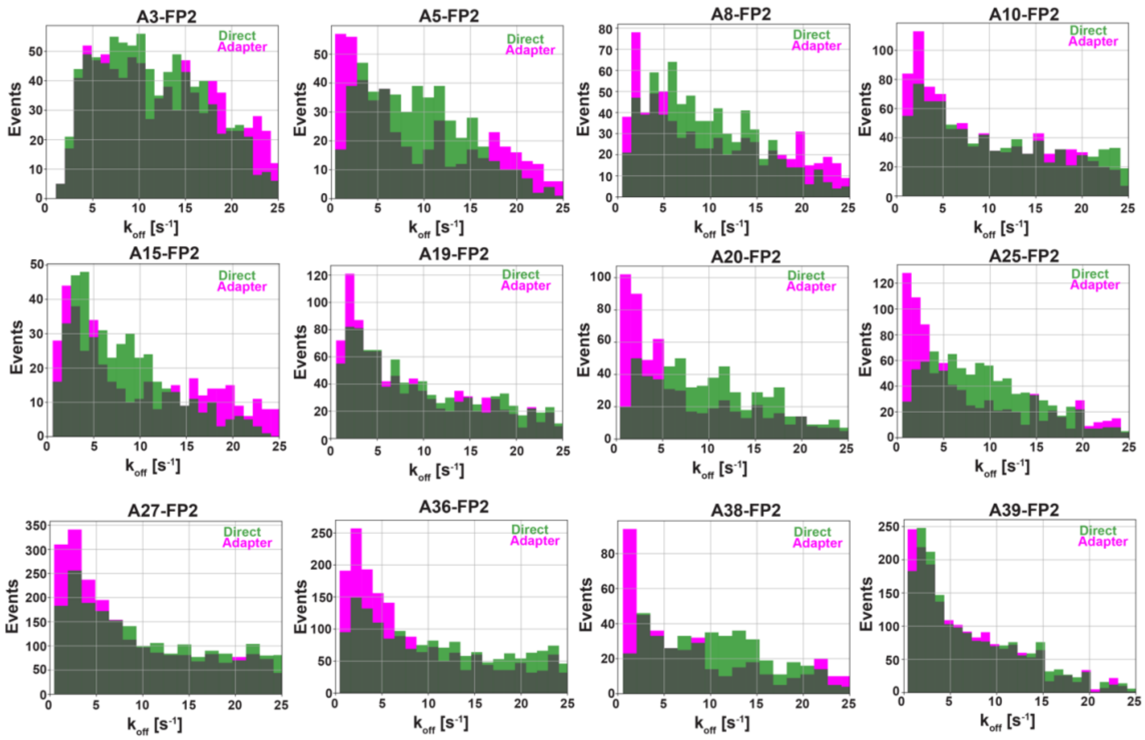

**Figure S8 – Measured dissociation rates for Transient Adapters for Imager FP2 (fluorogenic Imager).** Dissociation rates for direct and Transient Adapter-mediated binding for all 12 Transient Adapters with the FP2 Imager docking site sequence using the workflow of Figure S1.

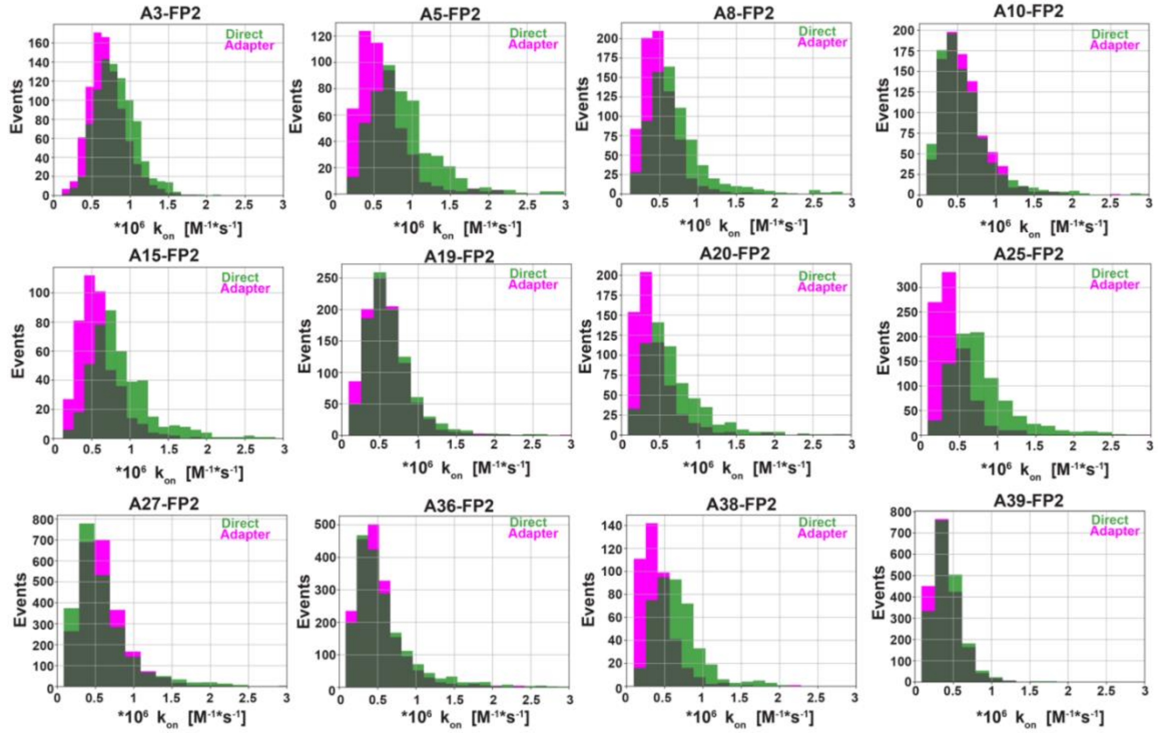

**Figure S9 – Measured association rates for Transient Adapters for Imager FP2 (fluorogenic Imager).** Association rates for direct and Transient Adapter-mediated binding for all 12 Transient Adapters with the FP2 Imager docking site sequence using the workflow of Figure S1.

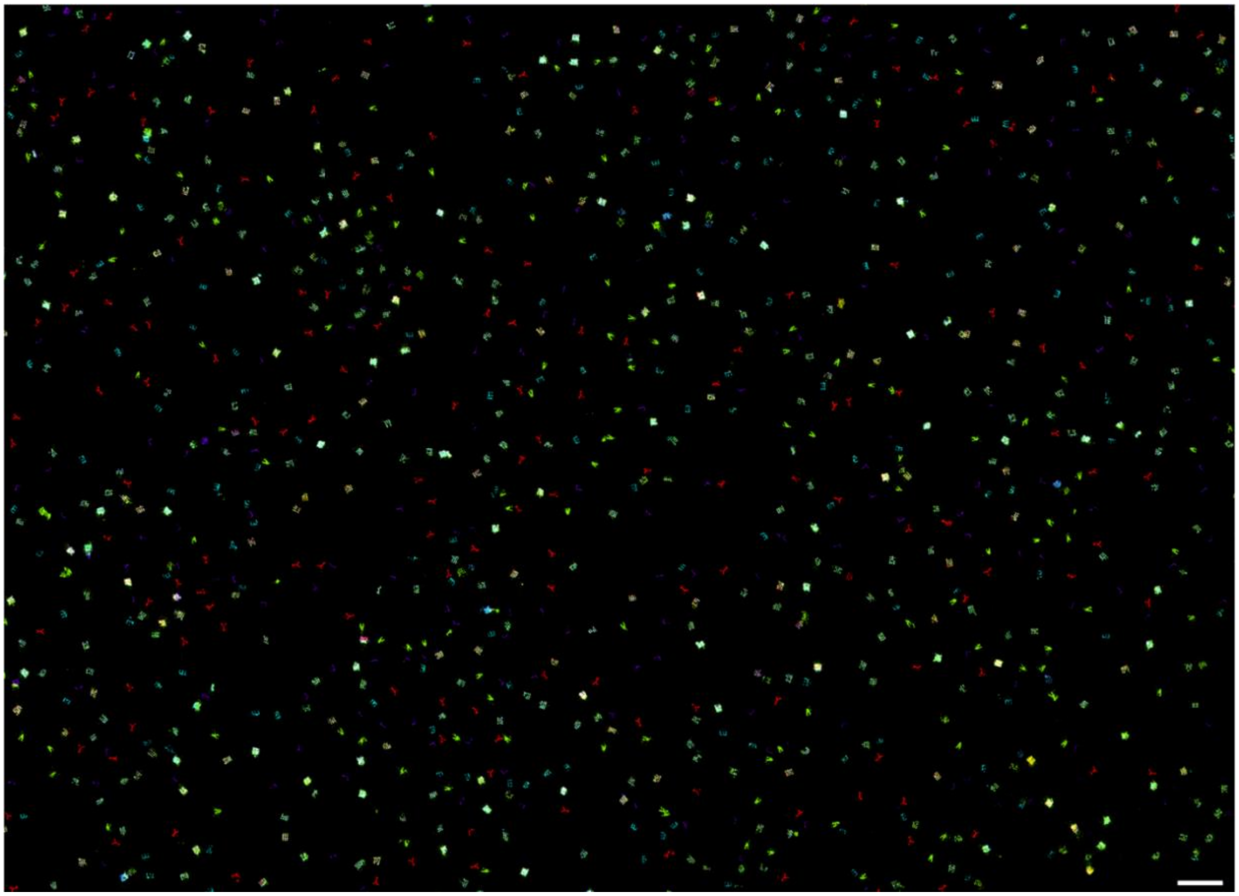

**Figure S10 – Representative field of view of the 4-plex DNA origami letters experiment (Figure 1e).** Imaged DNA origami nano structures: Round 1: 'Y' & 20-nm grid & 10-nm grid (red); Round 2: 'A' & 20-nm grid & 10-nm grid (green), Round 3: 'L' & 20-nm grid & 10-nm grid (magenta); Round 4: 'E' & 20-nm grid & 10-nm grid (cyan). The 20-nm and 10-nm grids were imaged in each round and used for drift correction and alignment of the individual rounds. Scale bar 500 nm.

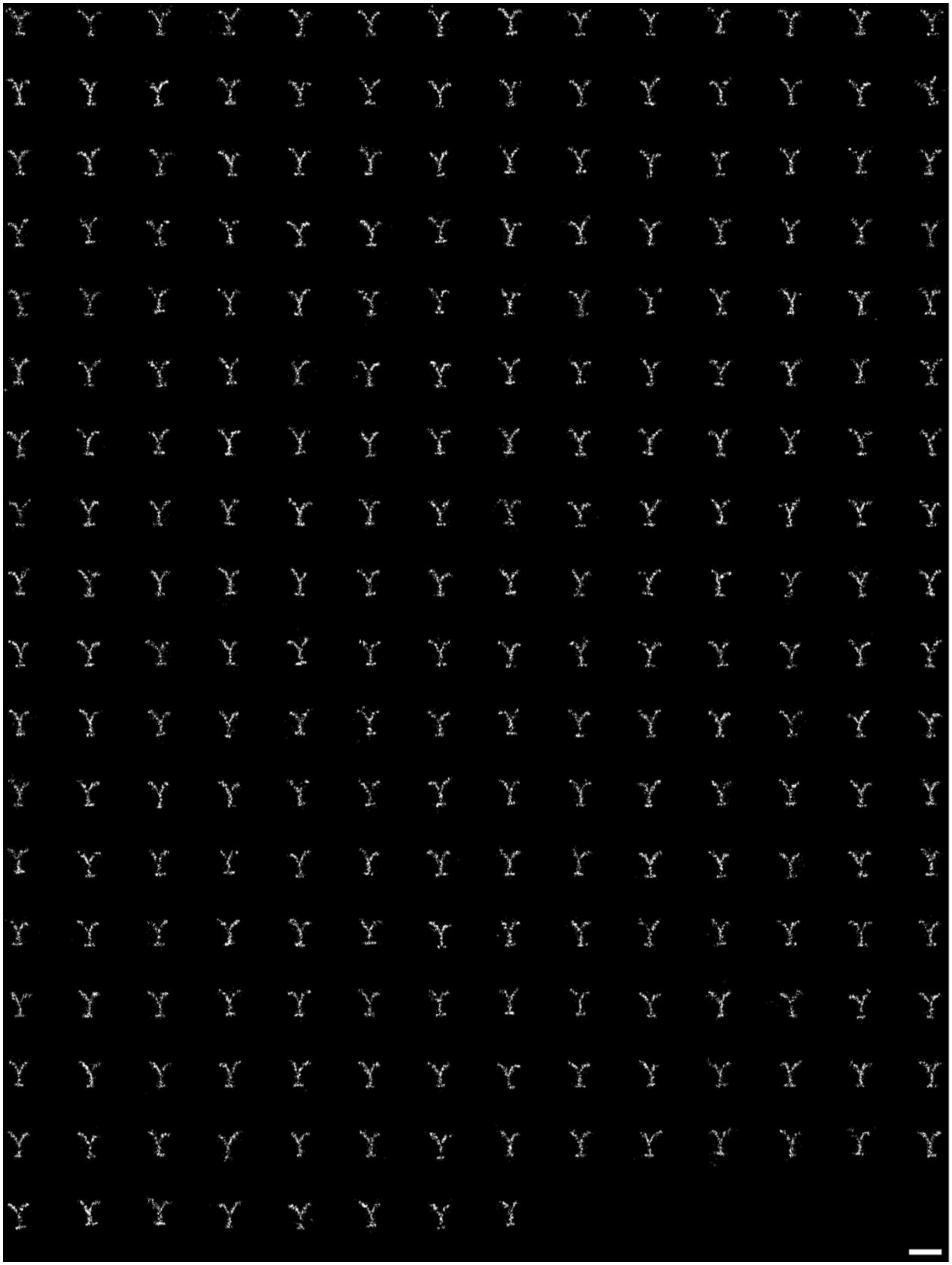

**Figure S11 – Single DNA origami structures displaying the letter 'Y'.** Images of the 246 single structures used to produce the averaged image depicted in **Figure 1e**. Scale bar 100 nm.

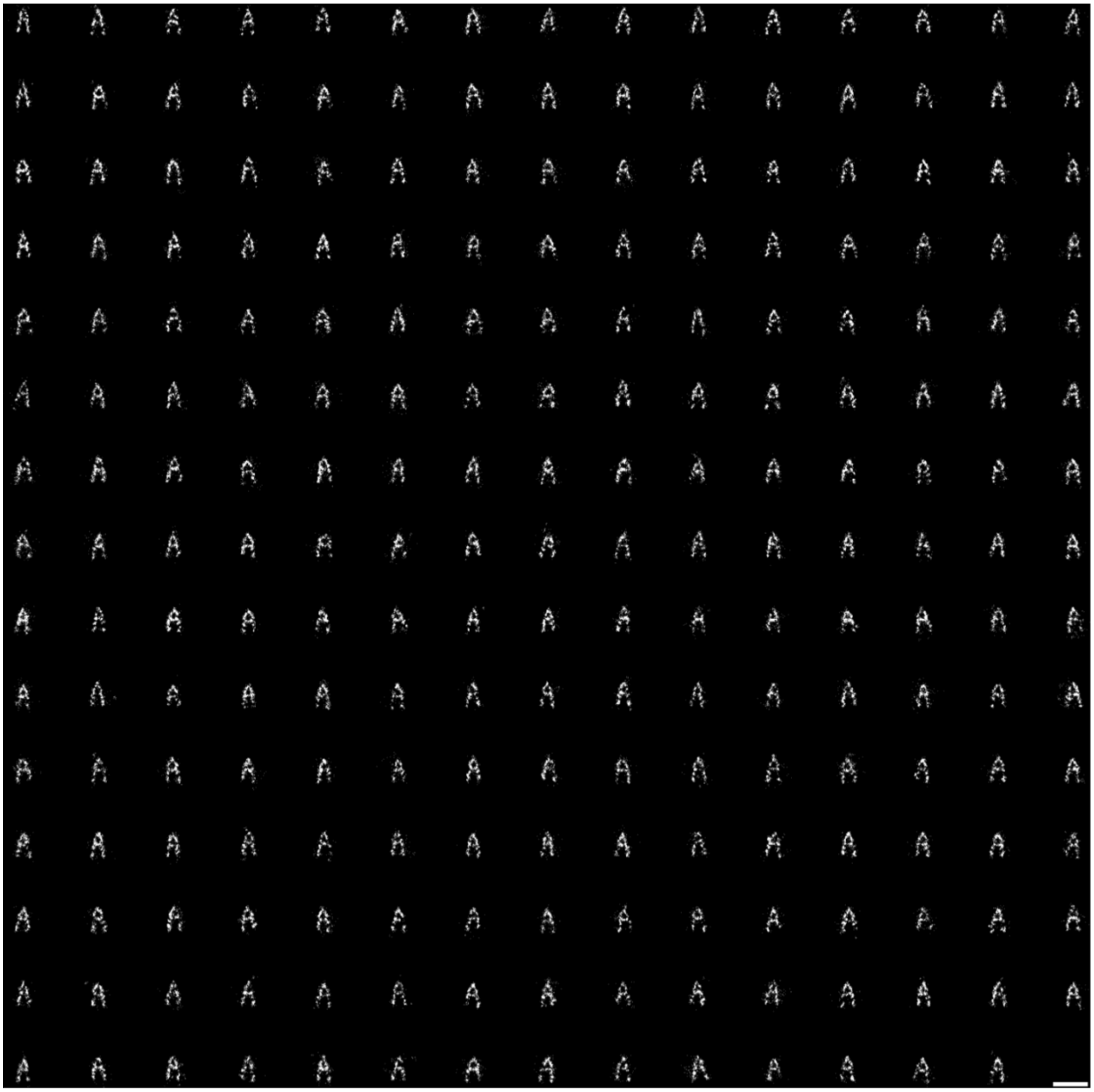

**Figure S12 – Single DNA origami structures displaying the letter 'A'.** Images of 224 single structures used to produce the averaged image depicted in **Figure 1e**. Scale bar 100 nm.

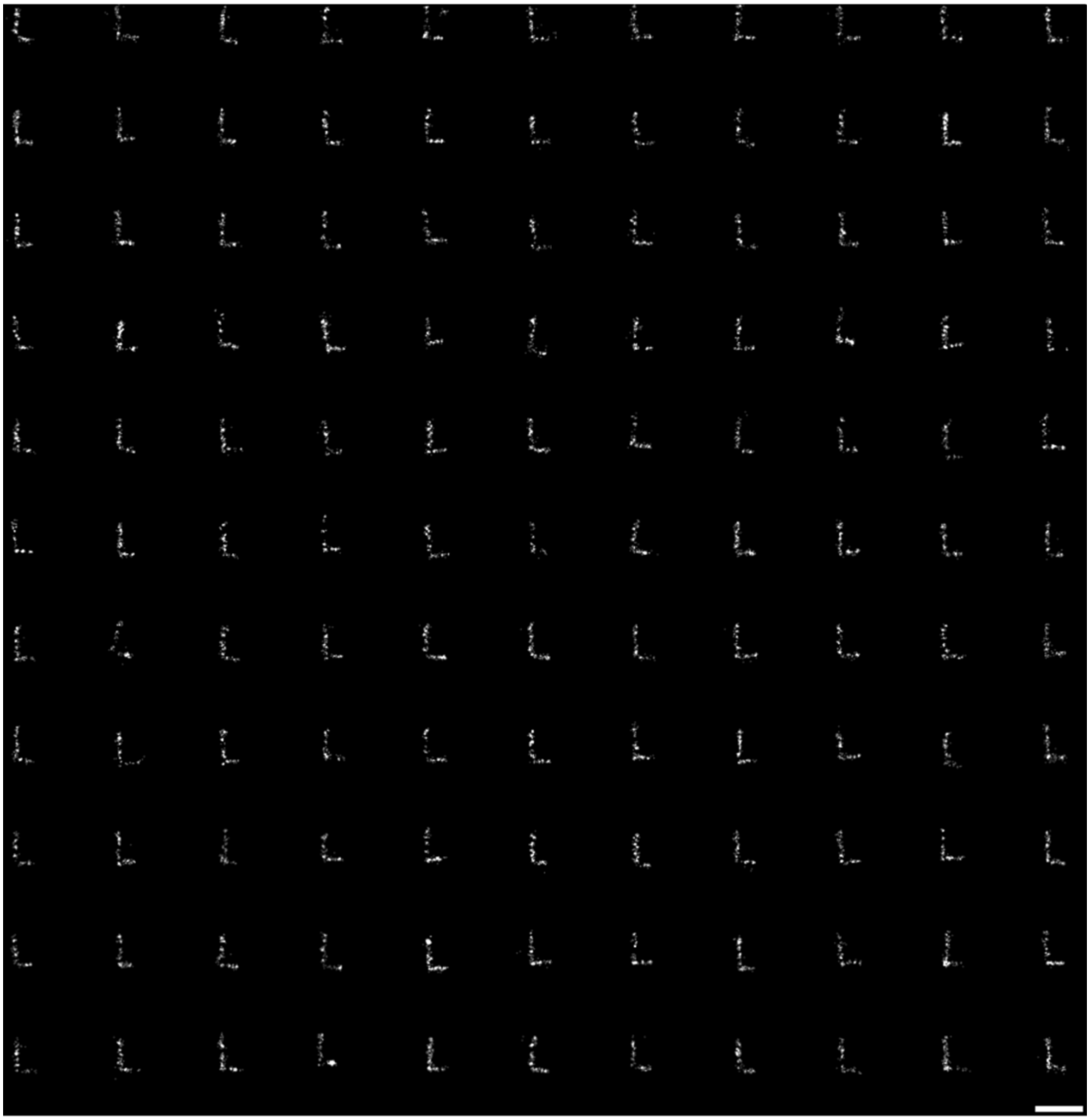

**Figure S13 – Single DNA origami structures displaying the letter 'L'.** Images of 121 single structures used to produce the averaged image depicted in **Figure 1e**. Scale bar 100 nm.

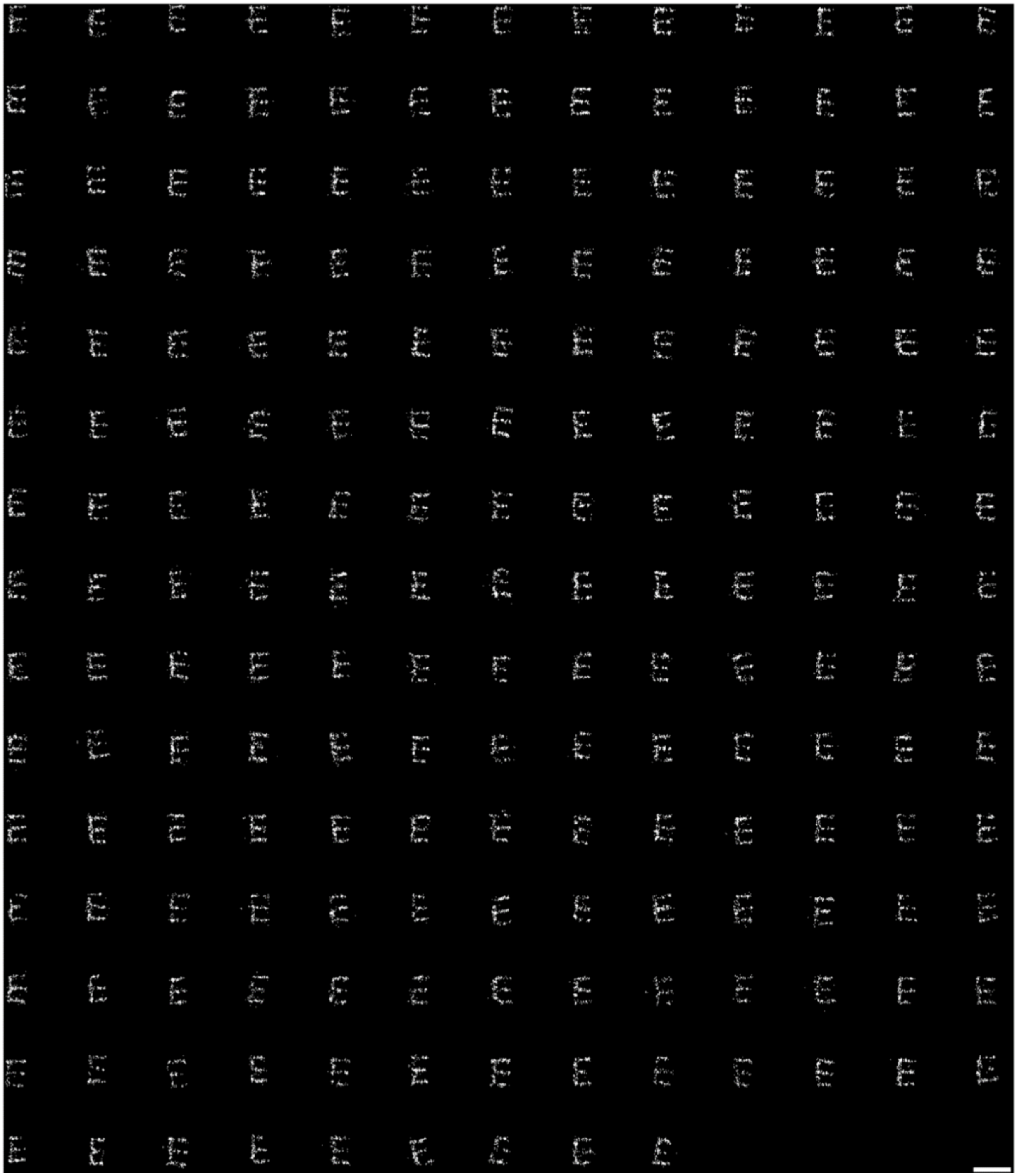

**Figure S14 – Single DNA origami structures displaying the letter 'E'.** Images of 191 single structures used to produce the averaged image depicted in **Figure 1e**. Scale bar 100 nm.

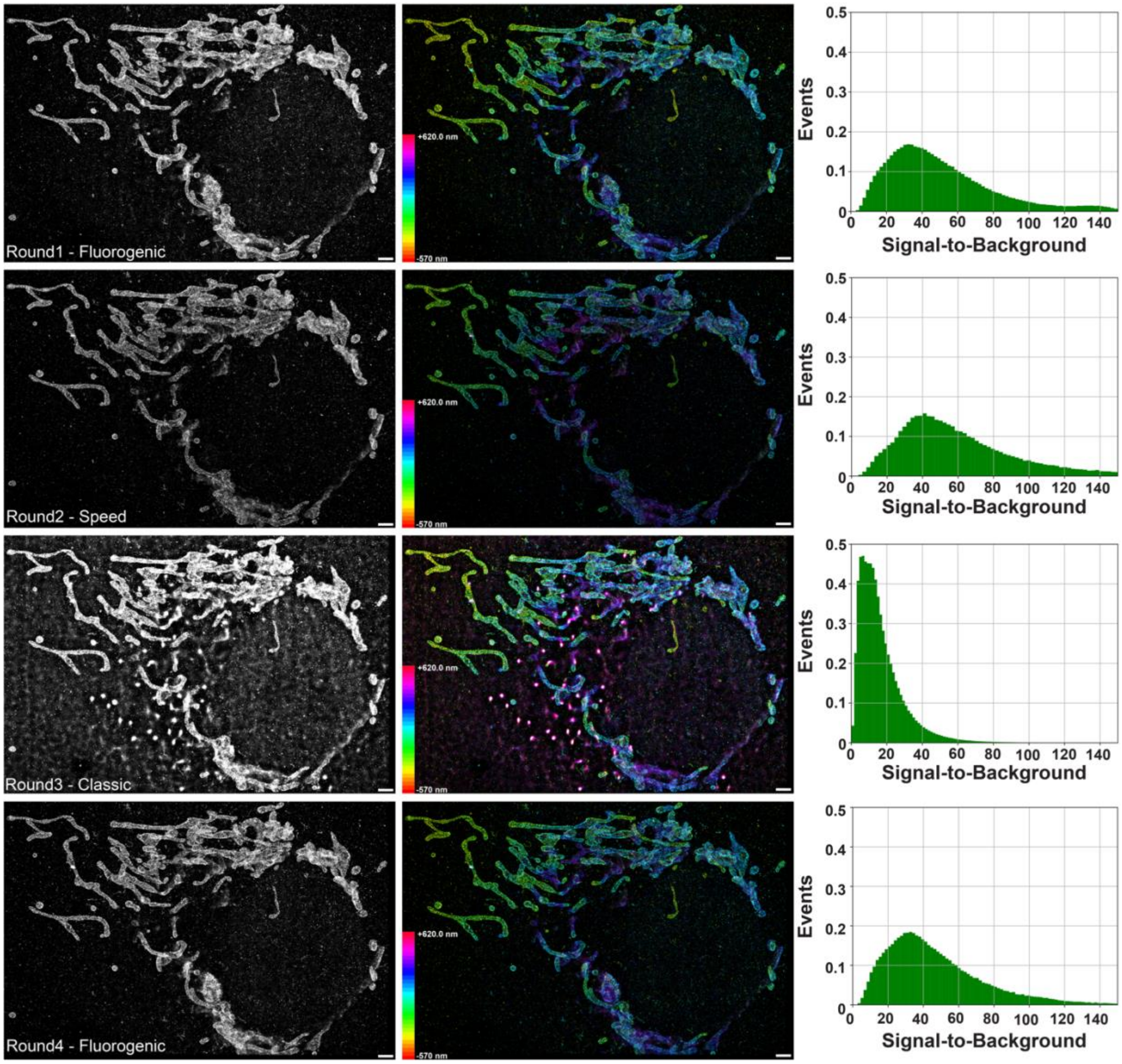

**Figure S15 – Direct comparison of imaging performance for fluorogenic, speed, and classical Imagers**. Mitochondria were immunolabeled with a primary antibody against Tom20. Secondary antibodies were conjugated with a docking site for the A3 Transient Adapter. In the first round of the sequential imaging experiment, the fluorogenic Imager FP2 and a Transient Adapter to FP2 were used. After data acquisition, both were washed out and replaced by the speed Imager R2 and a Transient Adapter with a 5xR2 motif. After a second round of data acquisition the Imager and Transient Adapter were washed out once more and replaced by the classical Imager P1 and a corresponding Transient Adapter. A third round of data acquisition was followed by another round of washes and reintroduction of fluorogenic Imager FP2 and a Transient Adapter to FP2 to verify that sample degradation was negligible. A comparison shows a substantial improvement of the signal-to-background ratio using the fluorogenic and the speed Imagers (SNR $\approx$ 40) over the classical Imager P1 (SNR $\approx$ 8). Scale bar 2 µm.

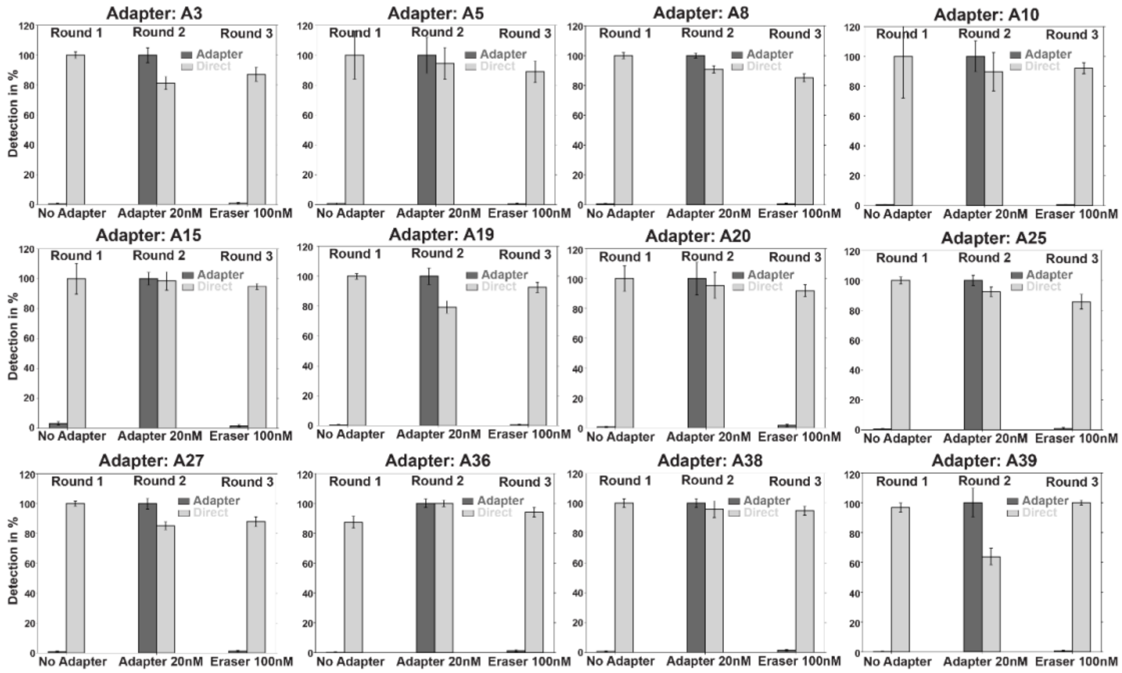

**Figure S16 – Quantification of erasing efficiency for all 12 Transient Adapter sequences.** A modified workflow of Figure S1 was used: in the first round, only the Imager but no Transient Adapter is introduced. In the second round, both the Transient Adapter (20 nM) and Imager are used. During the final round of imaging, the solution is replaced (no washes) by the corresponding Eraser strand (100 nM) and the same Imager. After waiting for 3 min, the third round of imaging is conducted. The analyzed single docking sites are divided into ten random groups, from which the mean and standard deviation are calculated. All values are normalized.

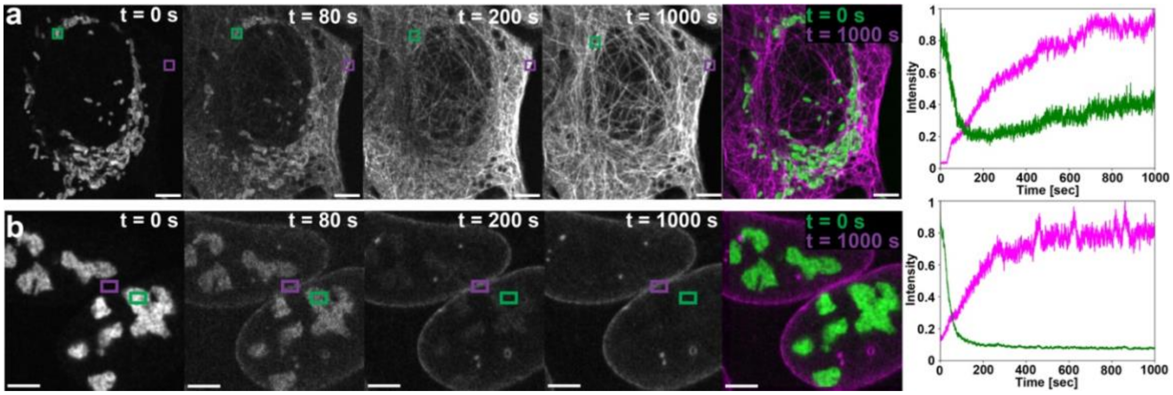

**Figure S17 – Observation of molecular target switch dynamics. (a)** U-2 OS cell labeled with antibodies against mitochondrial outer membrane protein Tom20 and α-tubulin. Before the shown time course, the sample was in a medium containing a Transient Adapter directing Imager R2 to mitochondria. At the start of image acquisition, a corresponding Eraser and a new Transient Adapter which directs the Imager to α-tubulin are added. The Eraser rapidly erases the mitochondria signal ($\tau_{1/2} \approx 60$ s) as the α-tubulin signal, mediated by the new Transient Adapter, increases ($\tau_{1/2} \approx 200$ s). **(b)** Equivalent experiment using antibodies against NPM1 (nucleolus) and LaminB1 (nuclear envelope). Data was acquired with a spinning disk microscope and is diffraction limited. Scale bar 5 µm.

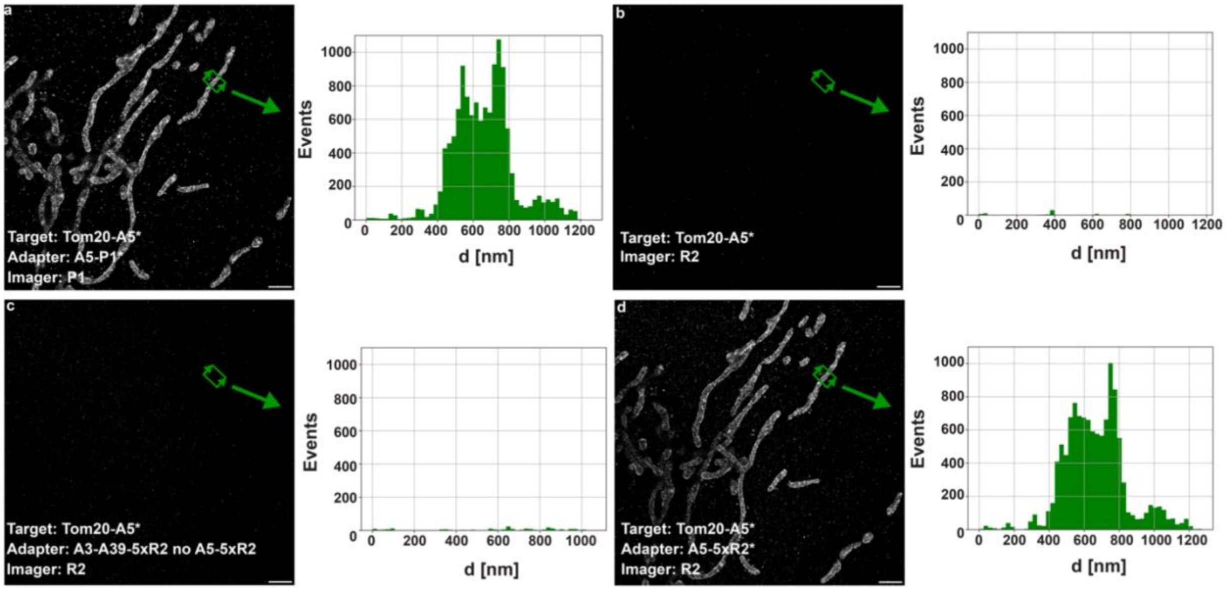

**Figure S18 – Evaluation of non-specific binding of the Transient Adapter and speed Imager system.** The mitochondrial protein Tom20 in U-2 OS cells is immunolabeled with a A5 docking site using primary and secondary antibodies. (**a**) In the first round, to identify a field of view, the classical P1 Imager and the Transient Adapter directing P1 to A5 are introduced. The histogram displays a one-dimensional projection of localizations along the arrows at the highlighted region of interest and shows robust signal. (**b**) After two brief washes, the R2 Imager is introduced. Since the corresponding Transient Adapter is absent, no specific binding is expected, and the corresponding histogram shows very few localization events. (**c**) In the third round of imaging, all Transient Adapters except for the correct one (A5-5xR2) are added to the R2 Imager. As none of the introduced Transient Adapters should be able to interact with the docking site, no specific interaction or downstream sampling is anticipated. This is confirmed by the histogram which again shows only very few localization events. (**d**) In the fourth round of imaging, the matching Transient Adapter (A5-5xR2) is introduced alongside the Imager R2. As expected, the histogram shows robust signal, comparable to (a). Scale bar 2 µm.

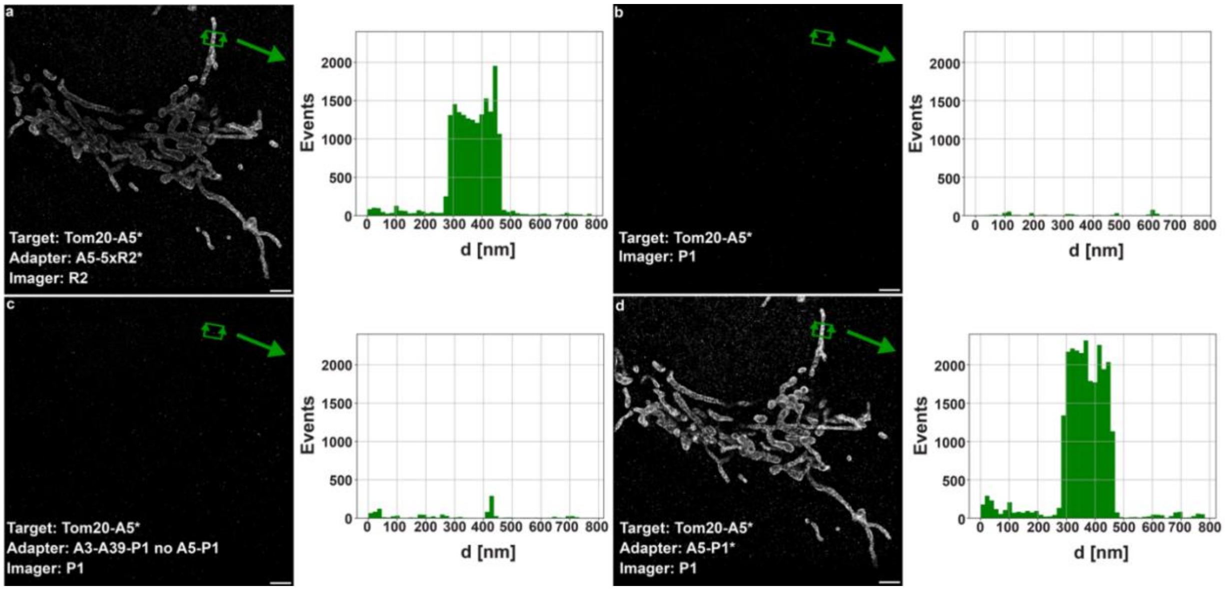

**Figure S19 – Evaluation of non-specific binding of the Transient Adapter and classical Imager system.** The mitochondrial protein Tom20 in U-2 OS cells is immunolabeled with a A5 docking site using primary and secondary antibodies. (**a**) In the first round, to identify a field of view, the R2 speed Imager and the Transient Adapter directing R2 to A5 are introduced. The histogram displays a one-dimensional projection of localizations along the arrows at the highlighted region of interest and shows robust signal. (**b**) After two brief washes, the P1 Imager is introduced. Since the corresponding Transient Adapter is absent, no specific binding is expected, and the corresponding histogram shows very few localization events. (**c**) In the third round of imaging, all Transient Adapters except for the correct one (A5-P1) are added to the P1 Imager. As none of the introduced Transient Adapters should be able to interact with the docking site, no specific interaction or downstream sampling is anticipated. This is confirmed by the histogram which again shows only very few localization events. (**d**) In the fourth round of imaging, the matching Transient Adapter (A5-P1) is introduced alongside the Imager P1. As expected, the histogram shows robust signal, comparable to (a). Scale bar 2 µm.

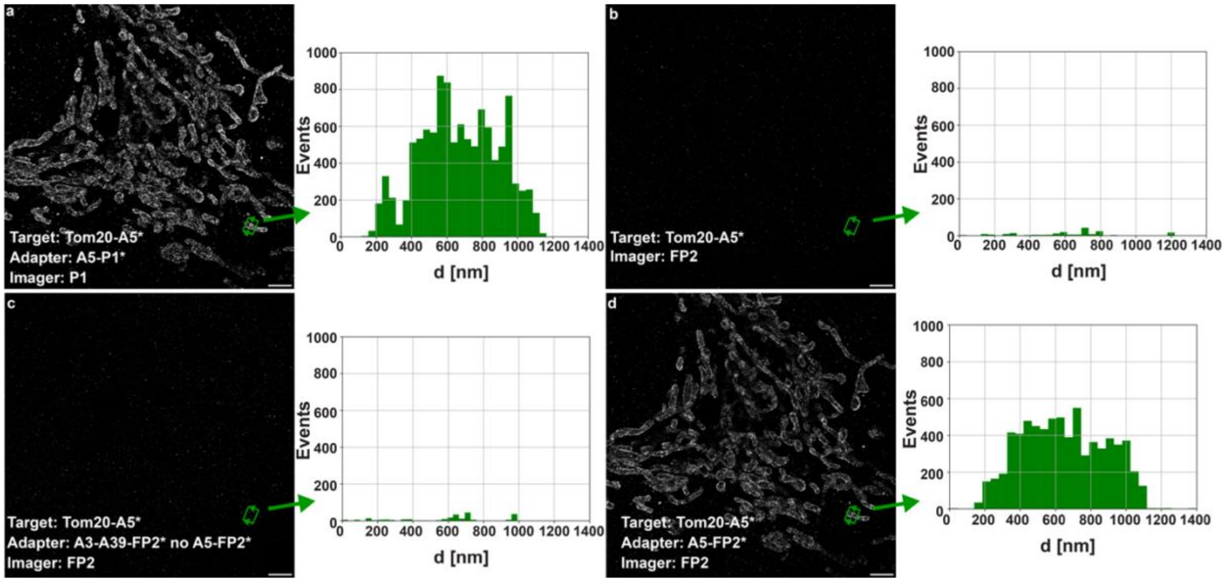

**Figure S20 – Evaluation of non-specific binding of the Transient Adapter and fluorogenic Imager system.** The mitochondrial protein Tom20 in U-2 OS cells is immunolabeled with a A5 docking site using primary and secondary antibodies. (**a**) In the first round, to identify a field of view, the classical P1 Imager and the Transient Adapter directing P1 to A5 are introduced. The histogram displays a one-dimensional projection of localizations along the arrows at the highlighted region of interest and shows robust signal. (**b**) After two brief washes, the FP2 Imager is introduced. Since the corresponding Transient Adapter is absent, no specific binding is expected, and the corresponding histogram shows very few localization events. (**c**) In the third round of imaging, all Transient Adapters except for the correct one (A5-FP2) are added to the FP2 Imager. As none of the introduced Transient Adapters should be able to interact with the docking site, no specific interaction or downstream sampling is anticipated. This is confirmed by the histogram which again shows only very few localization events. (**d**) In the fourth round of imaging, the matching Transient Adapter (A5-FP2) is introduced alongside the Imager FP2. As expected, the histogram shows robust signal, comparable to (a). Scale bar 2 µm.

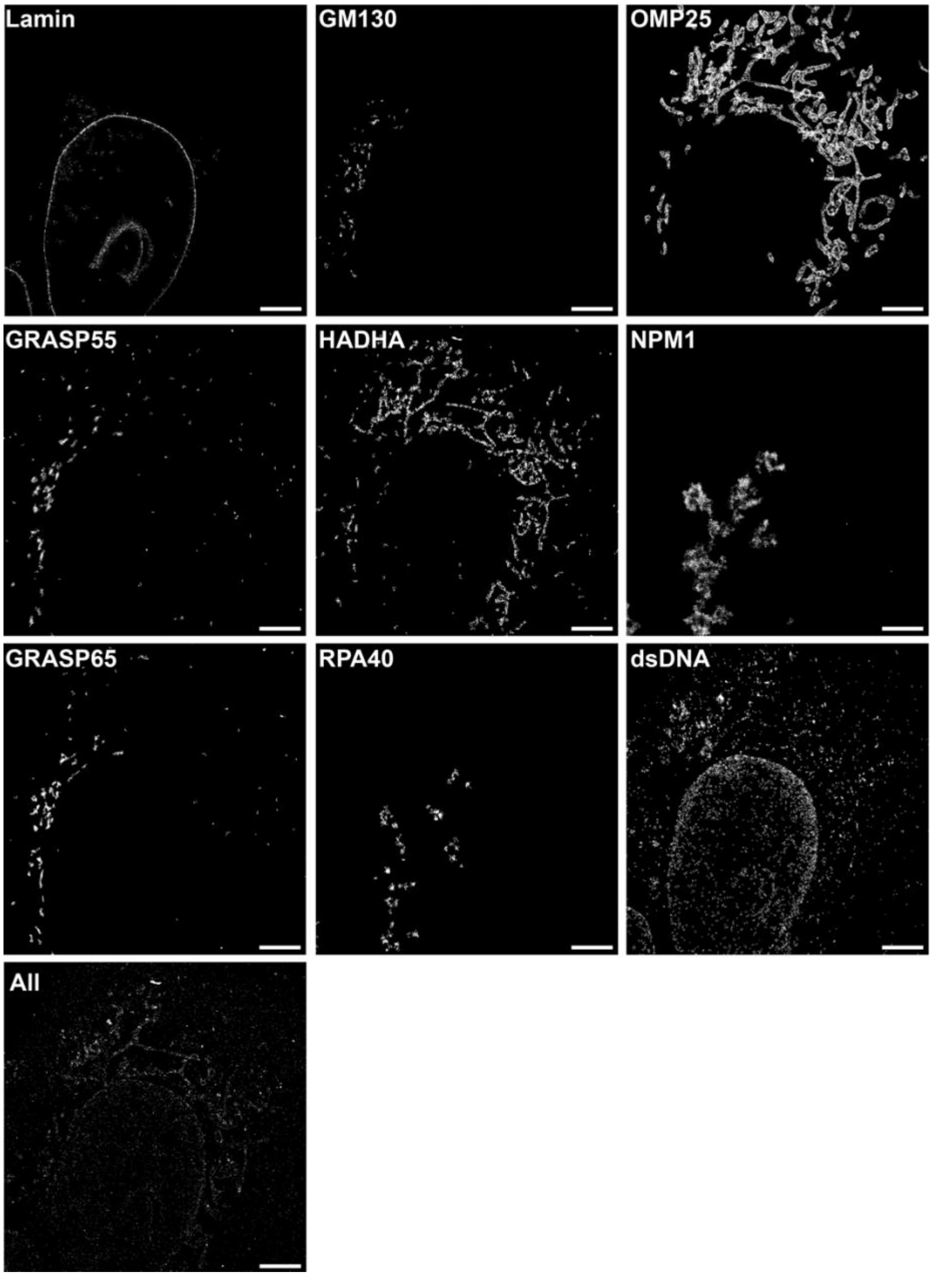

**Figure S21 – Images of the individual imaging rounds of Figure 3.** Additionally, to the single-target imaging rounds, for alignment purposes, the sample is imaged an additional time with all targets simultaneously. Scale bars: 5 µm.

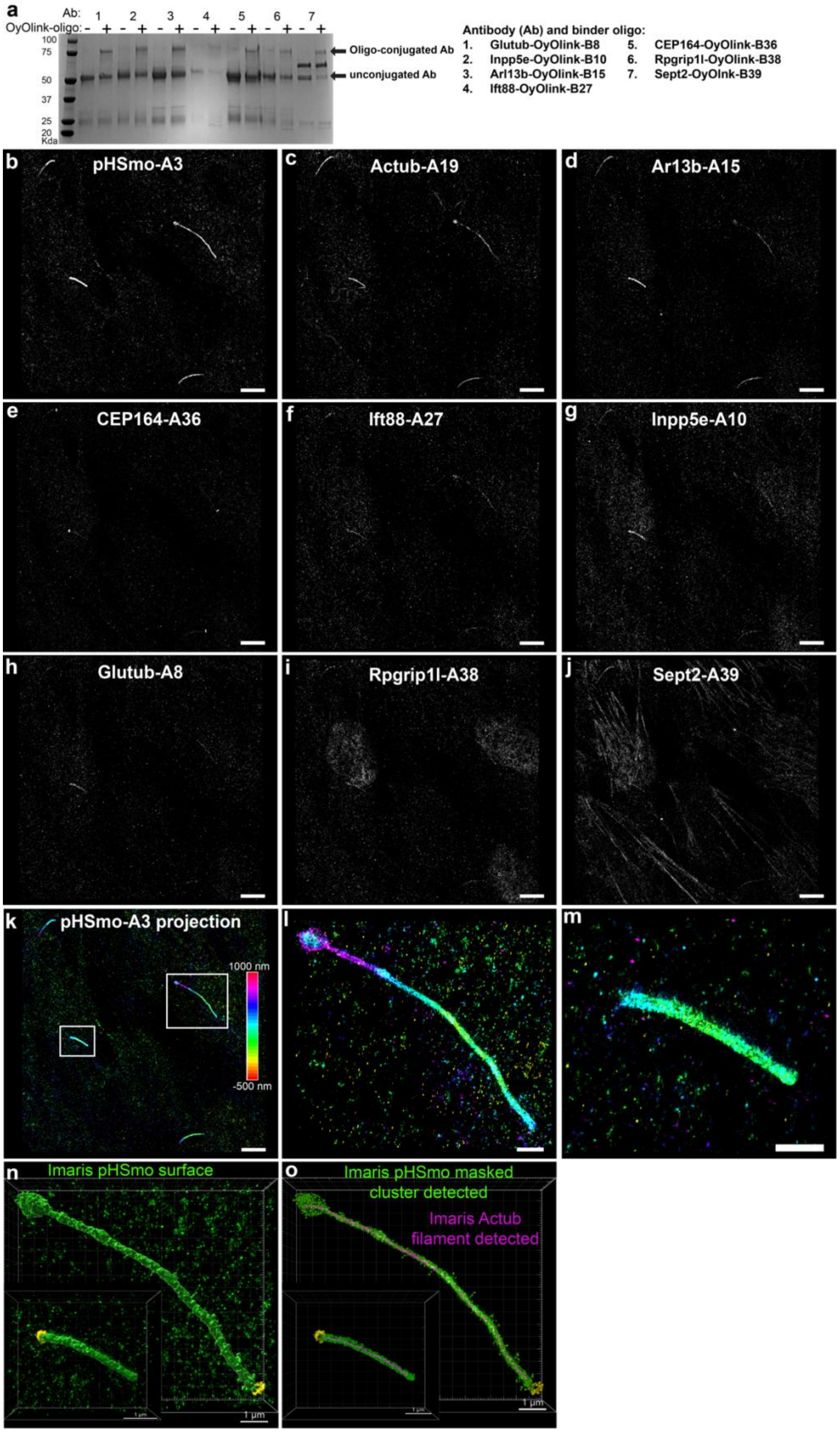

a
Ab: 1 2 3 4 5 6 7
OyOlink-oligo: - + - + - + - + - + - + - +
100
75
50
37
25
20
Kda
Oligo-conjugated Ab
unconjugated Ab
Antibody (Ab) and binder oligo:
1. Glutub-OyOlink-B8
2. Inpp5e-OyOlink-B10
3. Arl13b-OyOlink-B15
4. Ift88-OyOlink-B27
5. CEP164-OyOlink-B36
6. Rpgrip1l-OyOlink-B38
7. Sept2-OyOlnk-B39
b pHSmo-A3
c Actub-A19
d Ar13b-A15
e CEP164-A36
f Ift88-A27
g Inpp5e-A10
h Glutub-A8
i Rpgrip1l-A38
j Sept2-A39
k pHSmo-A3 projection
1000 nm
-500 nm
l
m
n Imaris pHSmo surface
1 μm
o Imaris pHSmo masked cluster detected
Imaris Actub filament detected
1 μm

**Figure S22 – Cilia-targeted antibody Light-Activated Site-specific Conjugation (LASIC) and images of the individual imaging rounds of Figure 4 with image segmentation.** (**a**) Coomassie blue-stained SDS-PAGE gel showing the direct conjugation of seven distinct OyOlink binder probe sequences (AlphaThera) to cilia-specific antibodies (Ab: 1-7; see legend in (a)). Conjugated heavy chain IgG transitions from 60 kDa (- lanes) to 75 kDa (+ lanes) upon incubation with OyOlink probe for 2 h under 365-nm UV light. (**b-j**) Images of the individual imaging rounds of Figure 4. (**k**) z-projection of the pH-Smo data, illustrating the full 1.5-µm z-range. (**l, m**) Zoomed-in views of the two analyzed cilia. (**n, o**) Examples of surface, clusters, and filaments generated with Imaris software for the pHSmo and ac-tub targets. Scale bars: 5 µm (b-k), 1 µm (l-o).

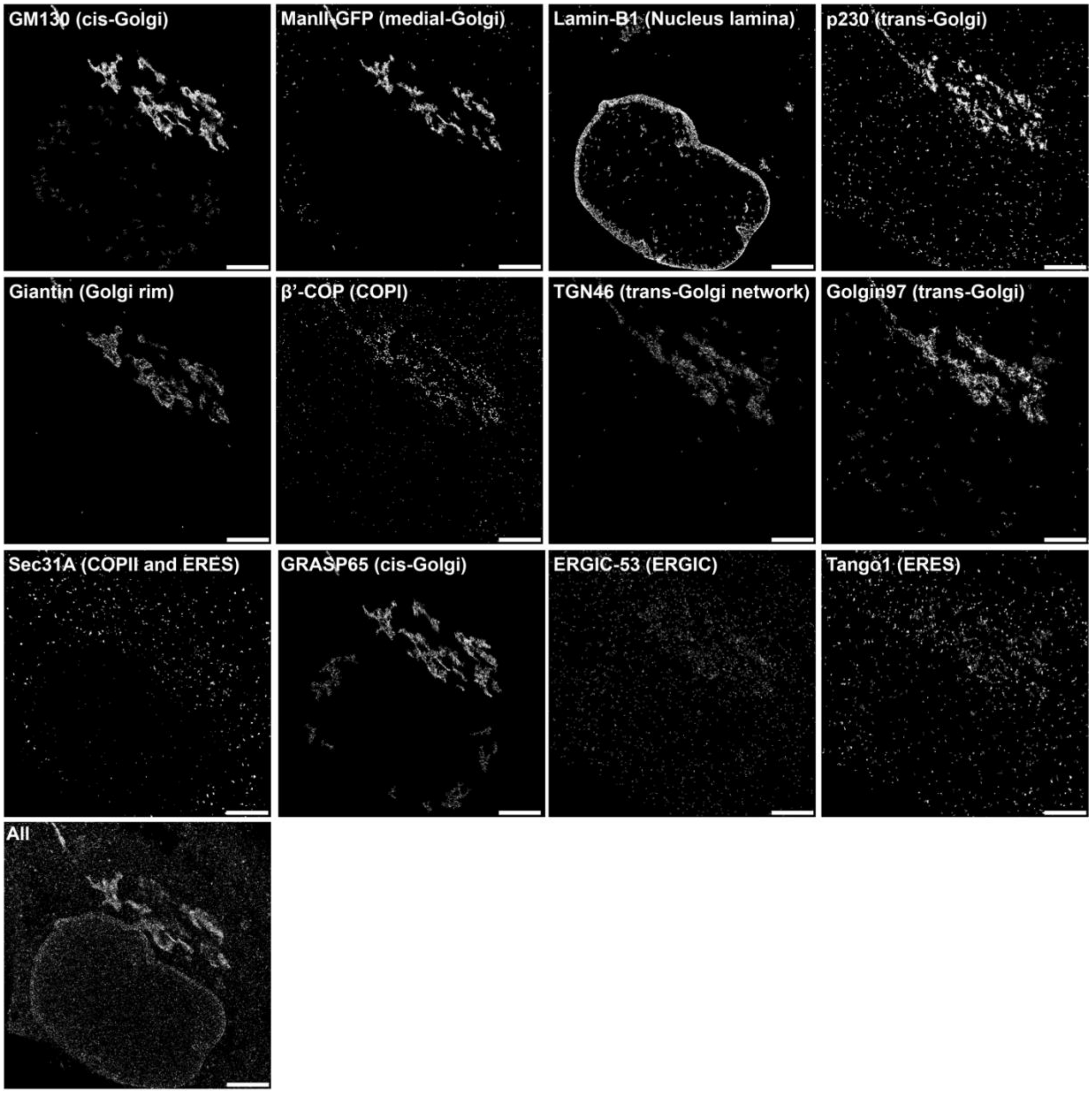

**Figure S23 – Images of the individual imaging rounds of Figures 5a-g.** Additionally to the single-target imaging rounds, for alignment purposes, the sample is imaged an additional time with all targets simultaneously. Scale bars: 5 µm.

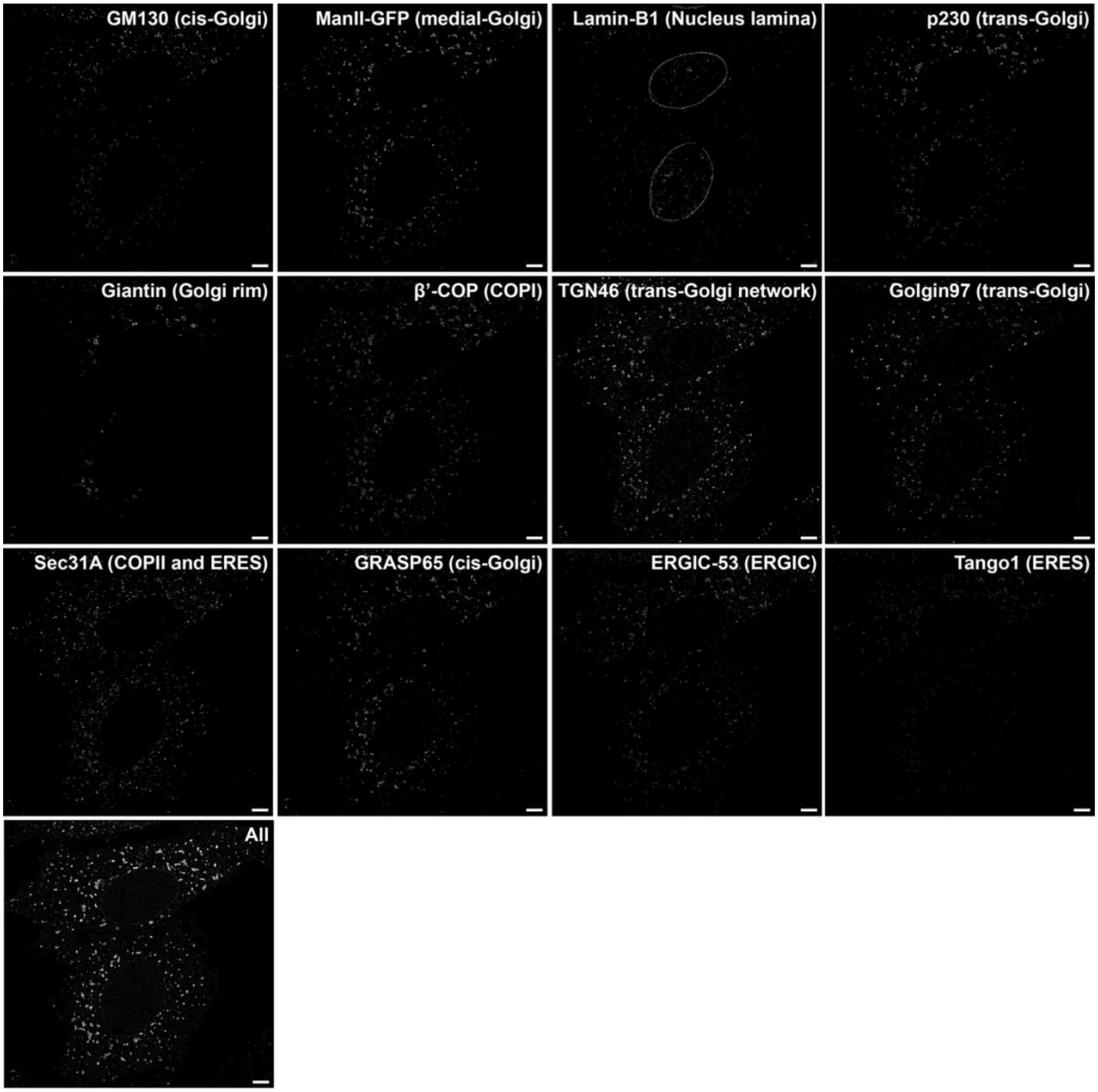

**Figure S24 – Images of the individual imaging rounds of Figures 5i-o.** Additionally to the single-target imaging rounds, for alignment purposes, the sample is imaged an additional time with all targets simultaneously. Scale bar: 5 µm.

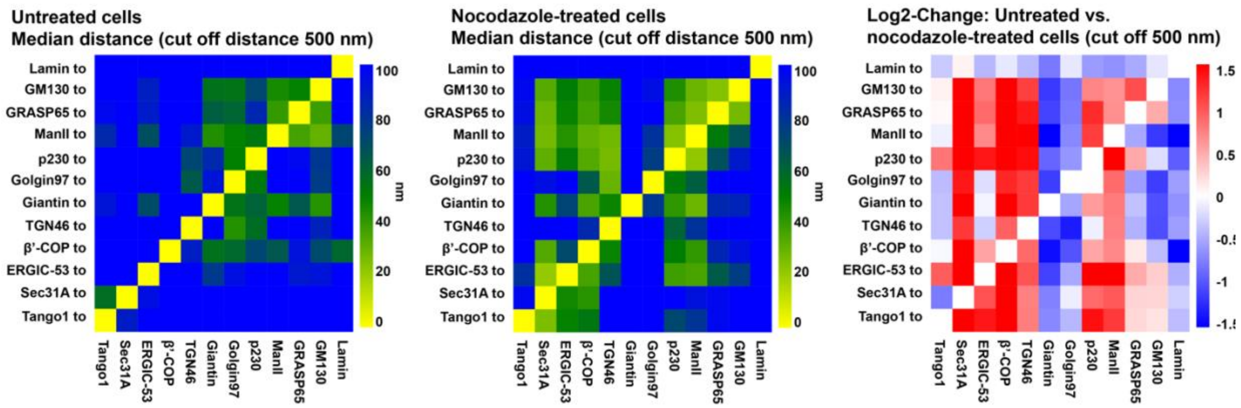

**Figure S25 – Distance Heatmaps and log2 change of proteins of the secretory pathway in untreated and nocodazole-treated HeLa cells shown in Figure 5.** The heatmap on the right represents the log2 change of distances (Distance Untreated cells/Distance Nocodazole-treated cells).

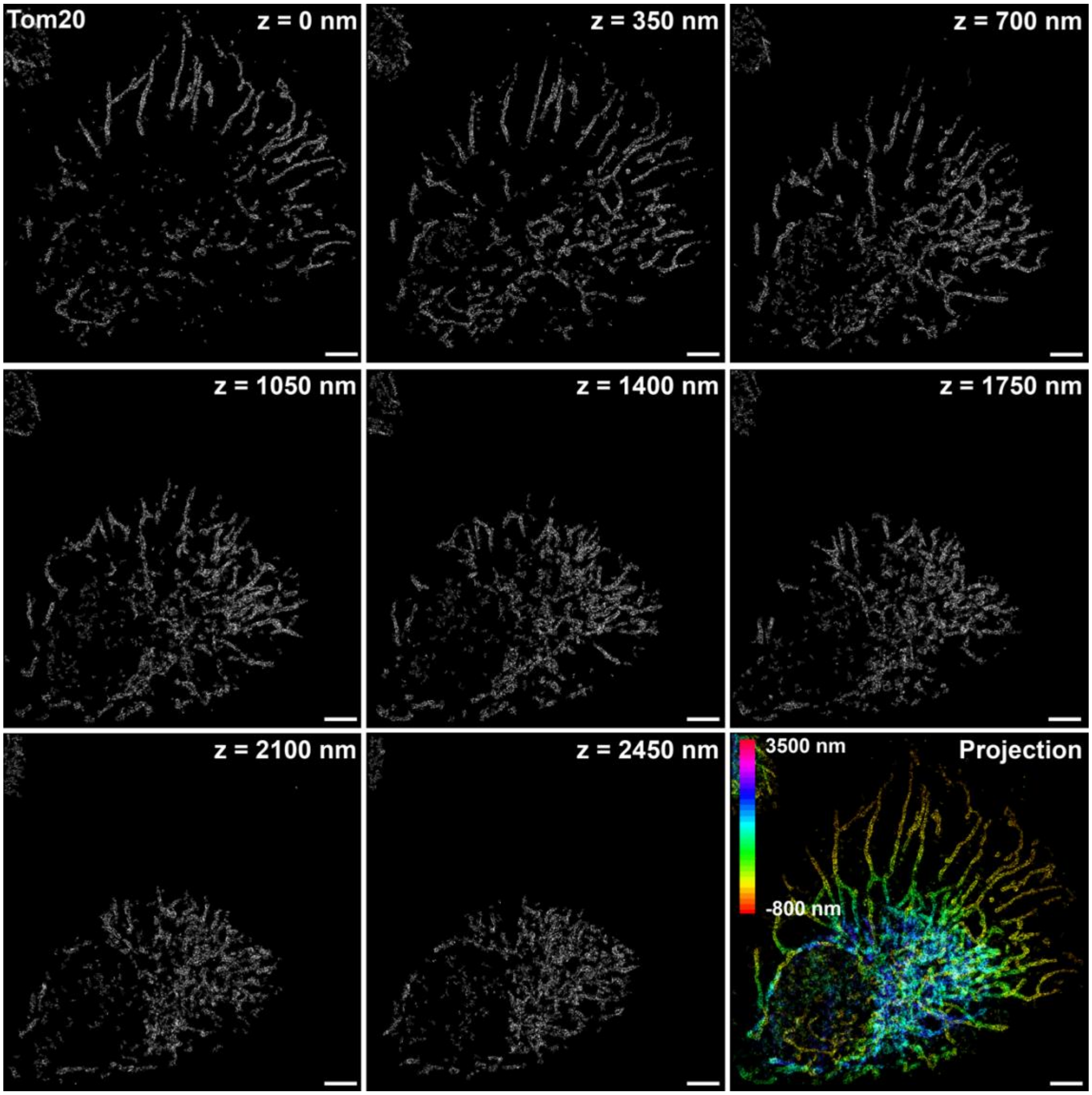

**Figure S26 – Individual optical sections of the Tom20 super-resolution image in Figure 6b.** The axial spacing between the optical sections is 350 nm. The last panel shows a z-projection with the color denoting the z-position of the localization events. Scale bar: 5 µm.

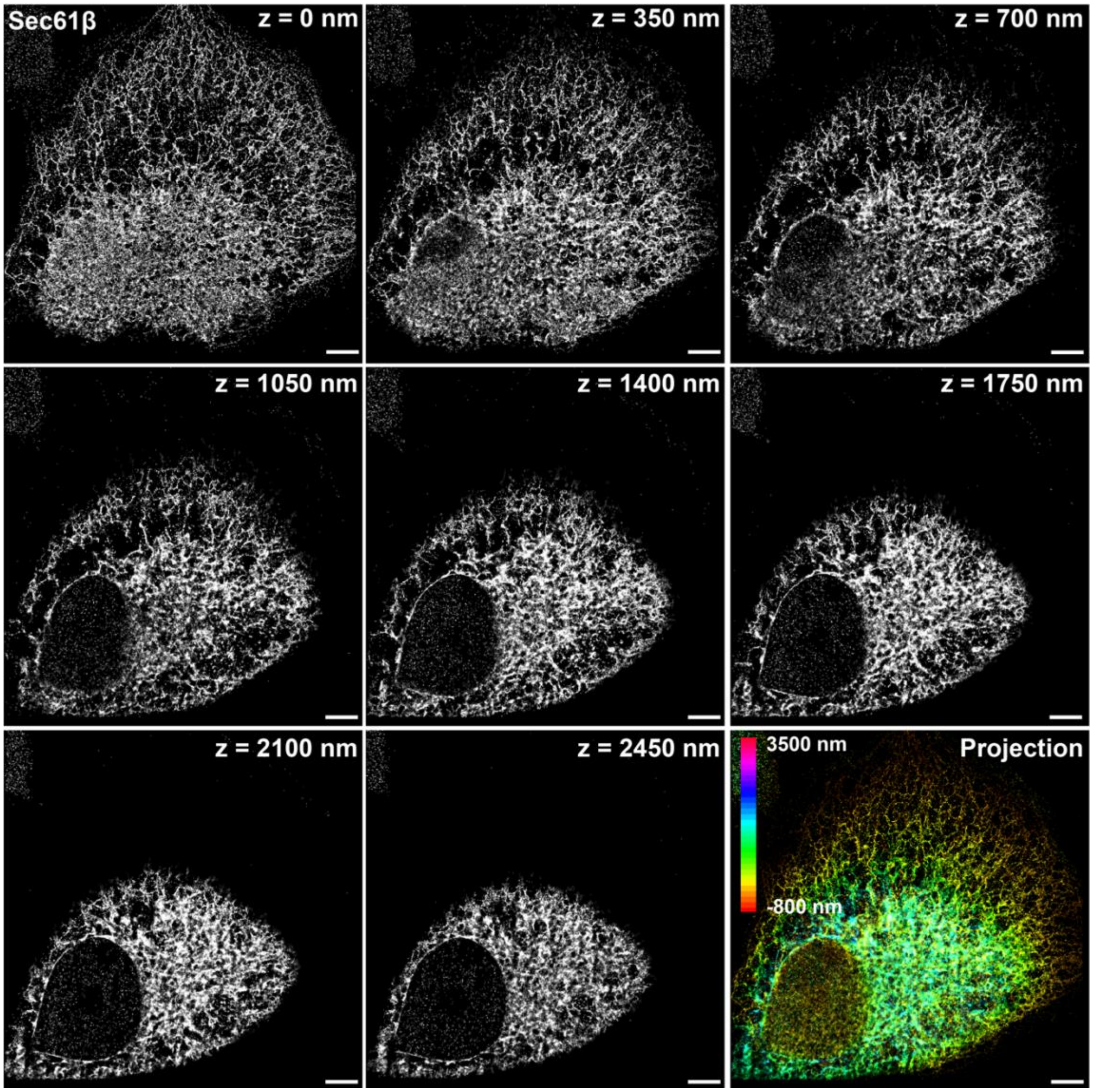

**Figure S27 – Individual optical sections of the Sec61β super-resolution image in Figure 6a.** The axial spacing between the optical sections is 350 nm. The last panel shows a z-projection with the color denoting the z-position of the localization events. Scale bar: 5 µm.

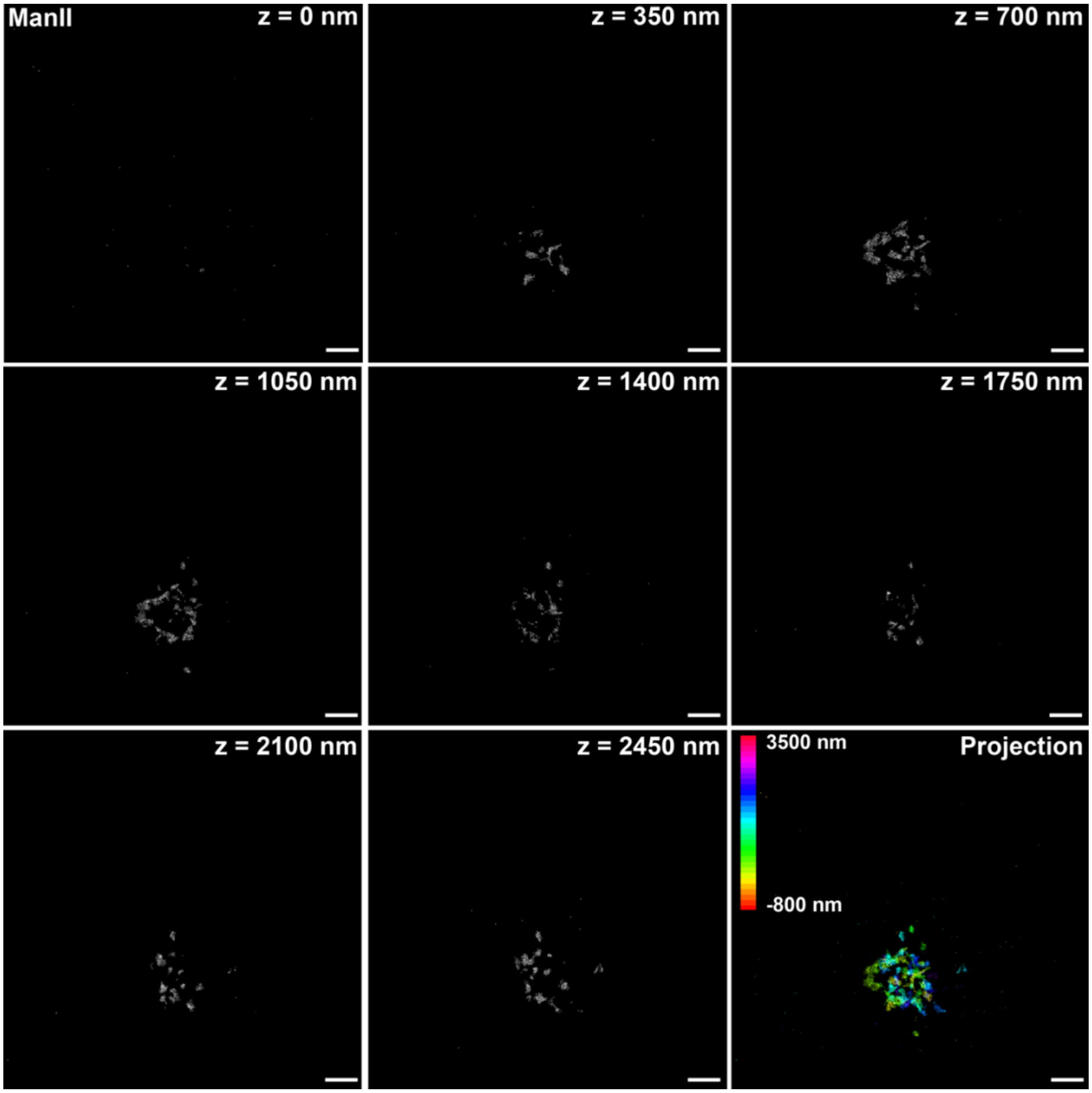

**Figure S28 – Individual optical sections of the ManII super-resolution image in Figure 6d.** The axial spacing between the optical sections is 350 nm. The last panel shows a z-projection with the color denoting the z-position of the localization events. Scale bar: 5 µm.

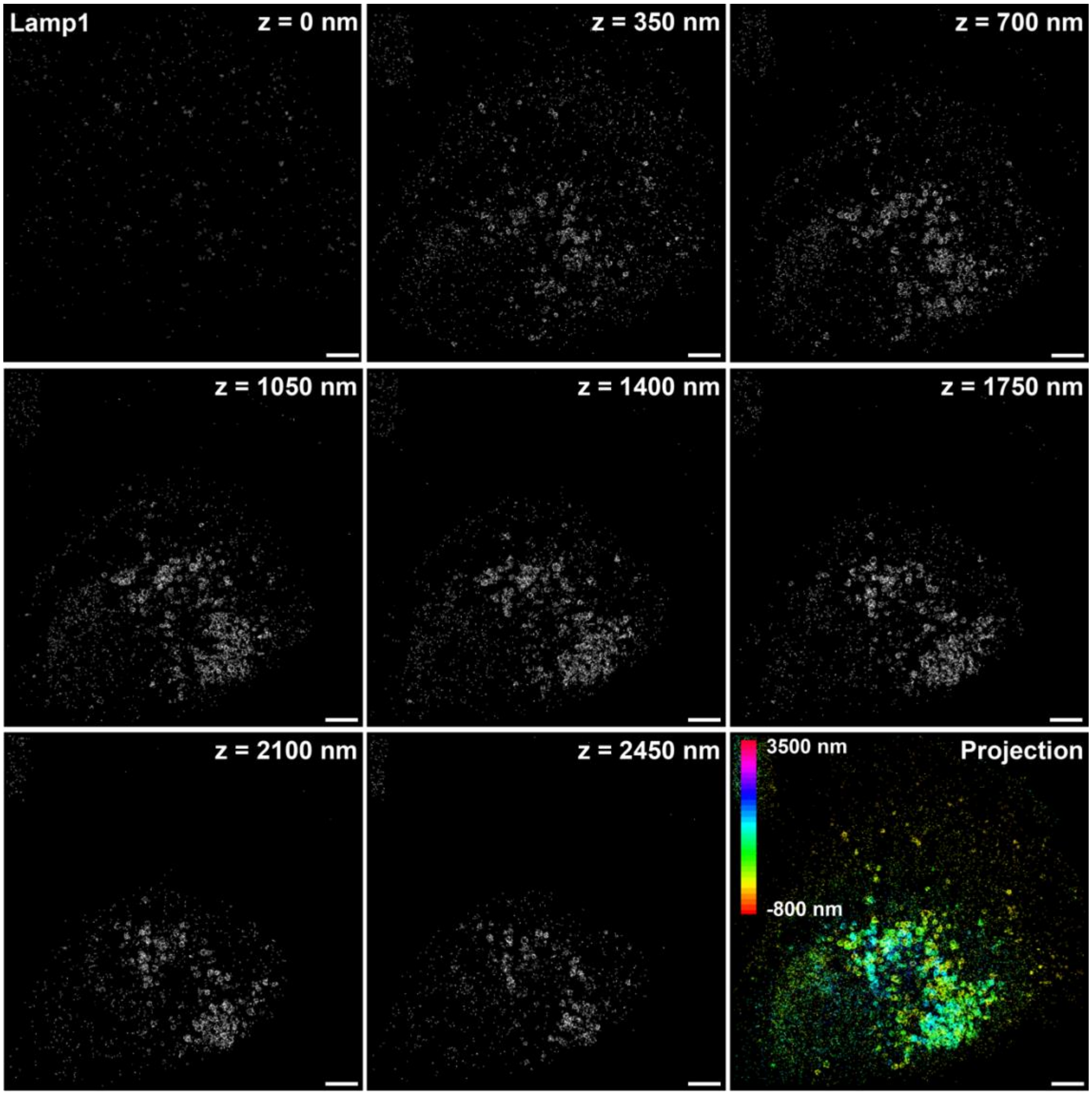

**Figure S29 – Individual optical sections of the Lamp1 super-resolution image in Figure 6a.** The axial spacing between the optical sections is 350 nm. The last panel shows a z-projection with the color denoting the z-position of the localization events. Scale bar: 5 µm.

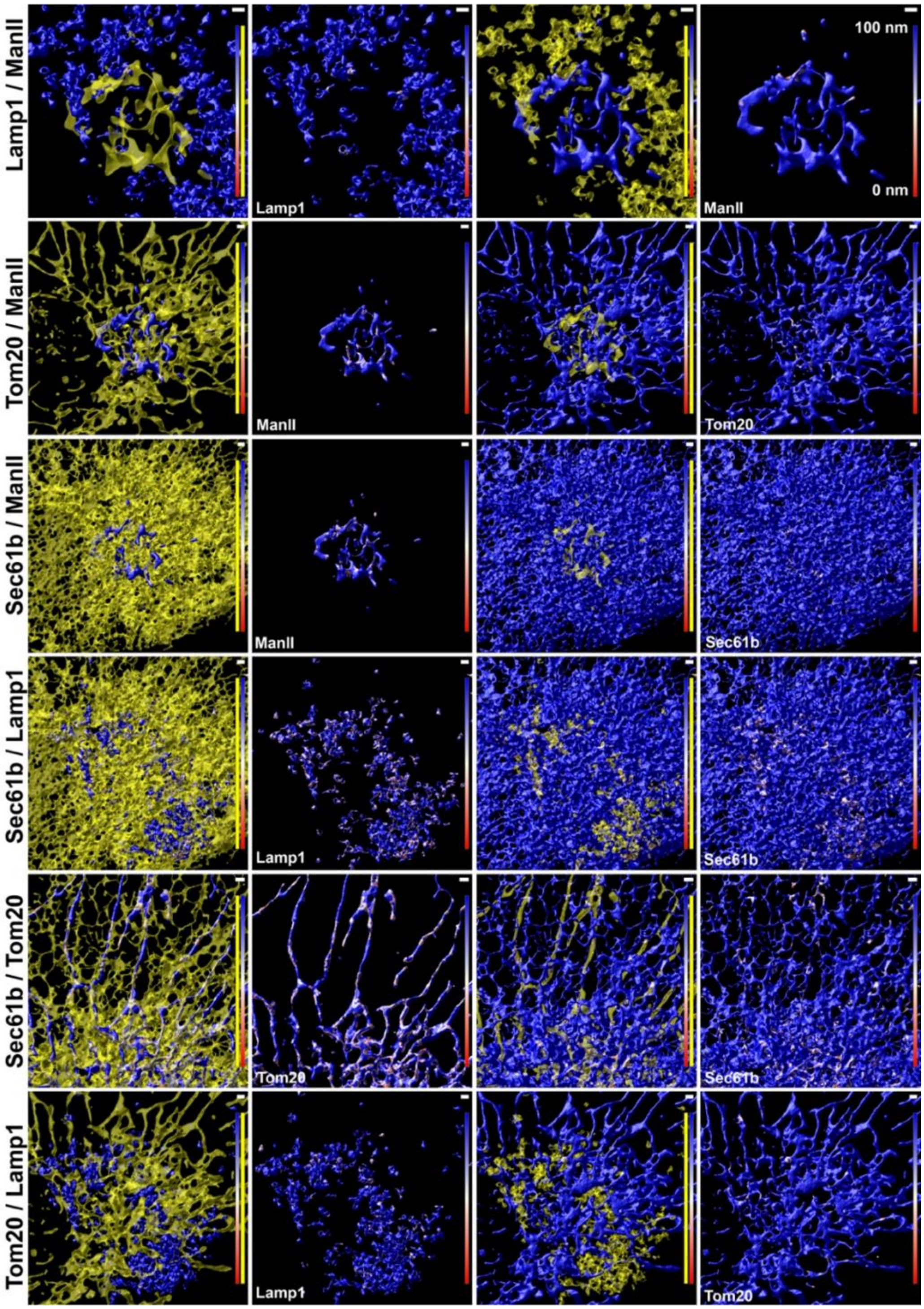

**Figure S30 – Contact sites between the ER (Sec61β), mitochondria (Tom20), lysosomes (Lamp1) and the Golgi complex (ManII-GFP).** The first and third columns display the pairs of organelles for which contact sites are calculated. The contact site is computed for the 'blue' organelle with respect to the 'yellow' organelle. The second and fourth columns present the corresponding contact site maps for the organelles shown in the first or third column. The colormap represents distances>100 nm in blue and distances<100 nm in white and red (see color bar). Scale bar: 1 µm.

| Imager name | Sequence | 5'-mod | 3'-mod |
|---|---|---|---|
| R2-6nt | TGGTGG | Cy3B | |
| P1-9nt | TAGATGTAT | Cy3B | |
| FP2 | AAGAAGTAAAGGGAG | Cy3B | BHQ2 |
| P3 | TAATGAAGA | | Cy3B |
| P5 | ATACATTGA | | Cy3B |
| PS3 | TCCTCCC | | Cy3B |
| R3 | GAGAGAG | | Cy3B |
| R4 | TGTGTGT | | Cy3B |

**Table S1 – Imager sequences.**

| Paper ID | Working ID | Sequence |
|---|---|---|
| A1 | A3 | TT TCTTCATTAGCG |
| A2 | A19 | TT ATAAAGTGTCCA |
| A3 | A5 | TT TCAATGTATGGC |
| A4 | A10 | TT ATAATGGATGGG |
| A5 | A27 | TT AAAAAGTTCGAG |
| A6 | A36 | TT ATAACAGAATCG |
| A7 | A39 | TT TTATGTTCTGCT |
| A8 | A15 | TT ATAGTGATTGGA |
| A9 | A38 | TT ATTTAGTGTAGC |
| A10 | A8 | TT ATGTTAATGGGT |
| A11 | A25 | TT ATAATCATGCTC |
| A12 | A20 | TT ATATGATCTCCG |
| P1 | P1 | TT ATA CAT CTA |
| P3 | P3 | TT TCT TCA TTA |
| P5 | P5 | TT TCA ATG TAT |
| PS3 | PS3 | AA GGGAGGA |
| 5xR3 | 5xR3 | TT TCCTCTCTCTCTCTCTC |
| 5xR4 | 5xR4 | TT CAACACACACACACACA |

**Table S1 – Docking-site sequences.**

| Paper ID | Working ID | Imager-DS | Spacer | Adapter-DS |
|---|---|---|---|---|
| A1-FP2 | A3-FP2 | CCTCGCTGAACCCCTTA | AA | CGCTAATGAA |
| A2-FP2 | A19-FP2 | CCTCGCTGAACCCCTTA | AA | TGGACACTTT |
| A3-FP2 | A5-FP2 | CCTCGCTGAACCCCTTA | AA | GCCATACATT |
| A4-FP2 | A10-FP2 | CCTCGCTGAACCCCTTA | AA | CCCATCCATT |
| A5-FP2 | A27-FP2 | CCTCGCTGAACCCCTTA | AA | CTCGAACTTT |
| A6-FP2 | A36-FP2 | CCTCGCTGAACCCCTTA | AA | CGATTCTGTT |
| A7-FP2 | A39-FP2 | CCTCGCTGAACCCCTTA | AA | AGCAGAACAT |
| A8-FP2 | A15-FP2 | CCTCGCTGAACCCCTTA | AA | TCCAATCACT |
| A9-FP2 | A38-FP2 | CCTCGCTGAACCCCTTA | AA | GCTACACTAA |
| A10-FP2 | A8-FP2 | CCTCGCTGAACCCCTTA | AA | ACCCATTAAC |
| A11-FP2 | A25-FP2 | CCTCGCTGAACCCCTTA | AA | GAGCATGATT |
| A12-FP2 | A20-FP2 | CCTCGCTGAACCCCTTA | AA | CGGAGATCAT |
| A1-P1 | A3-P1 | ATACATCTA | TT | CGCTAATGAA |
| A2-P1 | A19-P1 | ATACATCTA | TT | TGGACACTTT |
| A3-P1 | A5-P1 | ATACATCTA | TT | GCCATACATT |
| A4-P1 | A10-P1 | ATACATCTA | TT | CCCATCCATT |
| A5-P1 | A27-P1 | ATACATCTA | TT | CTCGAACTTT |
| A6-P1 | A36-P1 | ATACATCTA | TT | CGATTCTGTT |
| A7-P1 | A39-P1 | ATACATCTA | TT | AGCAGAACAT |
| A8-P1 | A15-P1 | ATACATCTA | TT | TCCAATCACT |
| A9-P1 | A38-P1 | ATACATCTA | TT | GCTACACTAA |
| A10-P1 | A8-P1 | ATACATCTA | TT | ACCCATTAAC |
| A11-P1 | A25-P1 | ATACATCTA | TT | GAGCATGATT |
| A12-P1 | A20-P1 | ATACATCTA | TT | CGGAGATCAT |
| A1-5xR2 | A3-5xR2 | ACCACCACCACCACCACCA | AA | CGCTAATGAA |
| A2-5xR2 | A19-5xR2 | ACCACCACCACCACCACCA | AA | TGGACACTTT |
| A3-5xR2 | A5-5xR2 | ACCACCACCACCACCACCA | AA | GCCATACATT |
| A4-5xR2 | A10-5xR2 | ACCACCACCACCACCACCA | AA | CCCATCCATT |
| A5-5xR2 | A27-5xR2 | ACCACCACCACCACCACCA | AA | CTCGAACTTT |
| A6-5xR2 | A36-5xR2 | ACCACCACCACCACCACCA | AA | CGATTCTGTT |
| A7-5xR2 | A39-5xR2 | ACCACCACCACCACCACCA | AA | AGCAGAACAT |
| A8-5xR2 | A15-5xR2 | ACCACCACCACCACCACCA | AA | TCCAATCACT |
| A9-5xR2 | A38-5xR2 | ACCACCACCACCACCACCA | AA | GCTACACTAA |
| A10-5xR2 | A8-5xR2 | ACCACCACCACCACCACCA | AA | ACCCATTAAC |
| A11-5xR2 | A25-5xR2 | ACCACCACCACCACCACCA | AA | GAGCATGATT |
| A12-5xR2 | A20-5xR2 | ACCACCACCACCACCACCA | AA | CGGAGATCAT |

**Table S3 – Adapter sequences.**

| Paper ID | Working ID | Association Rate (Adapter) *$10^6$ ($M^{-1}s^{-1}$) | Association Rate (Direct) *$10^6$ ($M^{-1}s^{-1}$) | Dissociation Rate (Adapter) (1/s) | Dissociation Rate (Direct) (1/s) |
|---|---|---|---|---|---|
| A1-FP2 | A3-FP2 | 0.68 | 0.8 | 12.5 | 11.1 |
| A2-FP2 | A19-FP2 | 0.53 | 0.57 | 7.14 | 8.33 |
| A3-FP2 | A5-FP2 | 0.55 | 0.8 | 8.33 | 9.1 |
| A4-FP2 | A10-FP2 | 0.59 | 0.51 | 7.69 | 11.1 |
| A5-FP2 | A27-FP2 | 0.53 | 0.49 | 7.69 | 10 |
| A6-FP2 | A36-FP2 | 0.45 | 0.46 | 6.67 | 12.5 |
| A7-FP2 | A39-FP2 | 0.35 | 0.4 | 5.55 | 5.88 |
| A8-FP2 | A15-FP2 | 0.55 | 0.79 | 8.33 | 7.14 |
| A9-FP2 | A38-FP2 | 0.4 | 0.6 | 7.69 | 10 |
| A10-FP2 | A8-FP2 | 0.45 | 0.62 | 9.1 | 8.33 |
| A11-FP2 | A25-FP2 | 0.4 | 0.7 | 5.55 | 9.1 |
| A12-FP2 | A20-FP2 | 0.4 | 0.57 | 6.49 | 10 |
| A1-P1 | A3-P1 | 1 | 1.09 | 2.04 | 1.3 |
| A2-P1 | A19-P1 | 1.4 | 1.42 | 1.72 | 1.23 |
| A3-P1 | A5-P1 | 1.58 | 1.39 | 1.79 | 1.23 |
| A4-P1 | A10-P1 | 1.62 | 1.46 | 1.75 | 1.25 |
| A5-P1 | A27-P1 | 1.45 | 1.36 | 2.08 | 1.41 |
| A6-P1 | A36-P1 | 1.38 | 1.45 | 2.44 | 1.56 |
| A7-P1 | A39-P1 | 1.75 | 1.35 | 2.27 | 1.69 |
| A8-P1 | A15-P1 | 2.02 | 2.2 | 1.37 | 1 |
| A9-P1 | A38-P1 | 1.42 | 1.38 | 1.61 | 1 |
| A10-P1 | A8-P1 | 1.96 | 1.82 | 1.35 | 1 |
| A11-P1 | A25-P1 | 1.31 | 1.37 | 1.64 | 1.02 |
| A12-P1 | A20-P1 | 1.61 | 1.48 | 1.79 | 1.19 |
| A1-5xR2 | A3-5xR2 | 58.33 | 53.94 | 4.76 | 5.88 |
| A2-5xR2 | A19-5xR2 | 40.95 | 46.3 | 6.25 | 6.67 |
| A3-5xR2 | A5-5xR2 | 40.3 | 42.95 | 6.67 | 6.67 |
| A4-5xR2 | A10-5xR2 | 39.58 | 34.57 | 4 | 5.26 |
| A5-5xR2 | A27-5xR2 | 49.52 | 45.75 | 5.88 | 7.14 |
| A6-5xR2 | A36-5xR2 | 52.25 | 51 | 5.55 | 7.14 |
| A7-5xR2 | A39-5xR2 | 60.34 | 55.67 | 5.26 | 5.26 |
| A8-5xR2 | A15-5xR2 | 40.09 | 42.48 | 5.88 | 7.14 |
| A9-5xR2 | A38-5xR2 | 38.17 | 47.12 | 7.14 | 7.14 |
| A10-5xR2 | A8-5xR2 | 35.36 | 39.59 | 7.69 | 7.69 |
| A11-5xR2 | A25-5xR2 | 42.53 | 41.28 | 7.14 | 7.14 |
| A12-5xR2 | A20-5xR2 | 47.39 | 42.64 | 5.55 | 6.67 |

**Table S4 – Erasers and their kinetics**

| Paper ID | Working ID | Complement to Adapter-DS sequence | Complement to Spacer | Partial complement to Imager-DS sequence |
|---|---|---|---|---|
| E1-FP2 | E3-FP2 | TTCATTAGCG | TT | TGG |
| E2-FP2 | E19-FP2 | AAAGTGTCCA | TT | TGG |
| E3-FP2 | E5-FP2 | AATGTATGGC | TT | TGG |
| E4-FP2 | E10-FP2 | AATGGATGGG | TT | TGG |
| E5-FP2 | E27-FP2 | AAAGTTCGAG | TT | TGG |
| E6-FP2 | E36-FP2 | AACAGAATCG | TT | TGG |
| E7-FP2 | E39-FP2 | ATGTTCTGCT | TT | TGG |
| E8-FP2 | E15-FP2 | AGTGATTGGA | TT | TGG |
| E9-FP2 | E38-FP2 | TTAGTGTAGC | TT | TGG |
| E10-FP2 | E8-FP2 | GTTAATGGGT | TT | TGG |
| E11-FP2 | E25-FP2 | AATCATGCTC | TT | TGG |
| E12-FP2 | E20-FP2 | ATGATCTCCG | TT | TGG |
| E1-P1 | E3-P1 | TTCATTAGCG | AA | TAG |
| E2-P1 | E19-P1 | AAAGTGTCCA | AA | TAG |
| E3-P1 | E5-P1 | AATGTATGGC | AA | TAG |
| E4-P1 | E10-P1 | AATGGATGGG | AA | TAG |
| E5-P1 | E27-P1 | AAAGTTCGAG | AA | TAG |
| E6-P1 | E36-P1 | AACAGAATCG | AA | TAG |
| E7-P1 | E39-P1 | ATGTTCTGCT | AA | TAG |
| E8-P1 | E15-P1 | AGTGATTGGA | AA | TAG |
| E9-P1 | E38-P1 | TTAGTGTAGC | AA | TAG |
| E10-P1 | E8-P1 | GTTAATGGGT | AA | TAG |
| E11-P1 | E25-P1 | AATCATGCTC | AA | TAG |
| E12-P1 | E20-P1 | ATGATCTCCG | AA | TAG |
| E1-R2 | E3-R2 | TTCATTAGCG | TT | TAA |
| E2-R2 | E19-R2 | AAAGTGTCCA | TT | TAA |
| E3-R2 | E5-R2 | AATGTATGGC | TT | TAA |
| E4-R2 | E10-R2 | AATGGATGGG | TT | TAA |
| E5-R2 | E27-R2 | AAAGTTCGAG | TT | TAA |
| E6-R2 | E36-R2 | AACAGAATCG | TT | TAA |
| E7-R2 | E39-R2 | ATGTTCTGCT | TT | TAA |
| E8-R2 | E15-R2 | AGTGATTGGA | TT | TAA |
| E9-R2 | E38-R2 | TTAGTGTAGC | TT | TAA |
| E10-R2 | E8-R2 | GTTAATGGGT | TT | TAA |
| E11-R2 | E25-R2 | AATCATGCTC | TT | TAA |
| E12-R2 | E20-R2 | ATGATCTCCG | TT | TAA |

**Table S5 – Eraser sequences.**

```
TTCCCTTCCTTTCTCGCCACGTTCGCCGGCTTTCCCCGTCAAGCTCTAAATCGGGGGCTCCCTTTAGGGTTCCGATTTAGTGCTTTACGGCACCTCGACCC
CAAAAAACTTGATTTGGGTGATGGTTCACGTAGTGGGCCATCGCCCTGATAGACGGTTTTTCGCCCTTTGACGTTGGAGTCCACGTTCTTTAATAGTGGAC
TCTTGTTCCAAACTGGAACAACACTCAACCCTATCTCGGGCTATTCTTTTGATTTATAAGGGATTTTGCCGATTTCGGAACCACCATCAAACAGGATTTTC
GCCTGCTGGGGCAAACCAGCGTGGACCGCTTGCTGCAACTCTCTCAGGGCCAGGCGGTGAAGGGCAATCAGCTGTTGCCCGTCTCACTGGTGAAAAGAAAA
ACCACCCTGGCGCCCAATACGCAAACCGCCTCTCCCCGCGCGTTGGCCGATTCATTAATGCAGCTGGCACGACAGGTTTCCCGACTGGAAAGCGGGCAGTG
AGCGCAACGCAATTAATGTGAGTTAGCTCACTCATTAGGCACCCCAGGCTTTACACTTTATGCTTCCGGCTCGTATGTTGTGTGGAATTGTGAGCGGATAA
CAATTTCACACAGGAAACAGCTATGACCATGATTACGAATTCGAGCTCGGTACCCGGGGATCCTCTAGAGTCGACCTGCAGGCATGCAAGCTTGGCACTGG
CCGTCGTTTTACAACGTCGTGACTGGGAAAACCCTGGCGTTACCCAACTTAATCGCCTTGCAGCACATCCCCCTTTCGCCAGCTGGCGTAATAGCGAAGAG
GCCCGCACCGATCGCCCTTCCCAACAGTTGCGCAGCCTGAATGGCGAATGGCGCTTTGCCTGGTTTCCGGCACCAGAAGCGGTGCCGGAAAGCTGGCTGGA
GTGCGATCTTCCTGAGGCCGATACTGTCGTCGTCCCCTCAAACTGGCAGATGCACGGTTACGATGCGCCCATCTACACCAACGTGACCTATCCCATTACGG
TCAATCCGCCGTTTGTTCCCACGGAGAATCCGACGGGTTGTTACTCGCTCACATTTAATGTTGATGAAAGCTGGCTACAGGAAGGCCAGACGCGAATTATT
TTTGATGGCGTTCCTATTGGTTAAAAAATGAGCTGATTTAACAAAAATTTAATGCGAATTTTAACAAAATATTAACGTTTACAATTTAAATATTTGCTTAT
ACAATCTTCCTGTTTTTGGGGCTTTTCTGATTATCAACCGGGGTACATATGATTGACATGCTAGTTTTACGATTACCGTTCATCGATTCTCTTGTTTGCTC
CAGACTCTCAGGCAATGACCTGATAGCCTTTGTAGATCTCTCAAAAATAGCTACCCTCTCCGGCATTAATTTATCAGCTAGAACGGTTGAATATCATATTG
ATGGTGATTTGACTGTCTCCGGCCTTTCTCACCCTTTTGAATCTTTACCTACACATTACTCAGGCATTGCATTTAAAATATATGAGGGTTCTAAAAATTTT
TATCCTTGCGTTGAAATAAAGGCTTCTCCCGCAAAAGTATTACAGGGTCATAATGTTTTTGGTACAACCGATTTAGCTTTATGCTCTGAGGCTTTATTGCT
TAATTTTGCTAATTCTTTGCCTTGCCTGTATGATTTATTGGATGTTAATGCTACTACTATTAGTAGAATTGATGCCACCTTTTCAGCTCGCGCCCCAAATG
AAAATATAGCTAAACAGGTTATTGACCATTTGCGAAATGTATCTAATGGTCAAACTAAATCTACTCGTTCGCAGAATTGGGAATCAACTGTTATATGGAAT
GAAACTTCCAGACACCGTACTTTAGTTGCATATTTAAAACATGTTGAGCTACAGCATTATATTCAGCAATTAAGCTCTAAGCCATCCGCAAAAATGACCTC
TTATCAAAAGGAGCAATTAAAGGTACTCTCTAATCCTGACCTGTTGGAGTTTGCTTCCGGTCTGGTTCGCTTTGAAGCTCGAATTAAAACGCGATATTTGA
AGTCTTTCGGGCTTCCTCTTAATCTTTTTGATGCAATCCGCTTTGCTTCTGACTATAATAGTCAGGGTAAAGACCTGATTTTTGATTTATGGTCATTCTCG
TTTTCTGAACTGTTTAAAGCATTTGAGGGGGATTCAATGAATATTTATGACGATTCCGCAGTATTGGACGCTATCCAGTCTAAACATTTTACTATTACCCC
CTCTGGCAAAACTTCTTTTGCAAAAGCCTCTCGCTATTTTGGTTTTTATCGTCGTCTGGTAAACGAGGGTTATGATAGTGTTGCTCTTACTATGCCTCGTA
ATTCCTTTTGGCGTTATGTATCTGCATTAGTTGAATGTGGTATTCCTAAATCTCAACTGATGAATCTTTCTACCTGTAATAATGTTGTTCCGTTAGTTCGT
TTTATTAACGTAGATTTTTCTTCCCAACGTCCTGACTGGTATAATGAGCCAGTTCTTAAAATCGCATAAGGTAATTCACAATGATTAAAGTTGAAATTAAA
CCATCTCAAGCCCAATTTACTACTCGTTCTGGTGTTTCTCGTCAGGGCAAGCCTTATTCACTGAATGAGCAGCTTTGTTACGTTGATTTGGGTAATGAATA
TCCGGTTCTTGTCAAGATTACTCTTGATGAAGGTCAGCCAGCCTATGCGCCTGGTCTGTACACCGTTCATCTGTCCTCTTTCAAAGTTGGTCAGTTCGGTT
CCCTTATGATTGACCGTCTGCGCCTCGTTCCGGCTAAGTAACATGGAGCAGGTCGCGGATTTCGACACAATTTATCAGGCGATGATACAAATCTCCGTTGT
ACTTTGTTTCGCGCTTGGTATAATCGCTGGGGGTCAAAGATGAGTGTTTTAGTGTATTCTTTTGCCTCTTTCGTTTTAGGTTGGTGCCTTCGTAGTGGCAT
TACGTATTTTACCCGTTTAATGGAAACTTCCTCATGAAAAAGTCTTTAGTCCTCAAAGCCTCTGTAGCCGTTGCTACCCTCGTTCCGATGCTGTCTTTCGC
TGCTGAGGGTGACGATCCCGCAAAAGCGGCCTTTAACTCCCTGCAAGCCTCAGCGACCGAATATATCGGTTATGCGTGGGCGATGGTTGTTGTCATTGTCG
GCGCAACTATCGGTATCAAGCTGTTTAAGAAATTCACCTCGAAAGCAAGCTGATAAACCGATACAATTAAAGGCTCCTTTTGGAGCCTTTTTTTTGGAGAT
TTTCAACGTGAAAAAATTATTATTCGCAATTCCTTTAGTTGTTCCTTTCTATTCTCACTCCGCTGAAACTGTTGAAAGTTGTTTAGCAAAATCCCATACAG
AAAATTCATTTACTAACGTCTGGAAAGACGACAAAACTTTAGATCGTTACGCTAACTATGAGGGCTGTCTGTGGAATGCTACAGGCGTTGTAGTTTGTACT
GGTGACGAAACTCAGTGTTACGGTACATGGGTTCCTATTGGGCTTGCTATCCCTGAAAATGAGGGTGGTGGCTCTGAGGGTGGCGGTTCTGAGGGTGGCGG
TTCTGAGGGTGGCGGTACTAAACCTCCTGAGTACGGTGATACACCTATTCCGGGCTATACTTATATCAACCCTCTCGACGGCACTTATCCGCCTGGTACTG
AGCAAAACCCCGCTAATCCTAATCCTTCTCTTGAGGAGTCTCAGCCTCTTAATACTTTCATGTTTCAGAATAATAGGTTCCGAAATAGGCAGGGGGCATTA
ACTGTTTATACGGGCACTGTTACTCAAGGCACTGACCCCGTTAAAACTTATTACCAGTACACTCCTGTATCATCAAAAGCCATGTATGACGCTTACTGGAA
CGGTAAATTCAGAGACTGCGCTTTCCATTCTGGCTTTAATGAGGATTTATTTGTTTGTGAATATCAAGGCCAATCGTCTGACCTGCCTCAACCTCCTGTCA
ATGCTGGCGGCGGCTCTGGTGGTGGTTCTGGTGGCGGCTCTGAGGGTGGTGGCTCTGAGGGTGGCGGTTCTGAGGGTGGCGGCTCTGAGGGAGGCGGTTCC
GGTGGTGGCTCTGGTTCCGGTGATTTTGATTATGAAAAGATGGCAAACGCTAATAAGGGGGCTATGACCGAAAATGCCGATGAAAACGCGCTACAGTCTGA
CGCTAAAGGCAAACTTGATTCTGTCGCTACTGATTACGGTGCTGCTATCGATGGTTTCATTGGTGACGTTTCCGGCCTTGCTAATGGTAATGGTGCTACTG
GTGATTTTGCTGGCTCTAATTCCCAAATGGCTCAAGTCGGTGACGGTGATAATTCACCTTTAATGAATAATTTCCGTCAATATTTACCTTCCCTCCCTCAA
TCGGTTGAATGTCGCCCTTTTGTCTTTGGCGCTGGTAAACCATATGAATTTTCTATTGATTGTGACAAAATAAACTTATTCCGTGGTGTCTTTGCGTTTCT
TTTATATGTTGCCACCTTTATGTATGTATTTTCTACGTTTGCTAACATACTGCGTAATAAGGAGTCTTAATCATGCCAGTTCTTTTGGGTATTCCGTTATT
ATTGCGTTTCCTCGGTTTCCTTCTGGTAACTTTGTTCGGCTATCTGCTTACTTTTCTTAAAAAGGGCTTCGGTAAGATAGCTATTGCTATTTCATTGTTTC
TTGCTCTTATTATTGGGCTTAACTCAATTCTTGTGGGTTATCTCTCTGATATTAGCGCTCAATTACCCTCTGACTTTGTTCAGGGTGTTCAGTTAATTCTC
CCGTCTAATGCGCTTCCCTGTTTTTATGTTATTCTCTCTGTAAAGGCTGCTATTTTCATTTTTGACGTTAAACAAAAAATCGTTTCTTATTTGGATTGGGA
TAAATAATATGGCTGTTTATTTTGTAACTGGCAAATTAGGCTCTGGAAAGACGCTCGTTAGCGTTGGTAAGATTCAGGATAAAATTGTAGCTGGGTGCAAA
ATAGCAACTAATCTTGATTTAAGGCTTCAAAACCTCCCGCAAGTCGGGAGGTTCGCTAAAACGCCTCGCGTTCTTAGAATACCGGATAAGCCTTCTATATC
TGATTTGCTTGCTATTGGGCGCGGTAATGATTCCTACGATGAAAATAAAAACGGCTTGCTTGTTCTCGATGAGTGCGGTACTTGGTTTAATACCCGTTCTT
GGAATGATAAGGAAAGACAGCCGATTATTGATTGGTTTCTACATGCTCGTAAATTAGGATGGGATATTATTTTTCTTGTTCAGGACTTATCTATTGTTGAT
AAACAGGCGCGTTCTGCATTAGCTGAACATGTTGTTTATTGTCGTCGTCTGGACAGAATTACTTTACCTTTTGTCGGTACTTTATATTCTCTTATTACTGG
CTCGAAAATGCCTCTGCCTAAATTACATGTTGGCGTTGTTAAATATGGCGATTCTCAATTAAGCCCTACTGTTGAGCGTTGGCTTTATACTGGTAAGAATT
TGTATAACGCATATGATACTAAACAGGCTTTTTCTAGTAATTATGATTCCGGTGTTTATTCTTATTTAACGCCTTATTTATCACACGGTCGGTATTTCAAA
CCATTAAATTTAGGTCAGAAGATGAAATTAACTAAAATATATTTGAAAAAGTTTTCTCGCGTTCTTTGTCTTGCGATTGGATTTGCATCAGCATTTACATA
TAGTTATATAACCCAACCTAAGCCGGAGGTTAAAAAGGTAGTCTCTCAGACCTATGATTTTGATAAATTCACTATTGACTCTTCTCAGCGTCTTAATCTAA
GCTATCGCTATGTTTTCAAGGATTCTAAGGGAAAATTAATTAATAGCGACGATTTACAGAAGCAAGGTTATTCACTCACATATATTGATTTATGTACTGTT
TCCATTAAAAAAGGTAATTCAAATGAAATTGTTAAATGTAATTAATTTTGTTTTCTTGATGTTTGTTTCATCATCTTCTTTTGCTCAGGTAATTGAAATGA
ATAATTCGCCTCTGCGCGATTTTGTAACTTGGTATTCAAAGCAATCAGGCGAATCCGTTATTGTTTCTCCCGATGTAAAAGGTACTGTTACTGTATATTCA
TCTGACGTTAAACCTGAAAATCTACGCAATTTCTTTATTTCTGTTTTACGTGCAAATAATTTTGATATGGTAGGTTCTAACCCTTCCATTATTCAGAAGTA
TAATCCAAACAATCAGGATTATATTGATGAATTGCCATCATCTGATAATCAGGAATATGATGATAATTCCGCTCCTTCTGGTGGTTTCTTTGTTCCGCAAA
ATGATAATGTTACTCAAACTTTTAAAATTAATAACGTTCGGGCAAAGGATTTAATACGAGTTGTCGAATTGTTTGTAAAGTCTAATACTTCTAAATCCTCA
AATGTATTATCTATTGACGGCTCTAATCTATTAGTTGTTAGTGCTCCTAAAGATATTTTAGATAACCTTCCTCAATTCCTTTCAACTGTTGATTTGCCAAC
TGACCAGATATTGATTGAGGGTTTGATATTTGAGGTTCAGCAAGGTGATGCTTTAGATTTTTCATTTGCTGCTGGCTCTCAGCGTGGCACTGTTGCAGGCG
GTGTTAATACTGACCGCCTCACCTCTGTTTTATCTTCTGCTGGTGGTTCGTTCGGTATTTTTAATGGCGATGTTTTAGGGCTATCAGTTCGCGCATTAAAG
ACTAATAGCCATTCAAAAATATTGTCTGTGCCACGTATTCTTACGCTTTCAGGTCAGAAGGGTTCTATCTCTGTTGGCCAGAATGTCCCTTTTATTACTGG
TCGTGTGACTGGTGAATCTGCCAATGTAAATAATCCATTTCAGACGATTGAGCGTCAAAATGTAGGTATTTCCATGAGCGTTTTTCCTGTTGCAATGGCT
GGCGGTAATATTGTTCTGGATATTACCAGCAAGGCCGATAGTTTG
```

**Table S6 – M13mp18 Scaffold sequence.**

**Table S7 – DNA origami staples for Frames. See Excel sheet.**

**Table S8 – DNA origami staples for 20 nm grids. See Excel sheet.**

**Table S9 – DNA origami staples for 10 nm grids. See Excel sheet.**

**Table S10 – DNA origami staples for the Y letter. See Excel sheet.**

**Table S11 – DNA origami staples for the A letter. See Excel sheet.**

**Table S12 – DNA origami staples for the L letter. See Excel sheet.**

**Table S13 – DNA origami staples for the E letter. See Excel sheet.**

| Oligo Name | Sequence |
|---|---|
| **Staple1 - BIOTIN** | ATTAAGTTTACCGAGCTCGAATTCGGGAAACCTGTCGTGC |
| **Staple2 - BIOTIN** | ATAAGGGAACCGGATATTCATTACGTCAGGACGTTGGGAA |
| **Staple3 - BIOTIN** | GCGATCGGCAATTCCACACAACAGGTGCCTAATGAGTG |
| **Staple4 - BIOTIN** | TTGTGTCGTGACGAGAAACACCAAATTTCAACTTTAAT |
| **Staple5 - BIOTIN** | ATTCATTTTTGTTTGGATTATACTAAGAAACCACCAGAAG |
| **Staple6 - BIOTIN** | CACCCTCAGAAACCATCGATAGCATTGAGCCATTTGGGAA |
| **Staple7 - BIOTIN** | AACAATAACGTAAAACAGAAATAAAAATCCTTTGCCCGAA |
| **Staple8 - BIOTIN** | AGCCACCACTGTAGCGCGTTTTCAAGGGAGGGAAGGTAAA |

**Table S14 – Biotin staple sequence.**

**Table S15 – Table for experimental settings for DNA origami experiments. See Excel sheet.**

| # | Protein | Host | Vendor | Cat. Number | Docking site |
|---|---|---|---|---|---|
| 1 | GM130 | Rabbit | Proteintech | 11308-1-AP | N/A |
| 2 | GOLGB1 Giantin | Rabbit | Sigma | HPA011555 | N/A |
| 3 | GOLGA1_1 Golgin-97 | Rabbit | Atlas Antibodies | HPA044329 | N/A |
| 4 | TGN46 | Rabbit | Proteintech | 10598-1-AP | N/A |
| 5 | ERGIC-3 | Rabbit | Abcam | ab129179 | N/A |
| 6 | LMAN1 ERGIC-53 | Mouse | Invitrogen | MA5-25345 | N/A |
| 7 | GRASP65 | Rabbit | Abcam | ab174834 | N/A |
| 8 | Anti-MIA3 (Tango1) | Rabbit | Sigma | HPA055922 | N/A |
| 9 | p230 | Mouse | BD Biosciences | 611280 | N/A |
| 10 | COPI (CMIA10) | Mouse | Rothman Lab | Custom | N/A |
| 11 | COPII (Sec31A) | Mouse | BD | 612350 | N/A |
| 12 | LaminB1 | Rabbit | Abcam | ab16048 | N/A |
| 13 | GM130 | Mouse | BD Biosciences | 610822 | N/A |
| 14 | GRASP55 | Rabbit | Proteintech | 10598-1-AP | N/A |
| 15 | HADHA | Mouse | Abcam | ab110302 | N/A |
| 16 | dsDNA | Mouse | abcam | ab3519 | N/A |
| 17 | RPA40 | Mouse | Santa Cruz | sc-374443 | N/A |
| 18 | NPM1 | Mouse | Novus Bio | NB600-1030 | N/A |
| 19 | Tom20 | Rabbit | Santa Cruz | sc-11415 | N/A |
| 20 | Alpha-Tubulin | Mouse | Sigma | T5168 | N/A |
| 21 | Glutamylated-tubulin | Rabbit | Millipore | AB3201 | OyOlink-A8 |
| 22 | Inpp5e | Rabbit | Proteintech | 17797-1-AP | OyOlink-A10 |
| 23 | Arl13b | Rabbit | Proteintech | 17711-1-AP | OyOlink-A15 |
| 24 | Ift88 | Rabbit | Proteintech | 13967-1-AP | OyOlink-A27 |
| 25 | CEP164 | Rabbit | Proteintech | 22227-1-AP | OyOlink-A36 |
| 26 | RPGRIP1L | Rabbit | Proteintech | 55160-1-AP | OyOlink-A38 |
| 27 | Septin2 | Rabbit | Abcam | ab187654 | OyOlink-A39 |
| 28 | Acetylated-tubulin | Mouse | Sigma | T6793 | A19 |
| 29 | Anti-Lamp1 | Rabbit | Cell Signaling Technology | 9091 | N/A |
| 30 | Anti-mCherry | Mouse | GeneTex | GT844 | N/A |
| 31 | Anti-mCherry | Mouse | GeneTex | GT857 | N/A |
| 32 | GFP-Nanobody | N/A | Massive Photonics | Custom | A3 |
| 33 | Anti-Mouse Nanobody | N/A | Massive Photonics | Custom | A10 |
| 34 | Anti-Mouse Nanobody | N/A | Massive Photonics | Custom | A19 |
| 35 | Anti-Mouse Nanobody | N/A | Massive Photonics | Custom | A25 |
| 36 | Anti-Mouse Nanobody | N/A | Massive Photonics | Custom | A27 |
| 37 | Anti-Mouse Nanobody | N/A | Massive Photonics | Custom | A36 |
| 38 | Anti-Mouse Antibody | N/A | Custom | Custom | A5 |
| 39 | Anti-Rabbit Nanobody | N/A | Massive Photonics | Custom | A8 |
| 40 | Anti-Rabbit Nanobody | N/A | Massive Photonics | Custom | A15 |
| 41 | Anti-Rabbit Nanobody | N/A | Massive Photonics | Custom | A20 |
| 42 | Anti-Rabbit Nanobody | N/A | Massive Photonics | Custom | A38 |
| 43 | Anti-Rabbit Nanobody | N/A | Massive Photonics | Custom | A39 |
| 44 | Anti-Rabbit Nanobody | N/A | Massive Photonics | Custom | 5xR3 |
| 45 | Anti-Rabbit Nanobody | N/A | Massive Photonics | Custom | 5xR4 |
| 46 | Anti-Rabbit Antibody | N/A | Custom | Custom | A5 |
| 47 | Anti-Rabbit Antibody | N/A | Custom | Custom | A19 |

**Table S16 – Antibodies and Nanobodies.**

**Table S17 – Table for experimental settings for cell experiments. See Excel sheet.**

| Figure | Number of imaging rounds | $c_{imager}$ | $c_{adapter}$ | Number of camera frames | Camera exposure time/ readout bandwidth | Laser power/ intensity | Further information |
|---|---|---|---|---|---|---|---|
| 1b | 1 | 500 pM | 20 nM | 10,000 | 100 ms/ 200 MHz | 50 mW/ ~1.6 kW/cm$^2$ | Table S15 |
| 1c | 1 | 5 nM - 10 nM (P1, P3, PS3) | 50 nM | 5,000 - 10,000 | 100 ms/ 200 MHz | 27 mW-50 mW/ ~0.8 kW/cm$^2$ ~1.6 kW/cm | Table S15 |
| 1d & S3 (each data point) | 1 | 5 nM - 10 nM (R2,PS3, P1) | 0 – 100 nM | 5,000- 30,000 | 33 ms-50 ms / 200 MHz | 15 mW-50 mW/ ~0.5 kW/cm$^2$ - 1.6 kW/cm$^2$ | Table S15 |
| 1e & S10-14 | 4 | 200 pM (R2) | 20 nM | 10,000 | 100 ms/ 200 MHz | 90 mW/ ~2.9 kW/cm$^2$ | Table S15 |
| 2 & S17 & Movie S1 & S2 | 2 | 10 nM (R2) | 20 nM | 5,000 | 200 ms/ 200 MHz | 5 mW/ 0.15 kW/cm$^2$ | Table S17 |
| 3 & Figure S21 | 10 | 100 pM – 4 nM (R2) | 20 nM | 40,000 | 20 ms/ 540 MHz | 70 – 100 mW/ 2.3 – 3.3 kW/cm$^2$ | Table S17 |
| 4 & S22 | 9 | 500 pM – 2 nM (R2) | 20 nM | 10,000 per plane | 20 ms/ 540 MHz | 50 mW 1.6 kW/cm$^2$ | Table S17 |
| 5 & S23 & S24 | 13 | 100 pM – 2 nM (R2) | 20 nM | 30,000 | 20 ms/ 540 MHz | 94 mW -188 mW 3.1 – 6.1 kW/cm$^2$ | Table S17 |
| 6 & S26-29 | 4 | 10 nM – 40 nM (FP2) | 50 nM | 20,000 – 40,000 per plane | 10 ms/ 540 MHz | 125 mW/ 4.1 kW/cm$^2$ | Table S17 |
| S4 – S9 | 4 | 5 nM (R2, PS3, P1, P5) | 20 nM & 50 nM | 5,000 - 30,000 | 33 ms – 100 ms 200Mhz/540 MHz | 25 – 90 mW 0.8 – 1.6 kW/cm$^2$ | Table S15 |
| S16 | 6 | 5 nM (R2, PS3, P5) | 20 nM | 5,000 - 30,000 | 33 ms – 50 ms 200Mhz/540 MHz | 15 – 50 mW 0.5 – 1.6 kW/cm$^2$ | Table S15 |
| S15 | 4 | 400 pM – 2 nM (P1, R2, FP2) | 20 nM | 20,000 | 30 ms/540 MHz | 80 mW/2.6 kW/cm$^2$ | Table S17 |
| S18 -S20 | 4 | 1 nM – 20 nM | 20 nM – 50 nM | 30,000 | 33 ms/540 MHz | 80 mW/2.6 kW/cm$^2$ | Table S17 |

**Table S18 – Summary of experimental conditions for imaging experiments.**
